# Multi-omics of the gut microbial ecosystem in the immunotherapy resistance in microsatellite instability-high gastrointestinal cancer patients

**DOI:** 10.1101/2023.03.07.531467

**Authors:** Siyuan Cheng, Zihan Han, Xiaochen Yin, Die Dai, Fang Li, Xiaotian Zhang, Ming Lu, Zhihao Lu, Xicheng Wang, Jun Zhou, Jian Li, Xiaohuan Guo, Panwei Song, Chuanzhao Qiu, Wei Shen, Qi Zhang, Ning Zhu, Xi Wang, Yan Tan, Lin Shen, Yan Kou, Zhi Peng

## Abstract

Despite the encouraging efficacy of anti-PD-1/PD-L1 immunotherapy in microsatellite instability-high/deficient mismatch repair (MSI-H/dMMR) advanced gastrointestinal cancer, many patients exhibit primary or acquired resistance. Using multi-omics approaches, we interrogated gut microbiome, blood metabolome and cytokines/chemokines of MSI-H/dMMR advanced gastrointestinal cancer patients (N=77) and identified a number of microbes (e.g. Alistipes putredinis) and metabolites (e.g. arginine and SCFA) highly associated with primary resistance. Fecal microbiota transplantation of patients’ stool replicated the clinical responsiveness as well as certain molecular signatures. Based on the clinical microbiome data, we developed a predictive machine learning model for primary resistance and achieved accuracy at 0.83 on an external validation set. Furthermore, several microbes were pinpointed which gradually changed during the process of acquired resistance. In summary, our study demonstrated the essential role of gut microbiome in drug resistance, and this could be utilized as a preventative diagnosis tool as well as therapeutic targets in the future.

## 1. Introduction

The mismatch repair-deficient (dMMR) subtype accounts for 15 - 22% of patients with gastrointestinal (GI) cancers^1–3^ and ~ 5% of metastatic/recurrent GI cancers^4–6^. The dMMR subtype patients are unable to recognize and repair certain spontaneous mutations, resulting in a quite high tumor mutation burden and microsatellite instability-high (MSI-H) status, therefore are more likely to benefit from anti-PD-1/PD-L1 immunotherapy. According to existing evidences, MSI status is one of the most effective biomarker of cancer immunotherapy. However, MSI-H/dMMR GI cancer patients present highly heterogeneous responses to immunotherapy and roughly 30% of MSI-H patients exhibit primary resistance to immunotherapy^7^. Moreover, ~ 17% of MSI-H patients present acquired resistance following a 2-year treatment, and the proportion usually increase with the prolonged treatment^7^. Considering the rather consistent molecular patterns within MSI-H/dMMR subtype, we believe understanding other factors which impact efficacy heterogeneity is of great importance to enhance immunotherapyresponse in dMMR/MSI-H patients.

In our previous work, we demonstrated that gut microbiome composition could predict the efficacy of anti-PD-1 immunotherapy in GI cancer patients^8^, and researches on other tumor types also implicate the role of gut microbiome in modulating host response of various immunotherapies^9–12^. Metabolites of the gut microbiome, such as short-chain fatty acids (SCFAs)^13–16^ and inosine^17^ are shown to influence host immunity – synergistically modulating anti-tumor effects. Two independent investigations demonstrated that responder-derived fecal microbiota transplantation (FMT) could restore anti-PD-1 responses in PD-1-refactory melanoma patients^18,19^. Despite of these encouraging findings, microbiota signatures related to efficacy vary from different cohorts^20^, and this phenomenon may be attributed to the heterogeneity between tumor types and even within the same cancer type. To certain aspect, MSI-H/dMMR patients present highly consistent molecular characteristics, and thus research in this subtype is more likely to find microbiota biomarkers that can be generalized and applied in multiple tumor types. Moreover, previous studies mainly focus on primary resistance, which further restricts their clinical translation, considering a rather large proportion of patients usually develop acquired resistance to immunotherapy treatment.

We present here the findings of our investigation into the gut microbiome profiles of 77 advanced MSI-H/dMMR GI cancer patients receiving anti-PD-1/PD-L1 therapy. To examine and identify interplay between gut microbiota, their metabolic products, and host immune responses germane to primary and acquired drug resistance, we applied integrated multi-omics analysis to paired blood and fecal samples at baseline and during treatments, then we further developed an effective predictive machine-learning model based on microbiome signatures. A summary of the microbial, metabolomic, and immunologic signatures identified in this study as relating to primary and/or acquired anti-PD-1/PD-L1 resistance ensues.

## 2. STAR Methods

### 2.1 Cohort recruitment

Advanced gastrointestinal cancer patients with MSI-H/dMMR were enrolled for participation at Beijing Cancer Hospital. All participants were at stage III/IV and undergoing treatment with anti PD-1/PD-L1, in some instances combined with anti-CTLA-4 immunotherapy. Response to treatment was evaluated every 6 weeks and confirmed no less than 4 weeks from the date first recorded. In accordance with guidelines put forth in Response Evaluation Criteria in Solid Tumors version 1.1 (RECIST 1.1), patients in this study deemed to be either responders or non-responders. Patients classified as responders were further divided into acquired resistance [AR; where the best response is stable disease (SD) or partial response (PR) and progressive disease (PD) is acquired after 6 months] or long responders [LR; where SD or partial response lasts more than 1year *sans* PD at the time of enlistment]. Patients were classified as non-responders if the best response was PD or SD or PR with acquired PD in 6 months. Clinical laboratory examination and antibiotic (ATB) usage within 30 days were documented for all patients. This study was approved by the Ethics Committee at Beijing Cancer Hospital. Informed content was obtained from all patients enrolled for the collection of clinical information and samples, and all tests and procedures were conducted in accordance with the Declaration of Helsinki.

### 2.2 Fecal microbiome sample preparation and data analysis

Stool samples were collected from patients at various timepoints throughout their immunotherapy regimen as previously described^8^ (**Figure.1**). Briefly, DNA was extracted using QIAamp PowerFecal DNA Kit (catalog No. 12830-50, Qiagen), followed by library construction and sequencing on an Illumina NovoSeq 6000 platform (Novo Gene). Raw reads were quality controlled via KneadData (version 0.6.1), which integrated several QC tools [*e.g*., FastQC (https://www.bioinformatics.babraham.ac.uk/projects/fastqc/) and Trimmomatic^21^. After trimming low-quality bases from reads and omitting reads less than 100 bp in length, bowtie2^22^ mapped filtered reads to the human genome (hg19), and host contaminants were removed. Taxonomic profiling was accomplished using MetaPhlAn2 with default parameters^23^, and functional analyses were achieved using HUMAnN2 alongside the UniRef90 reference database^24^.

**Figure 1.**
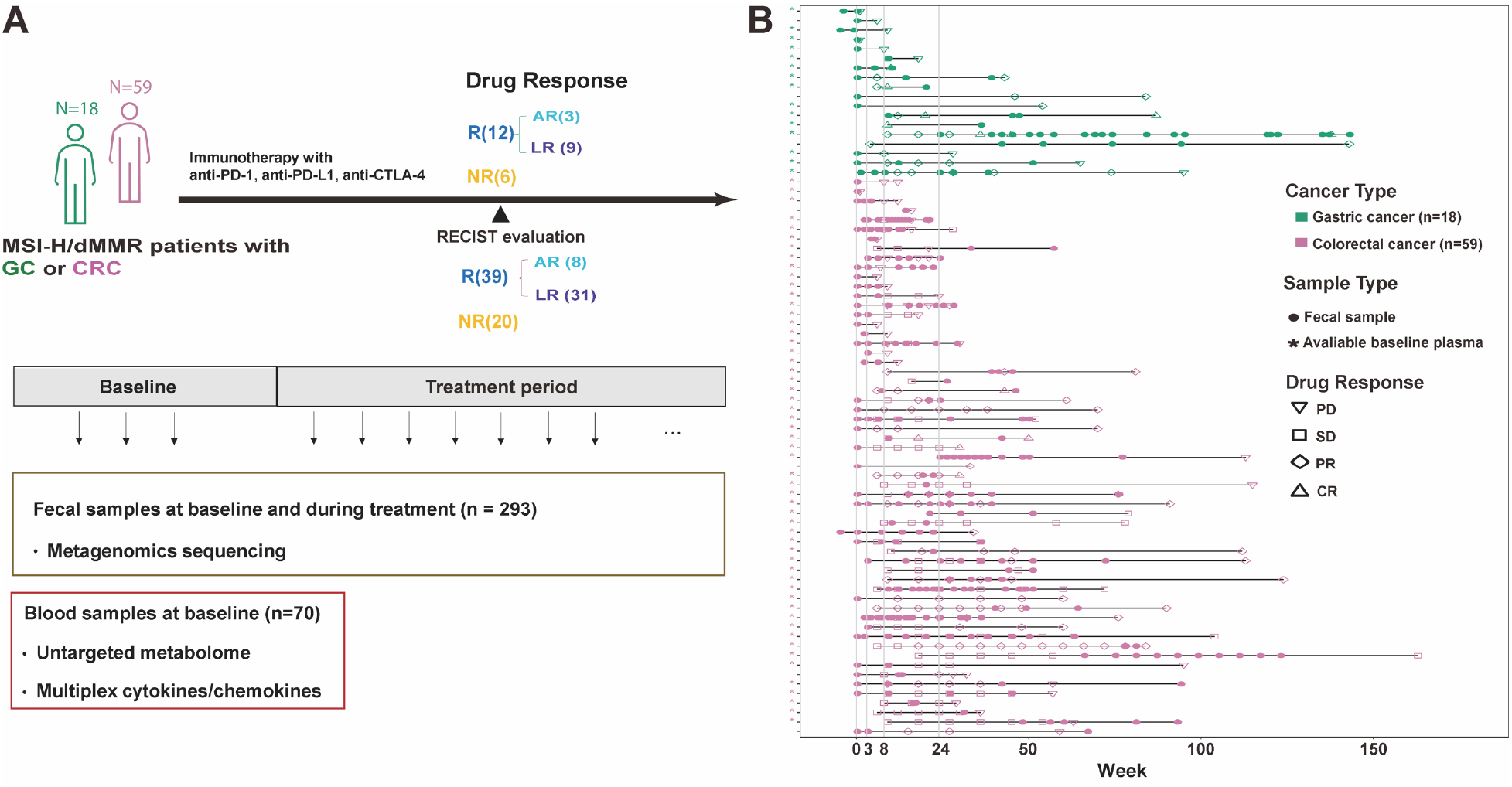
Study design and clinical sample collection. **(A)** 18 gastric (GC) and 59 colorectal cancer (CRC) patients with MSI-H/dMMR were recruited. Patients were classified as being responders (R) and non-responders (NR). Acquired resistance (AR) patients and long responders (LR) were further differentiated in responders. **(B)** In total, 293 stool and 70 blood plasma samples were collected. CR, complete response; PD, progressive disease; PR, partial response; SD, stable disease.

Once feature tables were generated (taxa, gene, pathway), alpha community diversity was calculated in accordance with Shannon indices while beta diversity was deduced based on Bray-Curtis distances. Permanova and beta dispersion analyses were applied to identify confounding clinical variables, such as age and gender. Features observed in a differential manner were then selected with MaAsLin2^25^ using previously identified confounding factors as fixed effects. When analyzing multiple samples from the same patient, “subject” was also treated as a random effect. All statistical analyses and plotting were performed in Rstudio (R version 3.6.3).

### 2.3 Blood metabolome sample preparation and data analysis

Baseline blood samples were drawn from 70 patients, and plasma was prepared by centrifugation at 3,000 rpm for 10 min at 4 °C and stored at –80 °C. In preparation for metabolomics analysis, plasma samples were thawed and centrifuged at 14,000 x g for 20 min in a cold room (4–8 °C), and supernatants were transferred to sterile 1.5 ml microfuge tubes. A volume of 400 μL methanol (pre-chilled to −80 °C) was added to each 100 μL supernatant. The final 80% (v/v) methanol solution was shaken and incubated at −80 °C for 2h. Following centrifugation (14,000 x g, 10 min, 4°C), supernatants were transferred into autosampler vials and subjected to direct UHPLC-MS analysis.

Ultimate 3000 UHPLC (Dionex) coupled with Orbitrap mass spectrometry (Thermo Fisher) was used to perform LC separation. In the negative mode, the BEH C18 column (2.1×100 mm, Waters) was applied for analysis at 0.25 mL/min. Mobile phase A was crafted by mixing 1L of HPLC-grade water containing 0.3953 g of Ammonium bicarbonate (pH ~8). Mobile phase B was 100% ACN. A gradient was established as follows: 0~3min, 1% B; 10~17 min, 99% B; 17.1~20.0 min, 1% B. In positive mode, samples were passed over a Waters BEH amide column (100 *2.1 mm, 1.7 μm) heated to 40°C using a gradient of 1% B to 99% B over 12.5 min (solvent A: 95% ACN + 5% H2O + 10mM NH4FA; solvent B: 50% IPA + 50% ACN +10mM NH4FA). Data with masses ranging from m/z 80-1200 to m/z 70-1050 were acquired at both the positive and negative ion modes with data dependent MSMS acquisition. The full scan and fragment spectra were collected at a resolution of 60,000 and 15,000, respectively. Detailed mass spectrometer parameters were as follows: spray voltage at 2.8 kV for negative and 3.2 kV for positive; capillary temperature set at 320°C; heater temperature set at 300°C; sheath gas flow rate set to 35; auxiliary gas flow rate set to 10. Metabolite identification was achieved based on Tracefinder (Thermofisher, CA) searches using a home-built database.

We applied the standard 80% rule to minimize the effect of missing values and retain only metabolites detectable in 80% or more patients in at least one group. QC samples were prepared by pooling equivalent aliquots of plasma from each of the samples and injected intermittently into the analytical process to monitor the stability of the method. Variables presenting relative standard deviations (RSD) greater than 30% in quality control (QC) samples were removed, replaced with one-half of the minimum value found in that data set. Total area normalization and logarithmic transformation was performed to stabilize variance across the intensity range. Mean-normalized data from positive and negative ion modes were combined for downstream analysis. PCA, Permanova, and beta dispersion analyses were performed on normalized feature tables to identify potential confounding factors, and pathway enrichment analyses were achieved using the MetaboAnalystR package^26^.

### 2.4 Blood SCFA quantification

Short chain fatty acids were analyzed with a 6500plus QTrap mass spectrometer (AB SCIEX, USA) coupled to an ACQUITY UPLC H-Class system (Waters, USA). An ACQUITY UPLC BEH C18 column (2.1×100mm, 1.7μm, Waters) was employed with mobile phase A: 100% water and mobile phase B: 100% ACN over a linear gradient of 0-1 min, 2% B; 1-10 min, 50% B;10-12 min, 98% B; 12-15 min, 2%B. Flow rate was 0.3 mL/min. The column chamber and sample tray were held at 45°C and 10°C, respectively. Data were acquired in multiple reaction monitormode, and ion transitions were optimized using chemical standards. Nebulizer (Gas1), heater (Gas2), and curtain gas pressures were set at 50, 50, and 35 psi, respectively. Source voltage was - 4500 V for negative ion mode, and optimal probe temperature was determined to be 550°C. SCIEX OS 1.6 software was used to identify metabolites and integrate peaks.

### 2.5 Blood cytokine/chemokine quantification

Concentrations of IL-1β, MIP-1α, PlGF, TARC, VEGF, IL-17A, IL-6, IP-10, MCP-1, IL-16, MIP-1β, IFN-γ, IL-5, TNF-α, IL-13, IL-15, MCP-4, CRP, GM-CSF, ICAM-1, IL-12p70, IL-2, IL-7, IL-8-HA-, Tie-2, VCAM-1, VEGF-D, Flt-1, bFGF, Eotaxin, Eotaxin-3, IL-10, IL-12-IL-23p40, IL-1α, IL-4, IL-8, MDC, SAA, TNF-β, and VEGF-C were measured from patient blood samples collected at baseline using V-PLEX Plus Kits (Catalog NO: K15198G, K15049G, K15190G, K15050G, K15047G; Meso Scale Diagnostics, Rockville, MD, USA), per manufacturer’s instructions. Two technical replicates were measured for each sample and average values were recorded. Assays were performed at AliveX Biotech (Shanghai, China).

### 2.6 Animal experiment and multi-omics data generation

This animal study was approved by the Animal Care and Use Committee at Beijing cancer hospital (Approval Number EAEC2021-02). We constructed the MC38 cell line transplant tumor model of C57BL/6J mice (female, aged 6 weeks, purchased from the Beijing Huafukang Bioscience http://www.hfkbio.com/). Animals were kept in the SPF condition. Antibiotics (ANVM, 1 g/L Ampicillin, 1 g/L neomycin, 1 g/L metronidazole, 0.5 g/L vancomycin) were administered through drinking water at 2 weeks prior to stool gavage (FMT). MC38 murine colon adenocarcinoma cells were cultured in DMEM medium (Gibco) with 10% FBS and 1% penicillin-streptomycin (Thermo Fisher Scientific) at 37°C with 5% CO_2_. MC38 cells were subcutaneously injected into the right flank of mice at a dosage of 5 x 10^5^ one week after the initiation of FMT. Once the tumor reached ~ 50 mm^3^, mice were intraperitoneally injected with 100ug PD-1 blockade (RMP1-14, BioXcell) every 3 days for 12 days (4 injections). Tumor volume was calculated by measuring length (a) and width (b) via vernier caliper and applying the equation v = ab^2^/2. To prepare the stool for gavage, one gram of stool sample was suspended in 5 mL of phosphate-buffered saline (PBS). Mice were orally gavaged with 200 μL of processed donor stool samples or PBS (as the negative control) every three days. Fecal material was obtained from 3 responders and 3 non-responders, with each gavaged to 3 mice and total 21 mice used in this study. All mice were sacrificed 4 weeks after tumor transplantation and tumor tissues were immediately fixed in formalin. Multiplex panel immunofluorescence (mIHC) staining of tumor tissues was performed by Crown Bioscience Inc. Briefly, FFPE blocks were sectioned with a manual rotary microtome, 4 μm thickness/section. The following antibodies were used with the Bond RX autostainer: anti-CD3 (dilution 1:500, Invitrogen), anti-CD8 (dilution 1:400, Cell Signaling), anti-CD4 (dilution 1:100, Cell Signaling), anti-Foxp3 (dilution 1:400, Cell Signaling), and anti-IFN-γ (dilution 1:100, Abcam). All stained sections were scanned with Vectra® multiplexed imaging systems at 20x magnification. High resolution imagery of whole sections was generated and subjected to quantification analysis with HALOTM software. Statistical analyses of tumor growth and immune cell infiltration was performed using one-way ANOVA followed by Tukey’s multiple comparisons test (p < 0.05). The method of blood metabolomics is the same as human metabolome analysis. Applying Spearman’s rank correlation coefficient, networks were constructed between blood metabolites and tumor mIHC results. RNA sequencing and data analysis were completed by Novogene Co., Ltd (Beijing, China). Briefly, total amounts and integrity of RNA were assessed using the RNA Nano 6000 Assay Kit of the Bioanalyzer 2100 system (Agilent Technologies, CA, USA) followed by library construction and sequenced by the Illumina NovaSeq 6000 (150bp PE). We used clusterProfiler R package (3.8.1) to test the statistical enrichment of differential expression genes in KEGG pathways (http://www.genome.jp/kegg/).

### 2.7 Machine learning model construction and validation

To identify potential responders at baseline level, we trained a XGBoost classifier using baseline metagenomic data of MSI-high responders and non-responders. Relative abundance of species and functional enrichment of pathways were preprocessed through log-transformation after adding a constant of 1 to avoid negative infinity. Cancer types were transformed into dummy variables to improve model performance. The whole dataset was split into train and test set using ‘train_test_split’ of ‘scikit-learn’ package with parameter ‘test_size=15’. As our training sample size is relatively small, we augmented the data by copying the original one and then adding a constant to each feature (diagnosis excluded) of individual sample. The constant is determined by random selection from a normal distribution where mean equals to 0 and standard deviation equals to half of the standard deviation of the sample, after which the absolute value is applied. We evaluated the performance of XGBoost classifier using 5-fold cross-validation. Hyperparameters of the model were automatically estimated using ‘hpsklearn’ python package with parameters ‘algo=tpe.suggest, max_evals=100, trial_timeout=300’. Measurement of feature importance was performed by using ‘SHAP’ package. An external in-house metagenomics dataset of 29 stool samples was also applied to test the model performance.

## 3. Results

### 3.1 Advanced MSI-H/dMMR GI cancer cohort

A total of 98 advanced gastrointestinal cancer patients with MSI-H/dMMR were consented and enrolled at the Beijing Cancer Hospital from February 2018 to October 2020, including 30 gastric cancer (GC) and 68 colorectal cancer (CRC) patients. Fifteen patients were excluded because anti-PD-1/PD-L1 was combined with chemo- or targeted therapies (n=12) or used as adjuvant therapy (n=3). Six patients did not provide sufficient fecal samples for metagenomic sequencing therefore were removed from further analysis (**Figure S1**). The current cohort included 77 patients, 18 of gastric cancer and 59 of colorectal cancer. All patients were identified as MSI-H or dMMR and thus received anti-PD-1/PD-L1 immunotherapy alone or combined with anti-CTLA4 blockade. We defined a patient as a responder (R) if the patient achieved an objective response (CR/PR/SD) lasting at least 6 months upon treatment start, or a non-responder (NR) (PD observed within 6 months of treatment start). Response rates (~ 34% primary resistance) across our unique cohort of 77 MSI-H/dMMR GI cancer patients were comparable to other cohorts^7^. No significant association was observed between clinical benefits and metadata such as age, gender, and BMI (**Table 1**). Baseline is defined as pre-treatment or no more than 3 weeks after immunotherapy treatment. Total 293 fecal samples were collected at baseline and along the treatment period (**Figure 1**). Meanwhile, baseline blood samples (plasma) from a subset of patients (15 GC, 55 CRC) were also collected for metabolome and cytokine/chemokine panel measurement.

**Table 1.**
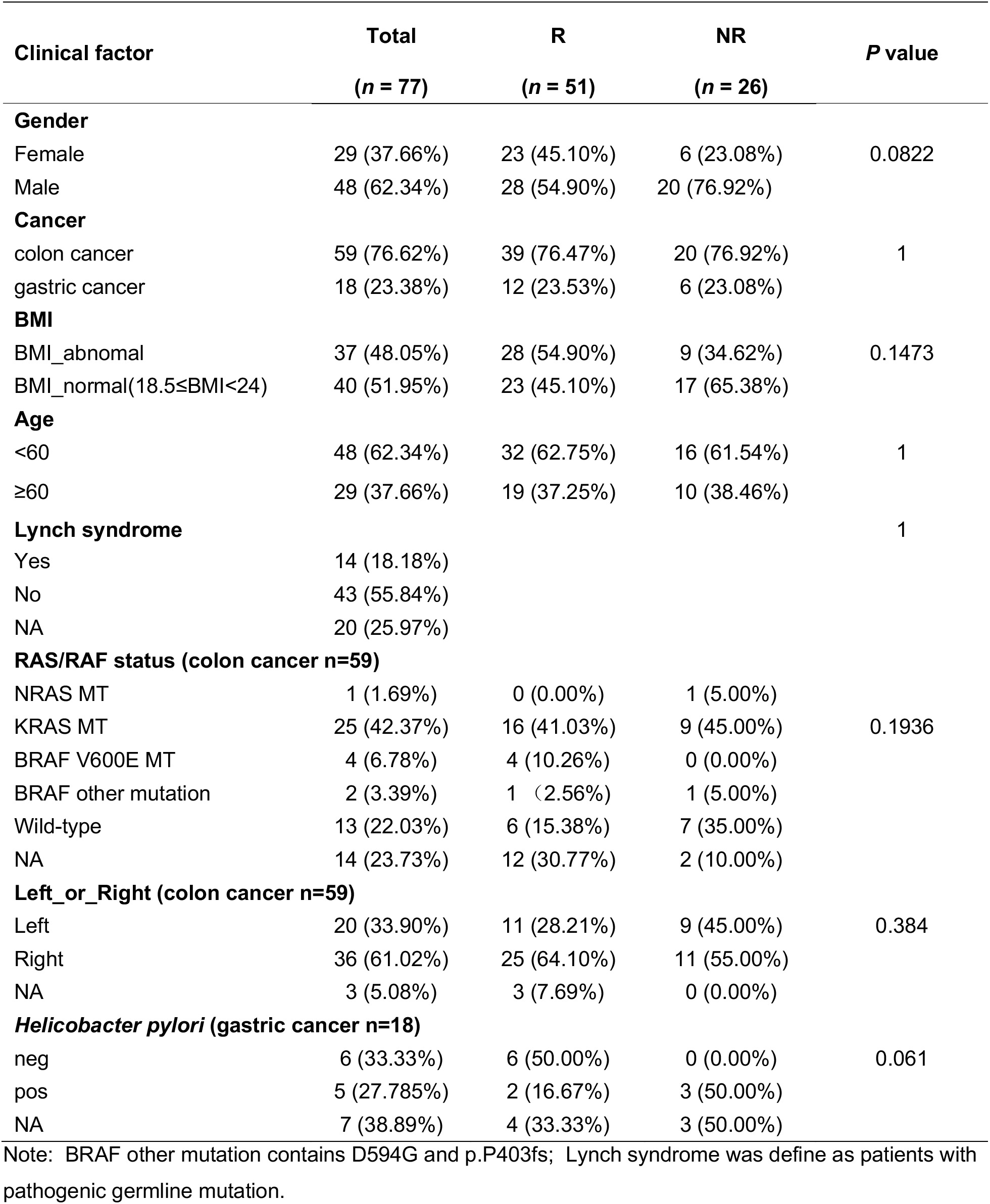
Cohort characteristics.

### 3.2 Biosignatures of patients with primary resistance to immunotherapy

To explore the global signature associated with primary resistance to immunotherapy, we performed multi-omics analysis of the baseline samples collected from the 77 patients. We first analyzed the metagenomics data collected from the fecal samples. Consistent with the findings of our previous study involving 74 GI cancer patients^8^, no significant differences were observed in alpha diversity between responders and non-responders based on Shannon indices (**Figure S2A**). Beta diversity depicting the overall gut microbial structure showed no significant difference between responders and non-responders (PERMANOVA *P*=0.5) (**Figure 2A**). We further explored the significantly altered species between R and NR patients and found that *Bacteroides caccae, Porphyromonadaceae, Parabacteroides, Acidaminococcaceae, Alistipes finegoldii, Paraprevotella clara, Bacteroides massiliensis* and *Alistipes putredinis* were enriched in responders, while *Veillonella parvula, Veillonella atypica, Peptostreptococcaceae, Streptococcus thermophilus* and *Micrococcaceae* were enriched in the non-responder group (**Figure 2B & Table S1**).

**Figure 2.**
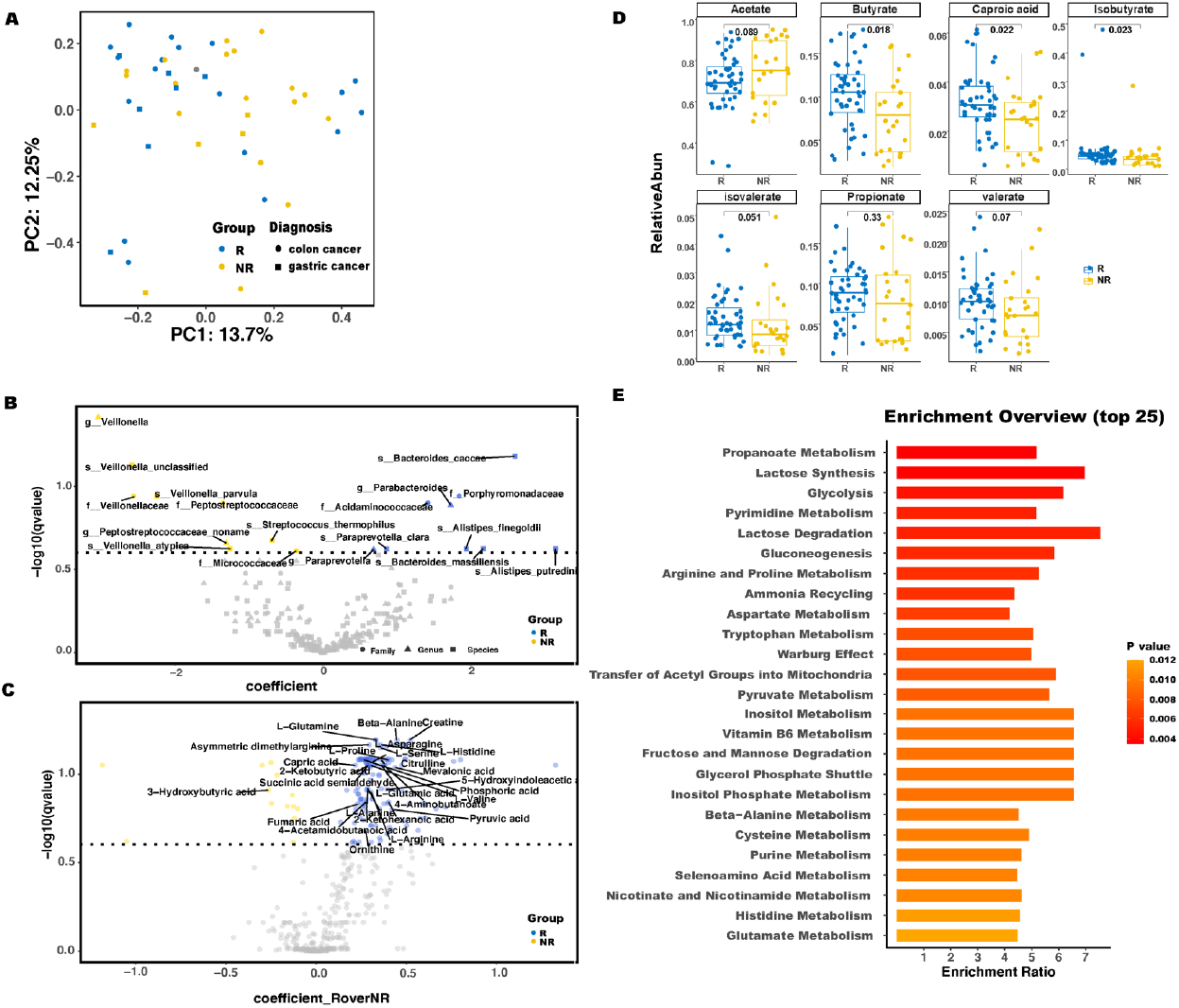
Gut microbiome composition, functional pathways and blood metabolites associated with drug response. **(A)** PCoA plot of fecal samples arranged by response and diagnosis using Bray–Curtis dissimilarity. The x- and y-axes show the first and second principal coordinates, along with the percentage of variance explained on each dimension. NR, non-responders; R, responders. **(B)** A volcano plot of taxa differentially abundant between responders and non-responders, based on MaasLin2 results. Taxa that differed significantly between groups (p-value <0.05) were named accordingly. **(C)** A volcano plot of metabolome differentially abundant between responders and non-responders, based on MaasLin2 results. Metabolome that differed significantly between groups (p-value <0.05) were named accordingly. **(D)** Differences in the relative abundance of different short-chain fatty acids between responders and non-responders, based on Wilcoxon test. **(E)** Top 25 functional pathways enriched in responders based on the untargeted metabolome data, sorted by p value.

To render an unbiased view of metabolic profiles of patients with different treatment responsiveness, we performed global metabolomic profiling using liquid chromatography-mass spectrometry (LC-MS) on baseline blood plasma samples. We detected and putatively identified 241 and 300 distinct spectral features by LC/MS [electrospray ionization (ESI)+] and LC/MS (ESI-) mode, respectively **(Table S2)**. Tight clustering of the QC samples in the PCA plot was observed in both the LC/MS (ESI+) and LC/MS (ESI-) modes (**Figure S3**), indicating good instrument stability and reproducibility of the obtained data. We compared the metabolite profiles of responders and non-responders to obtain potential pathways associated with primary drug resistance. In total, 143 metabolites were identified as differential between responders and non-responders (**Figure 2C &Table S3)**. Of these, two types of metabolites are noteworthy, including ones involved in arginine metabolism, such as L-arginine, ornithine, L-glutamine, asymmetric dimethylarginine, L-asparagine, L-serine, citrulline, L-proline. We also explored the relationship between arginine and gut microbes and found several microbes significantly associated with arginine quantities (**Table S4**), including *Paraprevotella clara* (*P*<0.05, cor=0.4) and *Veillonella atypica* (*P*<0.05, cor=−0.4), which were previously found to be differential features between responders and non-responders. The other type of metabolites is associated with short-chain fatty acids (SCFA) metabolism, such as 4-aminobutanoate, capric acid, 2-ketohexanoic acid, 4-acetamidobutanoic, 3-hydroxybutyric acid. Targeted metabolomics on SCFAs was also applied and we found butyrate, caproic acid and iso-butyrate were significantly enriched in responders (**Figure 2D**). Pathway enrichment analysis between responder and non-responder groups also showed significance in propanoate metabolism, which is related to SCFA metabolism, along with arginine and proline metabolism (**Figure 2E & Figure S4**).

### 3.3 Clinical benefits transferred by fecal microbiota transplantation in mice

To validate the role of the gut microbiome in mediating patients’ responses to immunotherapy, fecal samples from 6 patients (3 responders, 3 non-responders) were transplanted into an MC38 mouse model pre-treated with broad-spectrum antibiotics. While evaluating potential synergistic effects of FMT alongside anti-PD1 treatment (**Figure 3A**), we observed a significant reduction in tumor size in mice transplanted with responders’ fecal material (R group) compared to those receiving non-responders’ stool (NR group) or anti-PD1 alone (NC group) (**Figure 3B**). Using multiplex immunochemistry (mIHC) to appraise T-cell populations in the tumor microenvironments (TME), we observed increased fractions of helper T-cells (*P*=0.02) and cytotoxic T-cells (*P*=0.06) in mice receiving responders’ stool (**Figure 3C & 3D**). These results suggest that FMT can alter the immune cell landscape in TME and enhance immunotherapy.

**Figure 3.**
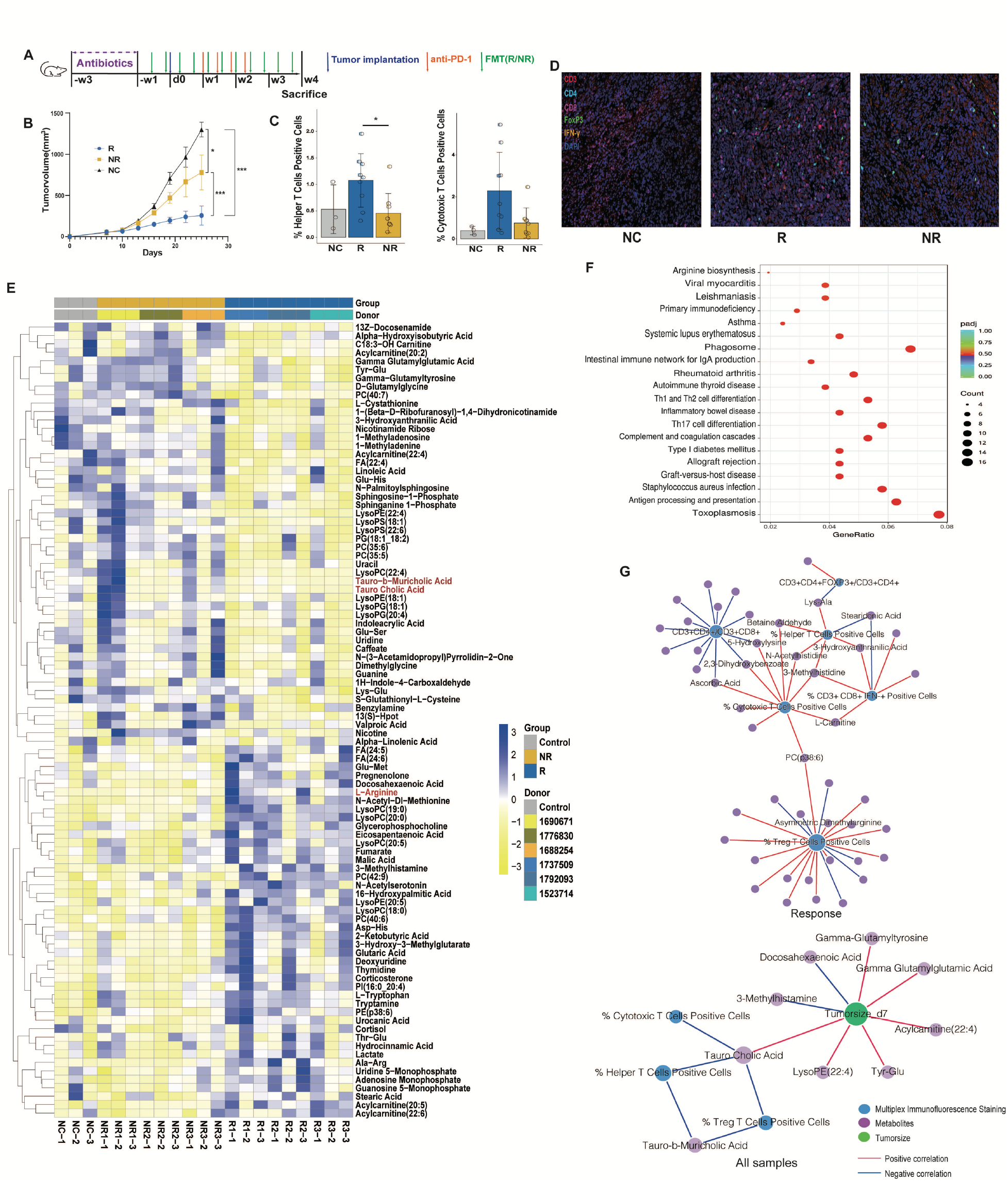
Enhanced anti-tumor effects in mice following responder fecal microbiota transplantation (FMT). **(A)** Experimental design. **(B)** Tumor growth curve. Graphic representation of the results of a one-way ANOVA followed by Tukey’s multiple comparisons test. **(C)** CD3+ CD4 helper and CD3+ CD8+ cytotoxic T-cell population in the tumor microenvironment, quantified through mIHC. Graphic representation of the results of a one-way ANOVA followed by Tukey’s multiple comparisons test. **(D)** Representative images of mIHC of tumor tissue samples. NC: negative control, R: responders, NR: non responders. Yellow=IFN-γ, green=Foxp3, magenta=CD8, red=CD3 and cyan=CD4. **(E)** Heatmap showed the differential metabolites identified from blood metabolome. **(F)** Dot plot showed significantly enriched KEGG pathways based on mouse tumor gene expression data. **(G)** Network analysis of tumor size, blood metabolites and T cell population in the tumor microenvironment. Spearman correlations were calculated between each feature comparison and those yielding p.adjust-value < 0.05 were retained. Node size indicates feature degree.

We further performed a multi-omics analysis to dissect the differences between the R and NR group, including metabolomics profiling of mouse blood samples and RNA-seq analysis of mouse tumor tissues. Total 97 differential metabolites were identified, including arginine, which was previously found in higher quantities in responder patients (**Figure 3E & Table S5**). RNA-seq pathway enrichment analysis also pinpointed arginine biosynthesis as significant (**Figure 3F**), in which the expression of four genes involved in the urea cycle (NOS2 / gm5424 / arg1 / arg2) was up-regulated in the R group (**Table S6**). These evidences indicate an active ammonia metabolism and arginine synthesis may impact the ICI response. Furthermore, we explored co-occurrence relationships between blood metabolites and tumor immune cell populations and found asymmetric dimethylarginine, a metabolite related with arginine metabolism, was significantly associated with T regulatory cells, indicating the role of arginine metabolism in modulating the immune function (**Figure 3G**).

### 3.4 Gut microbiome-based machine learning model well predicted the efficacy of immunotherapy

We sought to find out whether it is feasible to predict the response probability prior to ICI treatment and identify biomarkers associated with clinical outcomes using gut microbiome data. Three XGBoost models were built with microbial species data, microbial functional pathway data and both datasets together (**Figure 4**). To evaluate the robustness of the trained model, an external cohort comprising 29 stool samples from our in-house project was applied. We found the model integrated with both species and functional pathway information outperformed the others and achieved accuracy at 0.83 (AUC = 0.82) on the external dataset, indicating the additional predictive ability of functional pathway information (**Figure 4**). By analyzing feature contributions to the response status prediction by our model, *Bacteroides caccae, Vellonella*, phosphatidate metabolism, *Parabacteroides*, and *Bacteroides thetaiotaomicron* were top 5 indicative features to ICI response (**Figure 4F**).

**Figure 4.**
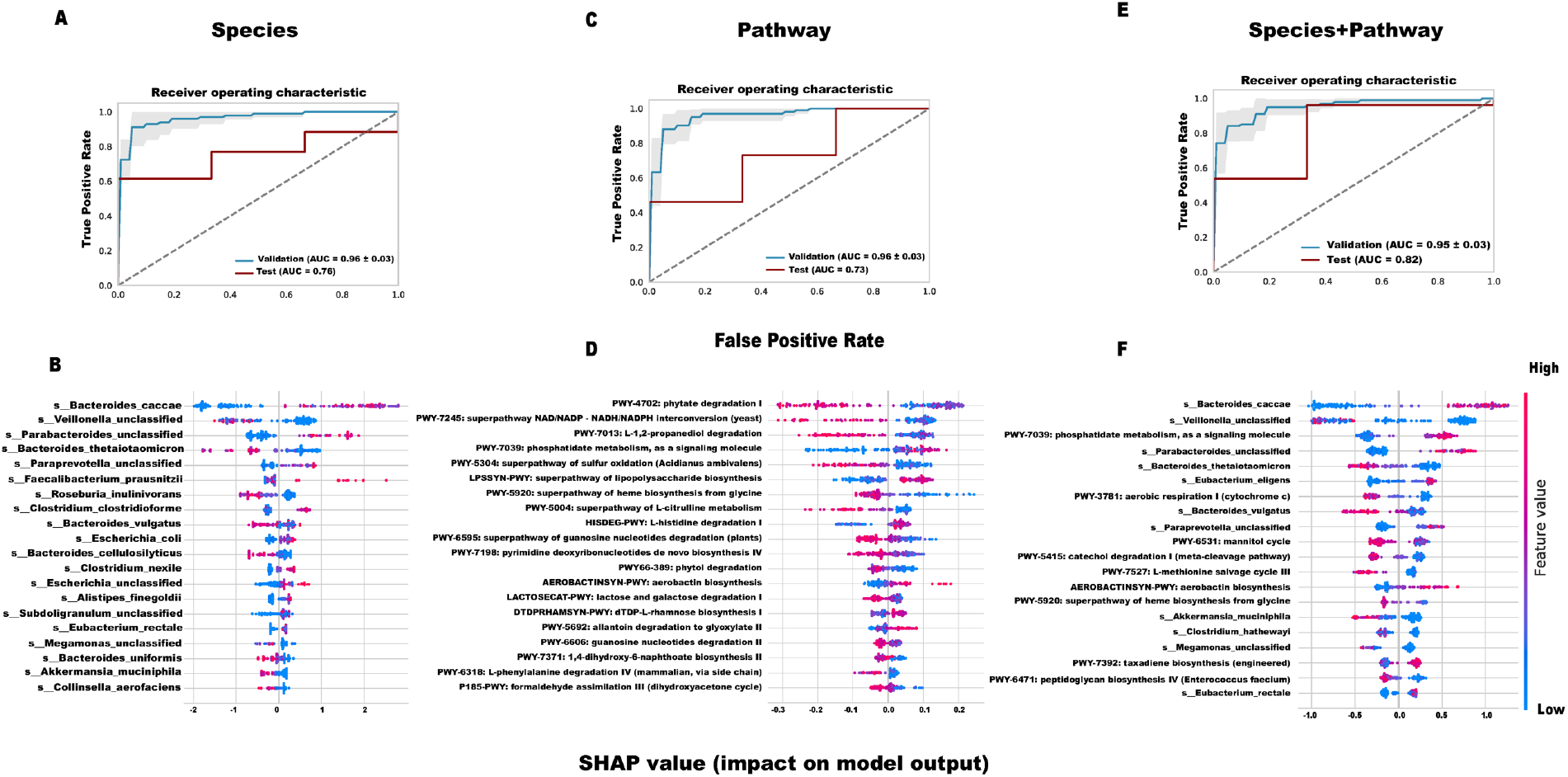
Machine learning model based on microbial biosignature. (A,C,E) Receiver operating characteristic(ROC) curve curves of machine learning models based on species(AUC=0.76), pathways (AUC=0.73) and species combined with pathways (AUC=0.82) respectively. **(B,D,F)** Feature contribution represented by the SHAP value in the above-mentioned three machine learning models based on species, pathways and species combined with pathways respectively.

### 3.5 Biosignatures of patients with acquired resistance to immunotherapy

During clinical practice, considerable number of patients develop acquired resistance (AR) to immunotherapy, defined as patients who fail to respond to ICI following objective response or prolonged SD for more than 6 months^27^. On the other hand, long responders (LR) were defined as sustained benefit over one year and without disease progression at the time of analysis. Thus, to understand the role of gut microbiome in the process of acquired resistance, we further divided responders into AR and LR group.

Baseline multi-omics analysis identified a few differential features (microbes, metabolites, or cytokines/chemokines), including significantly increased IL-5, hydrocinnamic acid, 2-hexadecenal and biliverdin in the AR group as well as salicyluric acid and glycoursodeoxycholic acid increased in the LR group (**Table S7**). To get a systematic view of gut microbiome’s role during acquired resistance, we further analyzed patients’ gut microbiome before and after they acquired ICI resistance. Total 18 species associated with drug resistance were identified (**Figure 5A**). Notably, two *Alistipes* species (*A. onderdonkii* and *A. putredinis*) found in lower quantities in acquired resistant samples were also decreased in the primary resistant group (NR), indicating their potential role in ICI resistance. Species such as *Ruminococcus bromii, Bifidobacterium adolescentis and Bifidobacterium pseudocatenulatum* significantly decreased during the drug resistance process and they were also found to increase along time in LR patients (**Figure 5B)**.

**Figure 5.**
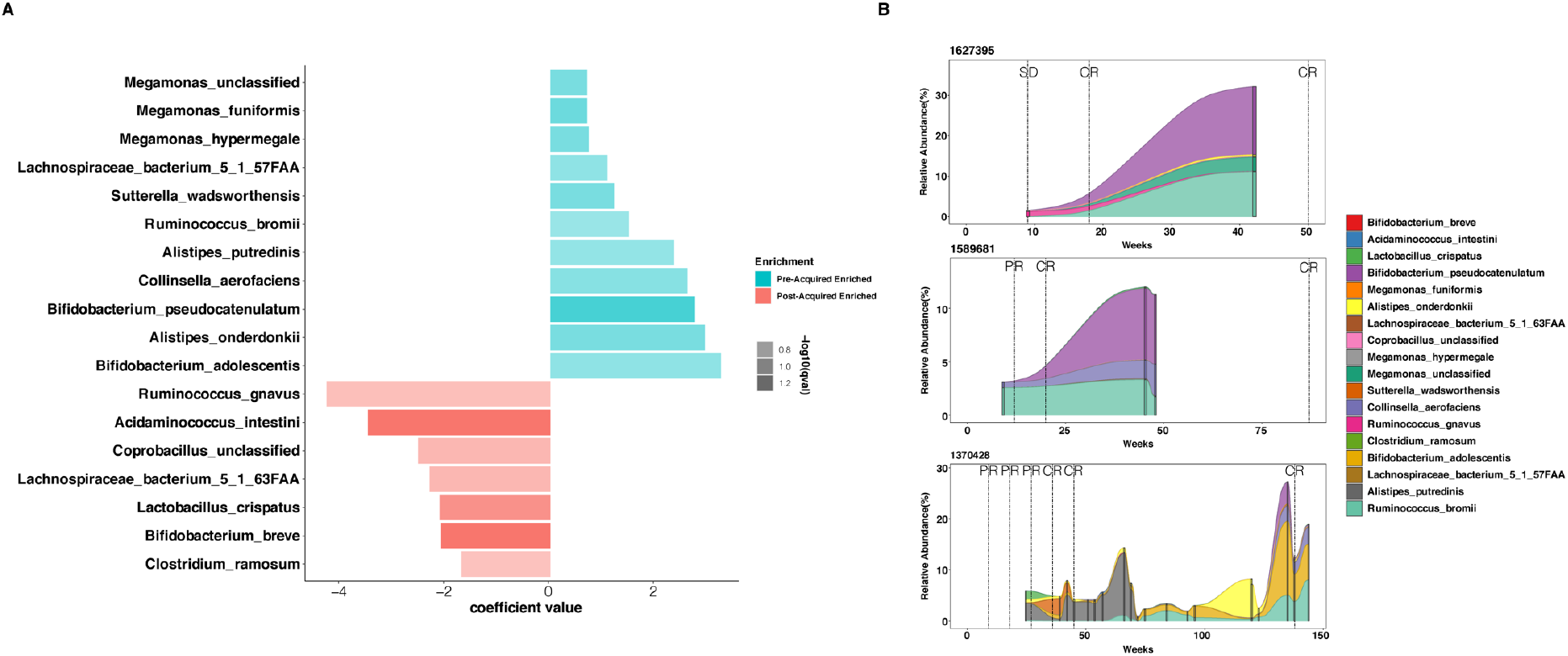
Gut microbiome associated with acquired resistance. **(A)** Differential microbes identified during the progression of drug resistance. **(B)** Microbial dynamics in representative LR patients, focusing on the differential microbes found during the progression of acquired resistance.

## 4. Discussion

The role of gut microbiome in mediating cancer immunotherapy has attracted increased attention in recent years^28,29^. While studies have shown that gut microbiome intervention can increase patients’ response rate to immunotherapy^9,11,17–19,30^, the results vary widely among cohorts. This is partially due to the heterogeneity of the patient populations studied. By studying a cohort of MSI-H/dMMR gastrointestinal cancer patients, we minimized the noise introduced by various disease subtypes. Although MSI-H/dMMR status is a key factor in determining whether a patient might benefit from ICI therapy, elucidating the interplay between gut microflora, host immune status, and drug resistance is of paramount importance in improving patient stratification.

We identified gut bacteria, blood metabolites, and corresponding functional pathways associated with primary resistance. Notably, L-arginine was enriched in responders of our cohort as well as the R group in the animal trial. L-arginine is a conditionally essential amino acid and its metabolism is regulated by intestinal flora^31^. Arginine is shown to promote immunotherapy by regulating T cell metabolism through three transcription regulators (baz1b, Psip1 and TSN)^32^. Multistage cooperative nanodrug based on arginine and cisplatin CpG have been shown to promote CD8+ T cell infiltration and exert a synergistic effect with anti-PD-L1 treatment both *in vitro* and *in vivo* ^33^. Engineered probiotic *Escherichia coli* Nissle 1917, which converts ammonia to L-arginine, could significantly increase T cell infiltration in the tumor microenvironment and enhance the efficacy of anti-PD-L1 immunotherapy ^34^. In the FMT animal trial, we found a significant increase in tumor-infiltrating helper T-cells and cytotoxic T-cells in mice that received microbiota transplantation from responders, which could possibly be due to the increased quantities of circulating arginine in the animals and needs further investigation. In this study, we also found the increase of certain SCFAs in responders, including butyrate. As the most studied microbiota-derived SCFAs, butyrate uncoupled the tricarboxylic acid cycle from glycolytic input in CD8^+^T cells and then enhanced memory potential of activated CD8^+^T cells^35^. He et al demonstrated that butyrate treatment directly boosted the antitumor cytotoxic CD8+ T cell responses both *in vitro* and *in vivo* in an ID2-dependent manner through the IL-12 signaling pathway^15^. Besides butyrate, capric acid and isobutyrate were also significantly enriched in responders from our MSI-H/dMMR cohort. Compared with butyrate, there is very limited research on the mechanisms by which caproic acid and isobutyrate regulate host immunity. Whether caproic acid and isobutyrate exhibit similar mechanisms to butyrate still needs further exploration. Our finding of the responder-enriched species and pathways that are capable of producing these SCFAs (*Porphyromonadaceae*) may provide extra evidence on the pro-immunotherapy effect of commensal bacteria^36^. We further developed machine learning models with the goal to evaluate the usability of the microbiome-related signature as anti-PD-1/PD-L1 immunotherapy biomarkers. With the combination of microbial composition and functional pathways, our predictive models exhibited excellent predictive performance in an independent external validation set. Because the molecular background of MSI-H/dMMR subtype patients are relatively homogenous, we believe the microbes/ functional pathways contributing to the models could potentially be applied as pan-tumor biomarkers in other cancer cohorts.

Another highlight of our work was the focus on acquired resistance. Having hypothesized that gut microflora plays an active role in mediating acquired resistance, we set out to identify significant differences in the gut microbiomes of AR patients before and after resistance. Several microbes possibly contributing to drug resistance were identified, including *Bifidobacterium*. Among the three *Bifidobacterium* species we identified (*B. adolescentis, B. pseudocatenulatum* and *B. breve*), the first two were enriched before acquired resistance while the last one was enriched after resistance occur. Although previous reports have shown commensal *Bifidobacterium* could promote immunotherapy efficacy in mouse models^17,37^, the effect of different species could vary as shown in our findings, considering the vast differences in the *Bifidobacterium* genus regarding host preference, metabolic functions etc ^38–40^. This again reminds us that the species and even strain variance should be taken into consideration when studying gut microbiome. Interestingly, *Ruminococcus bromii* was also found in descending trend during the progress of drug resistance. This species were also found to highly correlate with better clinical response in an on-going clinical trial evaluating the combination of PD1 and FMT in treating refractory metastatic gastrointestinal cancer (in-house data, not published). Meanwhile, we acknowledge the limited patient number in this analysis, and thus current findings warrant further confirmation in larger cohorts. Nevertheless, our preliminary investigation on acquired resistance is of paramount importance in understanding the role of gut microbiome during acquired resistance.

Interrogating a rare cohort of MSI-H/dMMR gastrointestinal cancer patients with a suite of integrated multi-omics analyses, we identified biosignatures whose differential presence/absence or relative abundance correlated significantly with primary or acquired modes of immunotherapeutic drug resistance. Especially, microbes, pathways and/or metabolites involved in arginine metabolism and SCFAs metabolism indicate certain common molecular mechanisms related to gut microbes and hosts’ response to immunotherapy. Robust machine learning models were also built to predict patient response to immuno-therapy in addition to the “golden” MSI status, which could greatly improve the accuracy of patient stratification to maximize treatment benefits. We believe these exciting findings will help to guide the clinical practice in cancer immunotherapy and further explore the feasibility and efficacy of FMT in reversing the immunotherapy resistance in gastrointestinal cancer patients.

## Author contributions

Y.K. and Z.P. conceptualized the study; X.Y. and X.G. performed the methodology; Resources, Z.P., S.C., Z.H., X.Z., M.L., Z.L., X.W., J.Z. and J.L. obtained the resources; S.C., Z.H., X.Y., D.D., F.L., N.Z. and Q.Z. performed the study investigation; P.S., D.D., F.L. performed the formal analysis; Z.P., S.C., Z.H. and X. Y. wrote the original draft; Y.T., Y.K. and L.S. reviewed and edited the article; Y.T., Y.K., Z.P. and L.S. acquired the funding. All authors have approved the manuscript.

## Data Availability Statement

Sequencing data generated in this study have been deposited in the China National GeneBank (CNGB) under accession number CNP0002257 (CNSA: db.cngb.org/cnsa).

## SUPPLEMENTARY FIGURES AND TABLES

**Figure S1.** Sample processing workflow

**Figure S2. (A)** Microbial alpha diversity between responders (R) and non-responders (NR); **(B)** Genus composition plot of each individual.

**Figure S3.** Principle component plots for blood metabolome **(A)** based on positive detection mode **(B)** and negative detection mode respectively.

**Figure S4. (A)** Comparison of metabolites involved in the propanoate metabolism pathway in responder and non-responder groups. The right panel shows a KEGG pathway with identified metabolites in red circle; **(B)** Comparison of metabolites involved in the arginine and proline metabolism pathway in responder and non-responder groups. The right panel shows a KEGG pathway with identified metabolites in red circle.

**Table S1.** Significantly altered species between R and NR patients.

**Table S2**. Characteristics of compound spectral features identified by LC/MS [electrospray ionization (ESI)+] and LC/MS (ESI-) mode.

**Table S3**. Differential metabolites between R and NR patients.

**Table S4**. Relationship between arginine and gut microbes.

**Table S5.** Differential metabolites between mice transplanted with R or NR patients’ stool samples.

**Table S6.** Differentially expressed host genes based on RNA-seq analysis.

**Table S7.** Differential metabolites between AR and LR patients’ baseline samples.

