## Supplementary figures and images for "Multi-omics of the gut microbial ecosystem in the immunotherapy resistance in microsatellite instability-high gastrointestinal cancer patients"

### Supplementary Figures

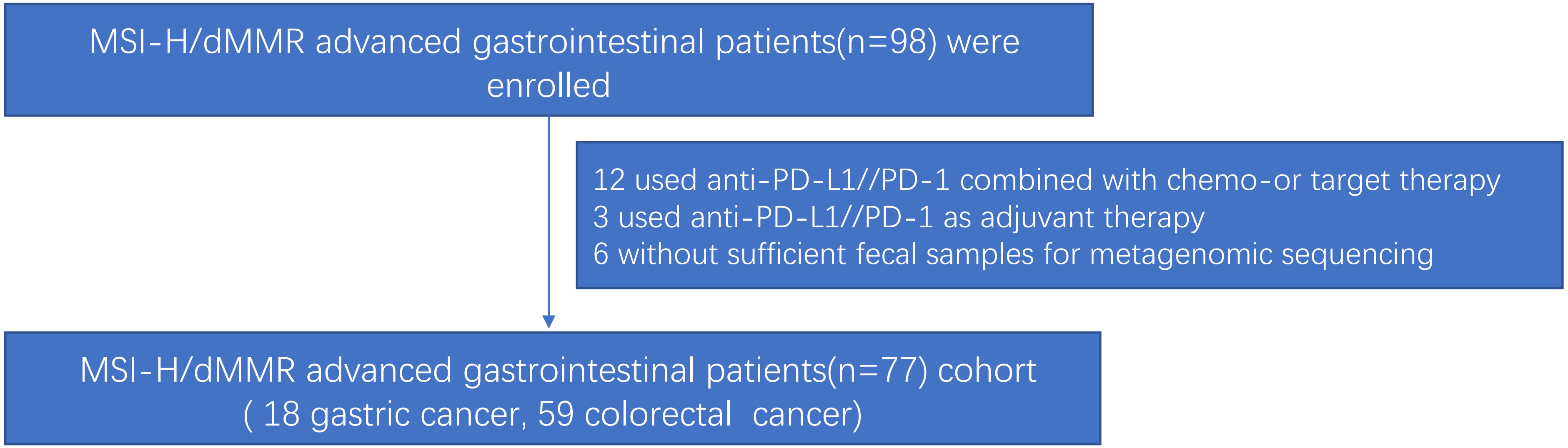

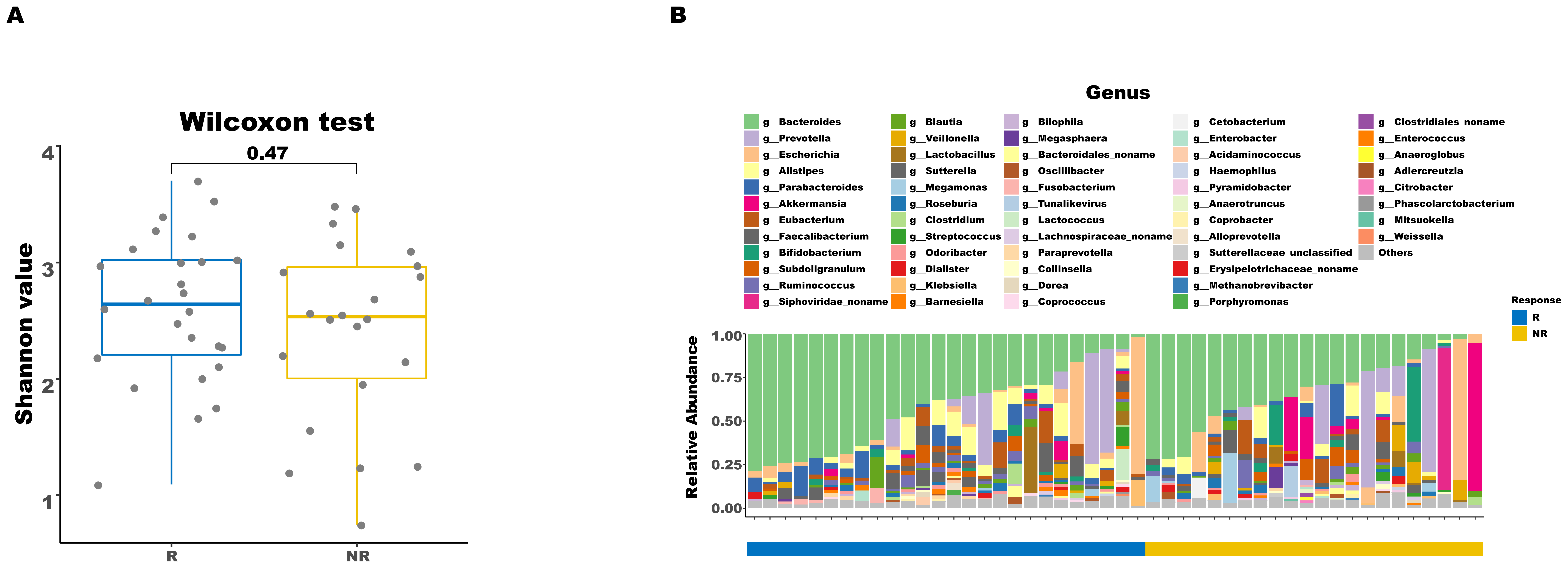

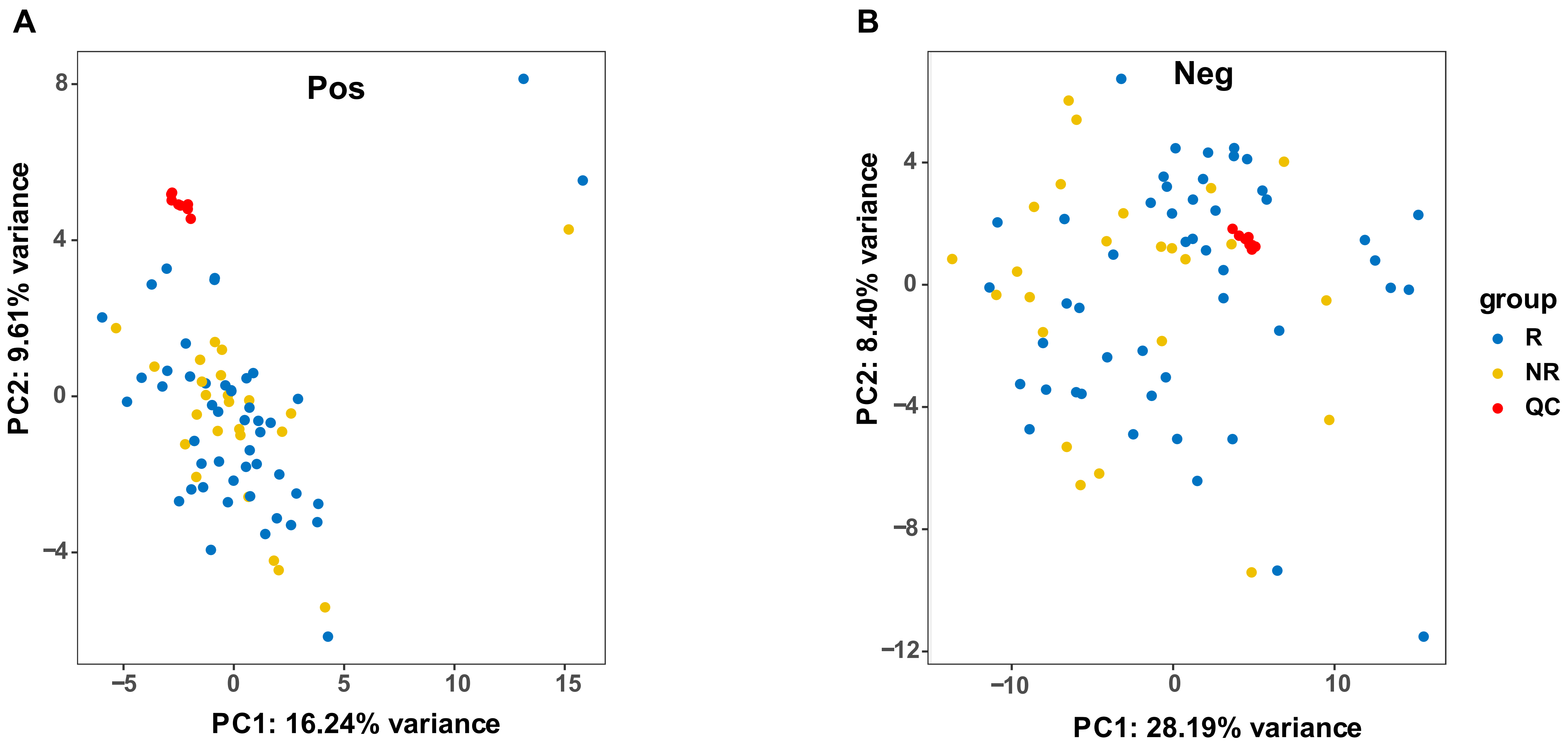

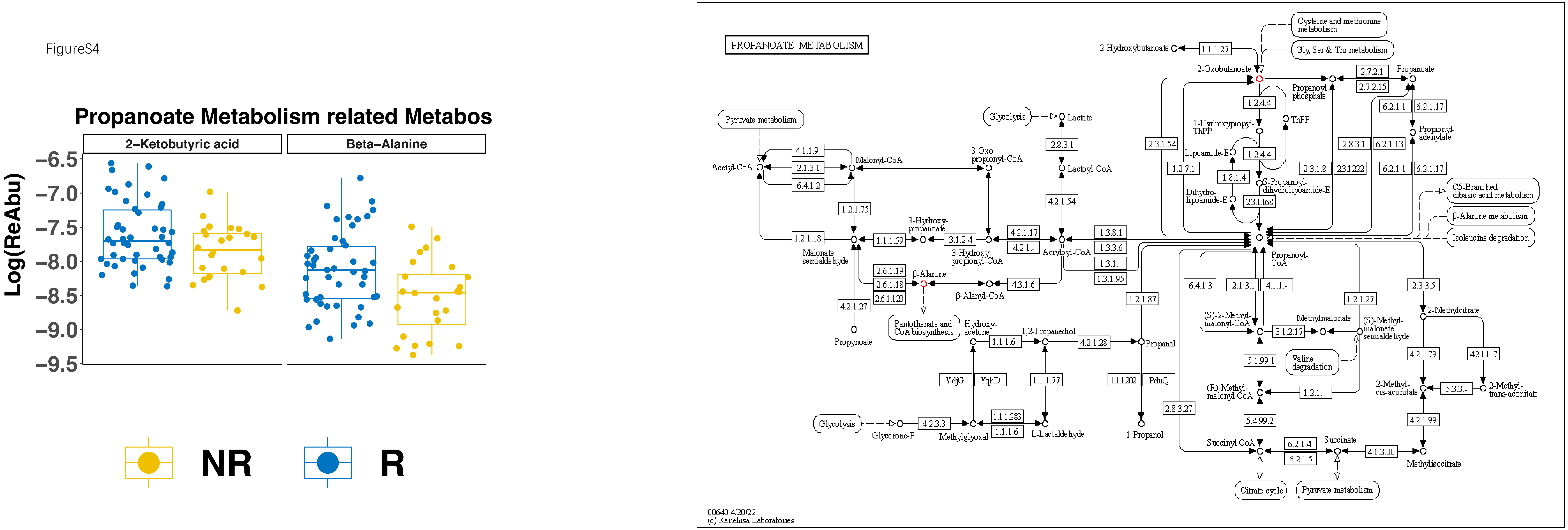

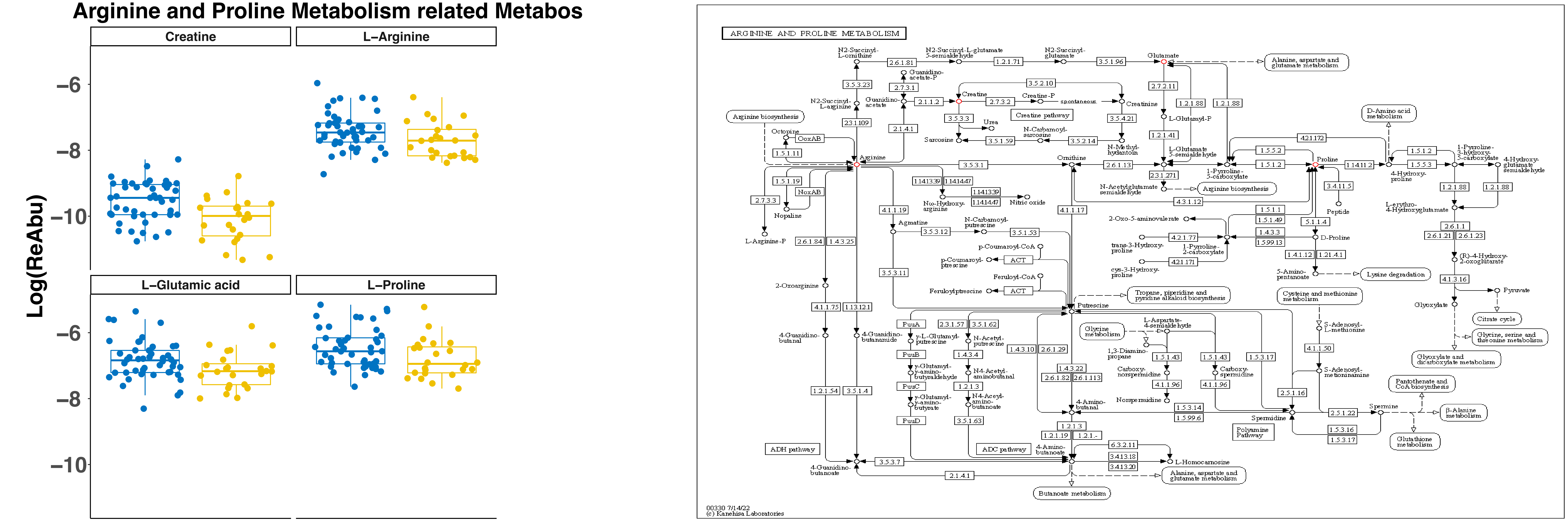
