## Supplementary Tables for "Multi-omics of the gut microbial ecosystem in the immunotherapy resistance in microsatellite instability-high gastrointestinal cancer patients"

**Table S1: Significantly altered species between R and NR patients.**

| feature | metadata | value | coef | pval | qval | mean_NR | mean_R | median_NR | median_R |
|---|---|---|---|---|---|---|---|---|---|
| s__Bacteroides_caccae | Response | R | 2.597638495 | 0.002712175 | 0.065947873 | 0.723720909 | 2.279236923 | 0 | 1.576855 |
| s__Veillonella_unclassified | Response | R | -2.582447105 | 0.003449699 | 0.074414931 | 1.087318182 | 0.411620385 | 0.123585 | 0.00508 |
| s__Veillonella_parvula | Response | R | -2.250203099 | 0.006075101 | 0.114667532 | 1.081590455 | 0.191419615 | 0.085745 | 0 |
| s__Streptococcus_thermophilus | Response | R | -0.68822098 | 0.017518621 | 0.211624936 | 0.012069545 | 0.000854231 | 0 | 0 |
| s__Bacteroides_massiliensis | Response | R | 2.171274778 | 0.021218102 | 0.237328399 | 0.403851818 | 2.305285385 | 0 | 0 |
| s__Alistipes_finegoldii | Response | R | 1.943154986 | 0.022614223 | 0.237593281 | 0.147247273 | 0.380416154 | 0 | 0.169415 |
| s__Alistipes_putredinis | Response | R | 3.146183955 | 0.022931116 | 0.237593281 | 1.537482727 | 3.537311923 | 0 | 1.82423 |
| s__Veillonella_atypica | Response | R | -1.256104427 | 0.023601982 | 0.237593281 | 0.163028636 | 0.0616 | 0.023965 | 0 |
| s__Paraprevotella_clara | Response | R | 0.861716263 | 0.024493137 | 0.238610558 | 0.032367727 | 0.128361154 | 0 | 0 |
| s__Paraprevotella_unclassified | Response | R | 0.751109206 | 0.028377102 | 0.259693475 | 0.095074091 | 0.352257692 | 0 | 0 |
| s__Barnesiella_intestinihominis | Response | R | 1.246105704 | 0.044107945 | 0.370016652 | 0.256836364 | 0.895272308 | 0 | 0 |
| s__Clostridiales_bacterium_1_7_47FAA | Response | R | -0.855428255 | 0.049338159 | 0.382054461 | 0.116279545 | 0.00407 | 0 | 0 |
| s__Alistipes_unclassified | Response | R | -0.466547266 | 0.05094759 | 0.384654303 | 0.063325909 | 0.003333077 | 0 | 0 |
| s__Bacteroides_salyersiae | Response | R | 1.049442393 | 0.053259595 | 0.392302383 | 0.064592727 | 0.364646154 | 0 | 0 |
| s__Megamonas_rupellensis | Response | R | -0.904521568 | 0.056002322 | 0.402683366 | 0.20412 | 0.001691538 | 0 | 0 |
| s__Klebsiella_oxytoca | Response | R | -0.476871435 | 0.062940893 | 0.413220645 | 0.010386818 | 0.000596923 | 0 | 0 |
| s__Bifidobacterium_adolescentis | Response | R | 1.093971455 | 0.068713018 | 0.432319403 | 0.016635455 | 0.149181923 | 0 | 0 |
| s__Paraprevotella_xylaniphila | Response | R | 0.556276001 | 0.075411596 | 0.455486038 | 0.002704091 | 0.013529615 | 0 | 0 |
| s__Parabacteroides_merdae | Response | R | 1.46845139 | 0.077448164 | 0.458614619 | 0.798959545 | 2.357328462 | 0 | 0.64041 |
| s__Lachnospiraceae_bacterium_2_1_58FAA | Response | R | -1.162458604 | 0.084309399 | 0.471508121 | 0.131515909 | 0.032496154 | 0.006855 | 0 |
| s__Lachnospiraceae_bacterium_3_1_57FAA_CT1 | Response | R | 0.473325647 | 0.083396353 | 0.471508121 | 0.000893636 | 0.015588846 | 0 | 0 |
| s__Alistipes_onderdonkii | Response | R | 1.453804989 | 0.095512568 | 0.489328737 | 0.639715909 | 0.832457692 | 0 | 0.34225 |
| s__Alistipes_shahii | Response | R | 1.643077174 | 0.102770526 | 0.489328737 | 0.897737727 | 1.380681923 | 0 | 0.55955 |
| s__Haemophilus_parainfluenzae | Response | R | -1.02857494 | 0.103698805 | 0.489328737 | 0.169679091 | 0.091366154 | 0.020465 | 0 |
| s__Megamonas_unclassified | Response | R | -1.559136788 | 0.095588214 | 0.489328737 | 1.759128636 | 0.174053077 | 0 | 0 |
| s__Prevotella_buccae | Response | R | -0.745035269 | 0.091306639 | 0.489328737 | 0.558554091 | 0.006166923 | 0 | 0 |
| s__Megasphaera_micronuciformis | Response | R | -0.776770423 | 0.117128057 | 0.531252223 | 0.051781818 | 0.045379231 | 0 | 0 |
| s__Rothia_mucilaginosa | Response | R | -0.291687303 | 0.117860593 | 0.531252223 | 0.005630909 | 0.001595769 | 0 | 0 |
| s__Bacteroidales_bacterium_ph8 | Response | R | 1.024098005 | 0.125763497 | 0.550443134 | 0.309205909 | 0.535330769 | 0 | 0 |
| s__Alistipes_senegalensis | Response | R | 0.871682529 | 0.157907224 | 0.583140801 | 0.069983182 | 0.058303077 | 0 | 0 |
| s__Bacteroides_clarus | Response | R | 0.727040126 | 0.138574173 | 0.583140801 | 0.005186818 | 0.079910385 | 0 | 0 |
| s__Citrobacter_unclassified | Response | R | -0.556738763 | 0.166059963 | 0.583140801 | 0.054966818 | 0.002026923 | 0 | 0 |
| s__Clostridium_clostridioforme | Response | R | 0.787442186 | 0.162730274 | 0.583140801 | 0.030794091 | 0.153514231 | 0 | 0 |
| s__Coprobacter_fastidiosus | Response | R | 0.845017294 | 0.153266161 | 0.583140801 | 0.091548182 | 0.152751154 | 0 | 0 |
| s__Dialister_invisus | Response | R | -1.304471457 | 0.144371614 | 0.583140801 | 0.852885 | 0.359066154 | 0.016265 | 0 |
| s__Lachnospiraceae_bacterium_3_1_46FAA | Response | R | -0.784133019 | 0.160952861 | 0.583140801 | 0.051091818 | 0.030222692 | 0 | 0 |
| s__Megamonas_hypermegale | Response | R | -1.016383559 | 0.147528136 | 0.583140801 | 0.347397273 | 0.02559 | 0 | 0 |
| s__Parabacteroides_johnsonii | Response | R | 0.413670304 | 0.155470881 | 0.583140801 | 0.000595909 | 0.018681538 | 0 | 0 |
| s__Citrobacter_freundii | Response | R | -0.443442214 | 0.188020426 | 0.620026173 | 0.031425 | 0.003401538 | 0 | 0 |
| s__Parasutterella_excrementihominis | Response | R | 0.979761969 | 0.195739169 | 0.624306439 | 0.057489091 | 0.154187692 | 0 | 0.039345 |
| s__Actinomyces_odontolyticus | Response | R | -0.209125798 | 0.224984709 | 0.65792859 | 0.004763182 | 0.001543846 | 0 | 0 |
| s__Alistipes_indistinctus | Response | R | 0.86533397 | 0.220175524 | 0.65792859 | 0.077279545 | 0.334888462 | 0 | 0 |
| s__Burkholderiales_bacterium_1_1_47 | Response | R | 0.883851274 | 0.231176018 | 0.662706049 | 0.071565909 | 0.122478846 | 0 | 0.030365 |
| s__Eubacterium_rectale | Response | R | 1.494404732 | 0.234709728 | 0.662706049 | 2.264841818 | 2.379109231 | 0.032785 | 0.75899 |
| s__Oxalobacter_formigenes | Response | R | 0.326982525 | 0.234799825 | 0.662706049 | 0.001225 | 0.010143846 | 0 | 0 |
| s__Fusobacterium_mortiferum | Response | R | 0.626293472 | 0.239334978 | 0.663111589 | 0.001578182 | 0.367649231 | 0 | 0 |
| s__Ruminococcus_callidus | Response | R | 0.984982677 | 0.243512351 | 0.66855209 | 0.115606364 | 0.296183846 | 0 | 0 |
| s__Bacteroides_sp_2_1_22 | Response | R | 0.540715467 | 0.269667358 | 0.689623832 | 0.017202727 | 0.370936538 | 0 | 0 |
| s__Bifidobacterium_pseudocatenulatum | Response | R | -0.977345907 | 0.266603686 | 0.689623832 | 1.560011818 | 0.161295769 | 0 | 0 |
| s__Streptococcus_parasanguinis | Response | R | -0.61388077 | 0.283506225 | 0.689623832 | 0.083666364 | 0.108462308 | 0.0126 | 0 |
| s__Streptococcus_vestibularis | Response | R | 0.275052559 | 0.280320138 | 0.689623832 | 0.004107727 | 0.014691538 | 0 | 0 |
| s__Subdoligranulum_unclassified | Response | R | 0.997806701 | 0.281517522 | 0.689623832 | 2.028221818 | 1.589636538 | 0.25554 | 0.522445 |
| s__Oscillibacter_unclassified | Response | R | 0.985481666 | 0.30072323 | 0.715105633 | 0.386275909 | 0.421143077 | 0.05563 | 0.118635 |
| s__Akkermansia_muciniphila | Response | R | -1.186783735 | 0.325707777 | 0.74490993 | 6.982830909 | 0.909342308 | 0.021785 | 0.01189 |
| s__Bilophila_unclassified | Response | R | 0.72513107 | 0.358236974 | 0.74490993 | 0.595849091 | 0.492404231 | 0.192255 | 0.39155 |
| s__Clostridium_bartlettii | Response | R | -0.402209562 | 0.330952819 | 0.74490993 | 0.023394545 | 0.016334231 | 0 | 0 |
| s__Collinsella_aerofaciens | Response | R | -0.738039409 | 0.331397058 | 0.74490993 | 0.350356818 | 0.191540769 | 0.07317 | 0.049465 |
| s__Fusobacterium_nucleatum | Response | R | -0.577086359 | 0.341538662 | 0.74490993 | 0.082268182 | 0.050631154 | 0 | 0 |
| s__Pyramidobacter_piscolens | Response | R | -0.623327355 | 0.340826815 | 0.74490993 | 0.203671818 | 0.063595769 | 0 | 0 |
| s__Ruminococcus_gnavus | Response | R | -0.782385727 | 0.346695373 | 0.74490993 | 0.830151364 | 1.022970385 | 0.288475 | 0.06122 |
| s__Bacteroides_thetaiotaomicron | Response | R | -0.750861675 | 0.390145996 | 0.770036814 | 2.345155909 | 2.098785769 | 1.40417 | 0.53581 |
| s__Megasphaera_unclassified | Response | R | 0.59237876 | 0.398081728 | 0.770645396 | 0.623546364 | 0.188555 | 0 | 0 |
| s__Anaerotruncus_unclassified | Response | R | 0.19467771 | 0.405583736 | 0.775229673 | 0.004397727 | 0.008798077 | 0 | 0 |
| s__Bacteroides_finegoldii | Response | R | 0.626255319 | 0.432401949 | 0.776217833 | 0.352320455 | 1.002053462 | 0 | 0 |
| s__Eubacterium_ramulus | Response | R | -0.420952096 | 0.428582823 | 0.776217833 | 0.057438636 | 0.025113462 | 0 | 0 |
| s__Faecalibacterium_prausnitzii | Response | R | 0.845907051 | 0.418938298 | 0.776217833 | 2.030580455 | 3.166515 | 1.483265 | 1.20491 |
| s__Klebsiella_pneumoniae | Response | R | -0.736137156 | 0.434373556 | 0.776217833 | 0.460932273 | 0.722291154 | 0.043005 | 0.01891 |
| s__Lachnospiraceae_bacterium_7_1_58FAA | Response | R | 0.313009204 | 0.415847959 | 0.776217833 | 0.013395 | 0.015741154 | 0 | 0 |
| s__Parabacteroides_unclassified | Response | R | 0.876822016 | 0.415323855 | 0.776217833 | 0.773731364 | 1.046231154 | 0 | 0.04067 |
| s__Ruminococcus_obeum | Response | R | 0.496222707 | 0.425771258 | 0.776217833 | 0.153649091 | 0.235073462 | 0.040305 | 0.058255 |
| s__Streptococcus_salivarius | Response | R | -0.471044512 | 0.451350136 | 0.783377822 | 0.287394091 | 0.52439 | 0.039025 | 0.01405 |
| s__Bifidobacterium_dentium | Response | R | -0.358508499 | 0.454405616 | 0.784174263 | 0.042135 | 0.015226923 | 0 | 0 |
| s__Veillonella_dispar | Response | R | -0.367804144 | 0.459427106 | 0.788335147 | 0.085802273 | 0.066781154 | 0 | 0 |
| s__Bifidobacterium_bifidum | Response | R | 0.41092316 | 0.473660919 | 0.803626952 | 0.042178636 | 0.048272308 | 0 | 0 |
| s__Clostridium_ramosum | Response | R | -0.159559746 | 0.472082165 | 0.803626952 | 0.004362727 | 0.002616154 | 0 | 0 |
| s__Megamonas_funiformis | Response | R | -0.201872798 | 0.4864321 | 0.807465729 | 0.022379545 | 0.017902308 | 0 | 0 |
| s__Fusobacterium_ulcerans | Response | R | 0.299440956 | 0.499022871 | 0.808246644 | 0.015714091 | 0.110289615 | 0 | 0 |
| s__Odoribacter_splanchnicus | Response | R | 0.459755761 | 0.497187612 | 0.808246644 | 0.560113182 | 0.677941538 | 0 | 0.09625 |
| s__Parabacteroides_distasonis | Response | R | 0.655582746 | 0.500470604 | 0.808246644 | 1.099531364 | 1.527457308 | 0.30569 | 0.26861 |
| s__Clostridium_nexile | Response | R | 0.378018311 | 0.525176 | 0.843633788 | 0.048727273 | 0.068449615 | 0 | 0 |
| s__Sutterella_wadsworthensis | Response | R | 0.477626384 | 0.535312597 | 0.850865285 | 1.063023636 | 1.524413077 | 0 | 0 |
| s__Coprobacillus_unclassified | Response | R | -0.290906441 | 0.541369099 | 0.851683471 | 0.016851364 | 0.027612308 | 0 | 0 |
| s__Alistipes_sp_AP11 | Response | R | 0.305422017 | 0.552807897 | 0.858724969 | 0.076307727 | 0.153930385 | 0 | 0 |
| s__Roseburia_unclassified | Response | R | 0.294164171 | 0.553501982 | 0.858724969 | 0.020105909 | 0.033540385 | 0 | 0 |
| s__Bacteroides_uniformis | Response | R | 0.512305872 | 0.578303424 | 0.868893701 | 4.752472273 | 4.801308462 | 1.78212 | 1.67192 |
| s__Clostridium_asparagiforme | Response | R | 0.271357867 | 0.568440685 | 0.868893701 | 0.022655 | 0.026783462 | 0 | 0 |
| s__Eubacterium_eligens | Response | R | 0.543132018 | 0.57099069 | 0.868893701 | 0.38746 | 0.429451154 | 0 | 0.07118 |
| s__Peptostreptococcaceae_noname_unclassified | Response | R | -0.121803135 | 0.577340704 | 0.868893701 | 0.010540909 | 0.006693462 | 0 | 0 |
| s__Eubacterium_ventriosum | Response | R | 0.372776496 | 0.596756015 | 0.892179784 | 0.052597727 | 0.146765 | 0.01509 | 0.006075 |
| s__Bilophila_wadsworthia | Response | R | -0.26698689 | 0.604014235 | 0.894177936 | 0.043285455 | 0.027183077 | 0.014095 | 0.01549 |
| s__Gordonibacter_pamelaeae | Response | R | -0.168649912 | 0.601123866 | 0.894177936 | 0.011086818 | 0.007626923 | 0 | 0 |
| s__Bacteroides_vulgatus | Response | R | 0.600630519 | 0.62006853 | 0.897552638 | 7.589899091 | 8.375409231 | 2.545105 | 2.16212 |
| s__Coprococcus_catus | Response | R | -0.273736298 | 0.613655651 | 0.897552638 | 0.055908182 | 0.02808 | 0 | 0 |
| s__Escherichia_unclassified | Response | R | -0.430425593 | 0.621153978 | 0.897552638 | 1.692913636 | 1.052281538 | 0.242325 | 0.3494 |
| s__Dorea_formicigenerans | Response | R | -0.228188006 | 0.655889362 | 0.902411093 | 0.072008182 | 0.043468462 | 0.022785 | 0.020435 |
| s__Eubacterium_hallii | Response | R | 0.329084534 | 0.636319051 | 0.902411093 | 0.107005455 | 0.20366 | 0.02499 | 0.04809 |
| s__Flavonifractor_plautii | Response | R | 0.239593753 | 0.643840605 | 0.902411093 | 0.042153182 | 0.033507692 | 0 | 0.00568 |
| s__Holdemania_filiformis | Response | R | -0.174702881 | 0.652146625 | 0.902411093 | 0.015675909 | 0.013446923 | 0 | 0 |
| s__Lachnospiraceae_bacterium_1_1_57FAA | Response | R | -0.300130221 | 0.647340788 | 0.902411093 | 0.129314545 | 0.144149231 | 0.01264 | 0.020835 |
| s__Lactobacillus_mucosae | Response | R | 0.160628661 | 0.652250555 | 0.902411093 | 0.015632727 | 0.018836154 | 0 | 0 |
| s__Prevotella_nigrescens | Response | R | -0.228246284 | 0.647761465 | 0.902411093 | 0.19181 | 0.040950769 | 0 | 0 |
| s__Roseburia_intestinalis | Response | R | 0.384126934 | 0.657385564 | 0.902411093 | 0.196479545 | 0.250557692 | 0 | 0.020755 |
| s__Roseburia_inulinivorans | Response | R | -0.386036769 | 0.640285753 | 0.902411093 | 0.523952273 | 0.32715 | 0.149285 | 0.11292 |
| s__Bacteroides_coprocola | Response | R | 0.269563408 | 0.705509567 | 0.915935232 | 0.790878182 | 2.151569615 | 0 | 0 |
| s__Bacteroides_dorei | Response | R | 0.448881327 | 0.699156266 | 0.915935232 | 1.845214545 | 1.461310769 | 0 | 0 |
| s__Bacteroides_eggerthii | Response | R | 0.239819978 | 0.679435317 | 0.915935232 | 0.427380909 | 1.391431154 | 0 | 0 |
| s__Bacteroides_intestinalis | Response | R | 0.341897598 | 0.718741928 | 0.915935232 | 0.475967727 | 0.257415769 | 0 | 0 |
| s__Bacteroides_xylanisolvens | Response | R | 0.306167249 | 0.710237917 | 0.915935232 | 0.170623636 | 0.227250769 | 0.03327 | 0.03587 |
| s__Clostridium_bolteae | Response | R | 0.282132427 | 0.709969177 | 0.915935232 | 0.197992727 | 0.234649231 | 0.04033 | 0.02811 |
| s__Clostridium_symbiosum | Response | R | 0.227496952 | 0.718796854 | 0.915935232 | 0.050849545 | 0.07507 | 0 | 0 |
| s__Holdemania_unclassified | Response | R | -0.077942764 | 0.70771975 | 0.915935232 | 0.003304091 | 0.002328846 | 0 | 0 |
| s__Lachnospiraceae_bacterium_8_1_57FAA | Response | R | 0.16677565 | 0.696239569 | 0.915935232 | 0.026655455 | 0.046767308 | 0 | 0 |
| s__Prevotella_copri | Response | R | 0.406312082 | 0.679349441 | 0.915935232 | 6.707398182 | 6.746433077 | 0 | 0 |
| s__Streptococcus_australis | Response | R | -0.077155961 | 0.697280376 | 0.915935232 | 0.003832273 | 0.002659615 | 0 | 0 |
| s__Dorea_unclassified | Response | R | -0.124871751 | 0.731680828 | 0.920698375 | 0.011428636 | 0.010667692 | 0 | 0 |
| s__Anaerostipes_unclassified | Response | R | 0.062424433 | 0.739698906 | 0.923948101 | 0.003189545 | 0.004961923 | 0 | 0 |
| s__Eggerthella_unclassified | Response | R | -0.157179962 | 0.740382253 | 0.923948101 | 0.033500455 | 0.036236923 | 0.007785 | 0 |
| s__Streptococcus_anginosus | Response | R | -0.135916056 | 0.746541995 | 0.927801162 | 0.026465455 | 0.039176538 | 0 | 0 |
| s__Clostridium_citroniae | Response | R | -0.141658426 | 0.750350436 | 0.928712425 | 0.020378182 | 0.021755385 | 0 | 0 |
| s__Bifidobacterium_longum | Response | R | 0.305886177 | 0.75647876 | 0.932475859 | 1.762060909 | 0.674095 | 0.039025 | 0.09834 |
| s__Eubacterium_siraeum | Response | R | 0.15995437 | 0.776219127 | 0.945138212 | 0.141190909 | 0.256506923 | 0 | 0 |
| s__Adlercreutzia_equolifaciens | Response | R | -0.123864878 | 0.80268371 | 0.955267646 | 0.068862273 | 0.022423462 | 0 | 0 |
| s__Clostridium_leptum | Response | R | 0.165549714 | 0.79760619 | 0.955267646 | 0.062944091 | 0.161824615 | 0 | 0 |
| s__Anaerotruncus_colihominis | Response | R | -0.138076259 | 0.837447999 | 0.959697023 | 0.189018636 | 0.058595769 | 0 | 0 |
| s__Bacteroides_fragilis | Response | R | -0.219051534 | 0.834646415 | 0.959697023 | 1.516592727 | 1.864118846 | 0.0453 | 0.012945 |
| s__Bacteroides_plebeius | Response | R | 0.188542606 | 0.812317332 | 0.959697023 | 1.214130455 | 1.846947692 | 0 | 0 |
| s__Clostridium_hathewayi | Response | R | -0.149669367 | 0.818763233 | 0.959697023 | 0.133651818 | 0.05921 | 0 | 0.008525 |
| s__Lactobacillus_salivarius | Response | R | -0.15068457 | 0.818973479 | 0.959697023 | 0.406966818 | 0.232339615 | 0 | 0 |

| | | | | | | | | | |
|---|---|---|---|---|---|---|---|---|---|
| s__Parabacteroides_goldsteinii | Response | R | -0.13379403 | 0.836111002 | 0.959697023 | 0.205455455 | 0.043324231 | 0 | 0 |
| s__Ruminococcus_bromii | Response | R | 0.136513014 | 0.845556601 | 0.959992833 | 1.379211818 | 1.115798077 | 0.202565 | 0.18728 |
| s__Ruminococcus_lactaris | Response | R | 0.118579046 | 0.865519014 | 0.965232509 | 0.111897273 | 0.144268077 | 0 | 0 |
| s__Ruminococcus_sp_5_1_39BFAA | Response | R | 0.086075607 | 0.86906867 | 0.965232509 | 0.393742273 | 0.179620385 | 0 | 0 |
| s__Subdoligranulum_sp_4_3_54A2FAA | Response | R | -0.06858097 | 0.861113631 | 0.965232509 | 0.192374545 | 0.225420385 | 0 | 0 |
| s__Bacteroides_ovatus | Response | R | -0.134952432 | 0.874202943 | 0.967066992 | 2.63617 | 2.189741154 | 0.72168 | 0.65524 |
| s__Bacteroides_cellulosilyticus | Response | R | 0.140497202 | 0.900652895 | 0.970088799 | 0.820161818 | 0.768847692 | 0.00744 | 0.01523 |
| s__Coprococcus_sp_ART55_1 | Response | R | 0.047884741 | 0.892946728 | 0.970088799 | 0.041463182 | 0.092994231 | 0 | 0 |
| s__Dorea_longicatena | Response | R | 0.091494334 | 0.900062728 | 0.970088799 | 0.163159091 | 0.232872692 | 0.0346 | 0.03829 |
| s__Roseburia_hominis | Response | R | 0.133002554 | 0.890679805 | 0.970088799 | 0.674591818 | 0.380433846 | 0.011195 | 0.14669 |
| s__Enterobacter_cloacae | Response | R | 0.071323355 | 0.906903588 | 0.971222991 | 0.082165455 | 0.266923077 | 0 | 0 |
| less_0.01p_total | Response | R | -0.021408062 | 0.917514344 | 0.975104527 | 0.086222727 | 0.090284231 | 0.077285 | 0.088735 |
| s__Bacteroides_faecis | Response | R | -0.0782871 | 0.920214537 | 0.975104527 | 0.271202273 | 0.140682692 | 0 | 0 |
| s__Escherichia_coli | Response | R | -0.109347052 | 0.91627027 | 0.975104527 | 5.556951364 | 5.759753462 | 0.583185 | 1.5324 |
| s__Eggerthella_lenta | Response | R | 0.01784404 | 0.941886195 | 0.984254778 | 0.004580909 | 0.008606154 | 0 | 0 |
| s__Lachnospiraceae_bacterium_5_1_63FAA | Response | R | -0.031042552 | 0.941761508 | 0.984254778 | 0.019755909 | 0.018313462 | 0.0063 | 0.00544 |
| s__Bacteroides_nordii | Response | R | 0.025375847 | 0.970108652 | 0.99865224 | 0.063219091 | 0.11355 | 0 | 0 |
| s__Ruminococcus_torques | Response | R | -0.026131725 | 0.96734544 | 0.99865224 | 0.502605455 | 0.573716923 | 0.2357 | 0.211695 |
| s__Anaerostipes_hadrus | Response | R | 0.002694349 | 0.99236524 | 0.999657047 | 0.011113636 | 0.011069615 | 0 | 0 |
| s__Bacteroides_stercoris | Response | R | -0.019157061 | 0.984494261 | 0.999657047 | 5.758554091 | 7.896688462 | 0.536585 | 0.41843 |
| s__Klebsiella_unclassified | Response | R | -0.00195109 | 0.993904964 | 0.999657047 | 0.0056 | 0.004201154 | 0 | 0 |
| s__Prevotella_stercorea | Response | R | -0.001701751 | 0.996346924 | 0.999657047 | 1.721553636 | 1.224924615 | 0 | 0 |
| s__Coprococcus_comes | Response | R | -0.000108216 | 0.999886085 | 0.999886085 | 0.106237727 | 0.175653462 | 0.054955 | 0.029755 |

### Table S2: Characteristics of compound spectral features identified by LC/MS [electrospray ionization (ESI)+] and LC/MS (E

| Compound Name | m/z (Expected) | RT | Library Score (%) | m/z (Delta (ppm)) | detection mode |
|---|---|---|---|---|---|
| (+)-a-Pinene | 137.13248 | 0.94552678 | N/A | -3.132475082 | positive |
| 2,4,5-Trimethoxybenzaldehyde | 197.08084 | 6.3132234 | 60 | -3.147406721 | positive |
| His-Phe | 303.14626 | 9.950714 | 52 | -3.0218638 | positive |
| Hypoxanthine | 137.04631 | 4.7687199 | 63 | -6.612525 | positive |
| Imidazole-4-acetaldehyde | 111.05529 | 11.365847 | N/A | -2.531417623 | positive |
| Indole | 118.06513 | 0.95031926 | 65 | -2.53595123 | positive |
| Indoleacetaldehyde | 160.07569 | 6.2903279 | 43 | -3.497407541 | positive |
| Indoleacrylic acid | 188.07061 | 8.1058963 | 95 | -3.343510574 | positive |
| 2,6-Dimethylheptanoyl carnitine | 302.23261 | 3.6473687 | 64 | -3.963842917 | positive |
| L-Alanine | 90.05495 | 9.7598313 | 33 | -1.489241721 | positive |
| L-Arginine | 175.11948 | 11.525388 | 95 | -6.317496803 | positive |
| L-Asparagine | 133.06132 | 10.400895 | 71 | -7.226192479 | positive |
| lauroylcarnitine | 344.27954 | 3.3643083 | 93 | -3.87767459 | positive |
| L-Carnitine | 162.11247 | 9.2915536 | 79 | -3.690721393 | positive |
| L-Cystine | 241.03165 | 13.217382 | 99 | -5.897413852 | positive |
| Leu-Leu | 245.18706 | 6.9066408 | 33 | -6.938001885 | positive |
| L-Glutamic acid | 148.06096 | 10.424103 | 87 | -6.992784016 | positive |
| L-Glutamine | 147.07694 | 10.278572 | 80 | -6.86044877 | positive |
| L-Histidine | 156.07727 | 11.342309 | 82 | -6.255265738 | positive |
| 2-Indolecarboxylic acid | 162.05496 | 9.344051 | 27 | 2.869704426 | positive |
| L-Histidine trimethylbetaine | 198.12425 | 10.448942 | 12 | -5.936852869 | positive |
| L-Homocitrulline/N6-Carbamoyl-DL-Lysine | 190.11917 | 10.341124 | 74 | -5.968980083 | positive |
| Linoleic acid | 281.24751 | 0.95330478 | 10 | -3.73350582 | positive |
| linolenyl carnitine | 422.32703 | 2.9892395 | 88 | -4.971697333 | positive |
| Linoleoyl Ethanolamide | 324.28971 | 0.90568418 | 54 | -3.692120492 | positive |
| Linoleyl carnitine | 424.34216 | 2.937537 | 100 | -3.918218167 | positive |
| L-Isoleucine | 132.10243 | 8.6889587 | 48 | -6.852801885 | positive |
| L-Kynurenine | 209.09259 | 8.1530345 | 90 | -5.724653361 | positive |
| L-Leucine | 132.10243 | 8.3503587 | 43 | -6.852801885 | positive |
| L-Lysine | 147.11333 | 11.78701 | 86 | -6.785904262 | positive |
| 2-Keto-6-aminocaproate | 146.08117 | 2.2235331 | N/A | -3.241476803 | positive |
| L-Methionine | 150.05885 | 8.9948822 | 70 | -6.047358926 | positive |
| L-Methionine S-oxide | 166.05324 | 10.260167 | 10 | -3.244686942 | positive |
| L-Octanoylcarnitine | 288.21696 | 4.1014708 | 70 | -4.058403388 | positive |
| L-Palmitoylcarnitine | 400.34214 | 2.9506594 | 100 | -3.806881721 | positive |
| L-Phenylalanine | 166.08678 | 8.1270488 | 70 | -6.163069754 | positive |
| L-Pipecolic acid | 130.08678 | 11.789937 | 30 | -6.582205246 | positive |
| L-Pipecolic acid-1 | 130.08678 | 8.4 | 30 | -6.582205246 | positive |
| L-Proline | 116.07113 | 11.896994 | 58 | -7.183176721 | positive |
| L-Proline-1 | 116.07113 | 9.34 | 58 | -7.183176721 | positive |
| L-Serine | 106.04987 | 10.324685 | 32 | -2.177770909 | positive |
| 2-Methylbutyrylglycine | 160.09682 | 9.3218864 | 13 | -2.3506 | positive |
| L-threonine | 120.06604 | 9.9438453 | 76 | -6.369632213 | positive |
| L-tryptophan | 205.09768 | 8.0997126 | 91 | -5.796853115 | positive |
| L-Urobilin | 595.34956 | 1.3890573 | N/A | -4.489928488 | positive |
| LysoPC(14:0) | 468.30901 | 6.2645404 | 100 | -4.351152623 | positive |
| LysoPC(16:0) | 496.34031 | 6.1479302 | 100 | -4.786232049 | positive |
| LysoPC(17:0) | 510.35542 | 6.0869572 | 100 | -2.590882149 | positive |
| LysoPC(17:1) | 508.33977 | 6.118697 | 100 | -4.078298917 | positive |
| LysoPC(18:0) | 524.37161 | 6.052027 | 100 | -4.961240661 | positive |
| LysoPC(18:1) | 522.35596 | 6.0864892 | 100 | -4.419754876 | positive |
| LysoPC(18:2) | 520.34031 | 6.1181551 | 100 | -4.843143917 | positive |
| 2-Pyrrolidinone | 86.06004 | 1.9641266 | 57 | -2.180454426 | positive |
| LysoPC(18:3) | 518.32412 | 6.1808445 | 100 | -4.238044583 | positive |
| LysoPC(19:0) | 538.38672 | 6.011087 | 100 | -1.72644125 | positive |
| LysoPC(20:0) (Low) | 552.40237 | 5.9675952 | 94 | -2.516060083 | positive |
| LysoPC(20:1) (Low) | 550.38726 | 5.9748312 | 99 | -4.285296333 | positive |
| LysoPC(20:2) | 548.37161 | 6.0500824 | 88 | -2.810757951 | positive |
| LysoPC(20:4) | 544.34031 | 6.0457706 | 100 | -4.756886942 | positive |
| LysoPC(22:4) | 572.37161 | 5.9766239 | 99 | -4.3000685 | positive |
| LysoPC(22:5) | 570.35596 | 6.0081512 | 98 | -4.072560248 | positive |
| LysoPC(22:6) | 568.34031 | 5.9953034 | 100 | -4.87262675 | positive |
| LysoPE(15:0) | 440.27717 | 6.4171773 | 11 | -3.002348917 | positive |
| 2-Pyrrolidinone-1 | 86.06004 | 2.72 | 57 | -2.180454426 | positive |

| | | | | | |
|---|---|---|---|---|---|
| LysoPE(16:0) | 454.29336 | 6.563035 | 100 | -4.345978833 | positive |
| LysoPE(18:1) | 480.30901 | 6.5037749 | 90 | -4.006435 | positive |
| LysoPE(18:1e) | 466.3292 | 6.1866007 | 21 | -3.124901 | positive |
| LysoPE(18:2e)+H | 464.31355 | 6.2090596 | N/A | -2.737918167 | positive |
| LysoPE(20:4) | 502.29336 | 6.43528 | 100 | -4.365078417 | positive |
| LysoPE(22:5) | 528.30901 | 6.3965853 | N/A | -2.547786667 | positive |
| Malonylcarnitine | 248.11288 | 10.269985 | 80 | -3.522311322 | positive |
| Methyladenine | 150.07743 | 9.0026005 | 10 | -3.351630579 | positive |
| Methylimidazoleacetic acid | 141.06585 | 9.5168478 | 35 | -2.968022295 | positive |
| 3 alpha,7 alpha,26-Trihydroxy-5beta-cholestane | 421.36762 | 0.88856586 | N/A | -3.322295083 | positive |
| Myristoylcarnitine | 372.31084 | 3.1284131 | 60 | -3.414990902 | positive |
| N-(3-acetamidopropyl)pyrrolidin-2-one | 185.12845 | 1.8969931 | N/A | -3.055789344 | positive |
| N1-Acetylspermidine | 188.17574 | 10.616276 | 71 | -2.662071475 | positive |
| N6,N6,N6-Trimethyl-L-lysine | 189.15975 | 11.148699 | 42 | -2.893318033 | positive |
| N-acetyl-D-tryptophan | 247.10824 | 6.3087676 | 20 | -5.359662213 | positive |
| N-Acetylglycine | 118.04987 | 8.5736215 | 60 | 4.651208099 | positive |
| N-Acetyl-L-alanine | 132.06552 | 9.7725449 | 29 | -2.792918033 | positive |
| N-Acetylputrescine | 131.11789 | 8.1941665 | 45 | -2.638233033 | positive |
| N-alpha-acetyl-L-asparagine | 175.07186 | 8.4129735 | 30 | -5.427847934 | positive |
| N-hexadecanoylsphinganine-1-phosphocholine | 705.59105 | 5.5982893 | N/A | -4.815708417 | positive |
| 3a,7a,12a-Trihydroxy-5b-cholestanoic acid | 451.3418 | 0.94980975 | N/A | -3.798447833 | positive |
| Niacinamide (VB3) | 123.05581 | 1.6490343 | 70 | -7.284761557 | positive |
| N-Palmitoylsphingosine | 538.51938 | 0.88825718 | 72 | -2.715591322 | positive |
| Octadecylamine | 270.31553 | 2.3702263 | 45 | -3.662663934 | positive |
| Oleic Acid | 283.26316 | 0.908915 | N/A | -3.388095246 | positive |
| Oleoyl Ethanolamide | 326.30535 | 0.9337509 | 44 | -3.843304672 | positive |
| Ornithine | 133.09767 | 11.897278 | 63 | -6.791161066 | positive |
| Palmitoyl sphingomyelin | 703.5754 | 5.7373878 | 98 | -5.443583223 | positive |
| Pantothenic Acid(VB5) | 220.11795 | 2.9909175 | 80 | -3.678264918 | positive |
| PC(30:0) | 706.53868 | 4.603938 | 100 | -4.198348667 | positive |
| PC(32:0) | 734.56998 | 4.5365688 | N/A | -4.59297075 | positive |
| 3a,7a-Dihydroxy-5b-cholestane | 405.37271 | 0.93241726 | 18 | -2.704127917 | positive |
| PC(32:1) | 732.55433 | 4.5171943 | 100 | -4.523575167 | positive |
| PC(32:2) | 730.53868 | 4.5194845 | 99 | -4.209416333 | positive |
| PC(34:1) | 760.58563 | 4.4183724 | 87 | -9.069563388 | positive |
| PC(34:2) | 758.56998 | 4.4184794 | 100 | -5.224520248 | positive |
| PC(34:3) | 756.55433 | 4.4208065 | 100 | -4.522600667 | positive |
| PC(35:2) | 772.58563 | 4.3681612 | 100 | -4.76273675 | positive |
| PC(35:5) | 766.53868 | 4.5346066 | 100 | -7.138307333 | positive |
| PC(36:2) | 786.59878 | 4.3506234 | 100 | -3.013471833 | positive |
| PC(36:3) | 784.58563 | 4.3260339 | 100 | -6.024376446 | positive |
| PC(36:4) | 782.56998 | 4.2438305 | 100 | -5.227026833 | positive |
| (25R)-5beta-cholestane-3alpha,7alpha,12alpha,26-tetrol | 437.36254 | 0.88856586 | N/A | -3.101727167 | positive |
| 3alpha,7alpha-Dihydroxy-5beta-cholestanate | 435.34689 | 0.94552678 | N/A | -3.191266583 | positive |
| PC(36:5) | 780.55433 | 4.256175 | 98 | -4.354871667 | positive |
| PC(38:5) | 808.58563 | 4.1652334 | 100 | -7.155614793 | positive |
| PC(38:6) | 806.56998 | 4.1871167 | 100 | -5.420216583 | positive |
| PC(38:7) | 804.55433 | 4.2046961 | 99 | -5.470367 | positive |
| PC(40:4) | 838.63258 | 4.1454879 | N/A | -7.589607167 | positive |
| PC(40:6) | 834.60128 | 4.1231836 | 100 | -5.463843417 | positive |
| PC(40:7) | 832.58563 | 4.0962322 | 100 | -4.59789375 | positive |
| PC(40:8) | 830.56998 | 4.1165067 | 100 | -4.666523417 | positive |
| PC(42:10) | 854.56998 | 3.973259 | 77 | -2.375544833 | positive |
| PC(42:9) | 856.58563 | 4.0449481 | 78 | -4.637703417 | positive |
| 3-Aminopropionaldehyde | 74.06004 | 9.9442679 | N/A | -1.104192869 | positive |
| PC(p32:0) | 718.57507 | 4.3710817 | 100 | -4.806583167 | positive |
| PC(p34:0) | 746.60637 | 4.3958038 | 99 | -5.495601917 | positive |
| PC(p34:1) | 744.59072 | 4.378589 | 100 | -4.603909 | positive |
| PC(p36:4) | 766.57507 | 4.0735774 | 100 | -5.059640583 | positive |
| PC(p36:5) | 764.55942 | 4.0930181 | 89 | -4.483462667 | positive |
| PC(p38:4) | 794.60637 | 4.0858472 | 90 | -5.258363417 | positive |
| PC(p38:6) | 790.57507 | 4.0094784 | 100 | -4.695017917 | positive |
| PE(35:0) | 734.56943 | 4.5365688 | N/A | -3.844239917 | positive |
| PE(36:4) | 740.52303 | 4.5684878 | 100 | -4.048156083 | positive |
| PE(38:6) | 764.52303 | 4.5463589 | 100 | -4.625614083 | positive |
| 3-Hydroxyanthranilic acid | 154.04987 | 2.2226258 | 54 | -3.574928033 | positive |
| PE(p38:4) | 752.55942 | 4.2887624 | N/A | -4.817863417 | positive |

| | | | | | |
|---|---|---|---|---|---|
| PE(p38:5) | 750.54377 | 4.2657404 | 69 | -4.761577667 | positive |
| PE(p38:6) | 748.52812 | 4.2927105 | 90 | -4.358439833 | positive |
| Phe-Phe | 313.15576 | 6.4284477 | 50 | -7.324960492 | positive |
| Phe-Trp | 352.16666 | 6.5351289 | 78 | -6.49764925 | positive |
| Phe-Tyr | 329.15068 | 0.87082583 | 10 | -6.80062675 | positive |
| Phytosphingosine (low) | 318.30027 | 2.5612747 | 96 | -3.76919377 | positive |
| Pivaloylcarnitine | 246.17053 | 5.3125795 | 85 | -5.572645868 | positive |
| Progesterone | 315.23186 | 0.89003735 | 49 | 9.161752333 | positive |
| Pro-Gln | 244.13028 | 10.04678 | 14 | -7.511052667 | positive |
| 3-Hydroxy-cis-5-tetradecenoylcarnitine | 386.29065 | 3.8323978 | 63 | -5.37165025 | positive |
| Pro-Ile | 229.15522 | 8.9564602 | 31 | -5.536418033 | positive |
| Propionylcarnitine | 218.13868 | 7.096336 | 75 | -3.254940738 | positive |
| Propionylglycine | 132.06607 | 12.852261 | 30 | -6.957487049 | positive |
| Pyridoxamine | 169.09715 | 4.0911548 | 11 | -3.173327787 | positive |
| Pyroglutamic acid | 130.04987 | 10.265 | 54 | -2.722833934 | positive |
| Spermidine | 146.16517 | 14.756954 | 86 | -3.107776803 | positive |
| Sphinganine | 302.30536 | 2.5189995 | 80 | -3.728057787 | positive |
| sphingosine-1-phosphate | 380.25604 | 7.4873149 | 93 | -3.370221 | positive |
| Succinylcarnitine | 262.12906 | 9.9500399 | 60 | -5.105832479 | positive |
| Taurine | 126.02247 | 9.3161688 | 30 | -6.720889344 | positive |
| 3-Hydroxyisovalerylcarnitine | 262.16492 | 8.7586956 | 75 | -3.118544333 | positive |
| Testosterone | 289.21621 | 4.0758249 | 20 | 6.986803884 | positive |
| Thr-Glu | 249.10975 | 11.227898 | 32 | -9.304610246 | positive |
| trans-Hexadec-2-enoyl carnitine | 398.32651 | 3.0088686 | 90 | -3.667272213 | positive |
| Trigonelline | 138.05548 | 8.9353933 | 67 | -6.459774016 | positive |
| Tyr-Tyr | 345.14559 | 2.7942675 | 10 | -6.354410744 | positive |
| Urocanic acid | 139.05073 | 3.250429 | 70 | -7.106029754 | positive |
| Xanthurenic acid | 206.04478 | 7.4318094 | N/A | -2.87783575 | positive |
| 3-hydroxy-N6,N6,N6-trimethyl-L-lysine | 205.15467 | 11.448132 | 20 | -3.107330917 | positive |
| 3-Methylhistidine | 170.0924 | 11.412276 | 83 | -3.070115738 | positive |
| 4-Aminopyridine | 95.06038 | 11.331716 | 100 | -2.135754344 | positive |
| 4-Androstenediol/Dihydrotestosterone | 291.23186 | 0.93138881 | N/A | -3.482428099 | positive |
| 4-Guanidinobutanoic acid | 146.0924 | 8.9857151 | 69 | -2.81681041 | positive |
| (2S,4R,5S)-Muscarine | 174.14886 | 4.6398087 | 30 | -2.818935082 | positive |
| 4-Pyridoxic Acid | 184.06096 | 2.5464599 | 40 | -5.55025314 | positive |
| 5-Hydroxyindoleacetic acid | 192.06552 | 8.1530484 | 62 | -2.990991967 | positive |
| 5-hydroxytryptophol | 178.08626 | 8.4780875 | N/A | -2.790644262 | positive |
| 5-Methylcytidine/2-O-Methylcytidine | 258.109 | 10.390118 | 3 | 0.466288852 | positive |
| 5'-Methylthioadenosine | 298.09736 | 2.4781825 | 39 | 8.2035295 | positive |
| 7a,12a-Dihydroxy-5b-cholestan-3-one | 419.35197 | 0.84819875 | N/A | -4.277459083 | positive |
| 7alpha,25-Dihydroxy-4-cholesten-3-one/Calcitriol | 417.33632 | 0.94733622 | N/A | -3.025659421 | positive |
| 7-Methylguanine | 166.07234 | 5.5704782 | 50 | -2.97757123 | positive |
| 9,12-Hexadecadienoylcarnitine | 396.31138 | 3.0567701 | 97 | -4.907436529 | positive |
| acetoacetate | 103.03897 | 9.3040576 | 55 | -2.402084918 | positive |
| (S)-1-Pyrroline-5-carboxylate/1-Pyrroline-2-carboxylic acid | 114.05495 | 2.4408657 | 12 | -2.671622951 | positive |
| Acetyl-N-formyl-5-methoxykynurenamine | 265.11828 | 6.3034092 | 21 | -3.434865164 | positive |
| AcylCarnitine(10:3) | 310.20129 | 4.2098685 | 90 | -4.28591686 | positive |
| AcylCarnitine(12:1) | 342.26389 | 3.4245378 | 100 | -3.859624426 | positive |
| AcylCarnitine(14:2) | 368.27954 | 3.2385927 | 100 | -3.858351901 | positive |
| AcylCarnitine(14:3) | 366.26389 | 3.3458123 | 100 | -3.737648264 | positive |
| AcylCarnitine(20:1) | 454.38909 | 2.7638258 | 43 | -3.333538167 | positive |
| AcylCarnitine(20:2) | 452.37344 | 2.7985424 | 79 | -3.121101333 | positive |
| AcylCarnitine(20:4) | 448.34214 | 2.8195011 | 100 | -3.654536116 | positive |
| AcylCarnitine(22:4) | 476.37344 | 2.7534302 | 90 | -1.99545025 | positive |
| Adenine | 136.0623 | 4.5680199 | 42 | -6.906075164 | positive |
| 17a-hydroxypregnenolone | 333.24242 | 0.89003735 | N/A | -0.848794083 | positive |
| Adenosine monophosphate | 348.07036 | 10.783793 | 60 | -3.489732049 | positive |
| Ala-Ala | 161.09316 | 6.6682166 | 21 | -9.665129836 | positive |
| Ala-Pro | 187.10881 | 8.779837 | 24 | -8.277095574 | positive |
| Alpha-Linolenic acid | 279.23186 | 0.93034192 | 100 | -3.423675574 | positive |
| Arg-Leu | 288.20411 | 10.72973 | 63 | -5.006923058 | positive |
| Arg-Phe | 322.18846 | 10.496573 | 67 | -6.019151795 | positive |
| Asp-Gly | 191.06734 | 11.521397 | 43 | -7.717788182 | positive |
| Asp-Val | 233.11429 | 9.9448407 | 30 | -4.634169672 | positive |
| Asymmetric dimethylarginine | 203.15025 | 10.734188 | 99 | -3.16983082 | positive |
| Benzenebutanoic acid | 165.09101 | 0.84670005 | 64 | -3.357374918 | positive |
| 1-aminocyclopropane-1-carboxylate | 102.05548 | 9.9560353 | 69 | -7.226763525 | positive |

| | | | | | |
|---|---|---|---|---|---|
| Benzylamine | 108.08078 | 1.0892529 | 31 | -2.783396066 | positive |
| Betaine | 118.08678 | 8.5658176 | 62 | -7.491518361 | positive |
| Betaine aldehyde | 102.09134 | 9.3051986 | 54 | -2.288614672 | positive |
| Bilirubin | 585.27076 | 0.89010453 | 93 | -4.336375833 | positive |
| Biliverdin | 583.25511 | 2.1409901 | 92 | -3.56115475 | positive |
| Butyrylcarnitine | 232.15433 | 5.9750547 | 70 | -3.832084463 | positive |
| Caproylcholine | 202.1807 | 5.7005589 | N/A | -6.448651475 | positive |
| Capryloylglycine | 202.14432 | 9.3080759 | N/A | -6.224372213 | positive |
| Cholesterol-H2O | 369.35155 | 0.87083588 | 100 | -3.825816475 | positive |
| Choline | 104.10699 | 5.0566115 | 73 | -2.552920164 | positive |
| 1H-Indole-4-carboxaldehyde | 146.06004 | 5.9212488 | 40 | -3.175469918 | positive |
| cis-5-Tetradecenoylcarnitine | 370.29521 | 3.1218933 | 100 | -3.890919918 | positive |
| Citrulline | 176.10349 | 10.526297 | 95 | -5.519906803 | positive |
| Creatine | 132.07728 | 9.7571538 | 81 | -6.947752131 | positive |
| Creatinine | 114.06671 | 4.9630581 | 75 | -7.17658123 | positive |
| Cytosine | 112.05106 | 5.3501439 | 56 | -6.874394508 | positive |
| D-(-)-Arabinose | 151.0601 | 4.1142982 | N/A | 5.346930328 | positive |
| D-(+)-Glucose | 181.07067 | 1.6398637 | 60 | 3.758669894 | positive |
| Decanoylcarnitine | 316.24824 | 3.6575105 | 85 | -3.926724959 | positive |
| Decenoylcarnitine(C10:1) | 314.23259 | 3.7777791 | 100 | -4.112471148 | positive |
| deoxycarnitine | 146.11808 | 9.2310298 | 60 | -6.79626959 | positive |
| 1-methyladenosine | 282.12021 | 8.9908877 | 29 | -4.763157213 | positive |
| Deoxycorticosterone | 331.22677 | 0.89003735 | N/A | -3.528709417 | positive |
| Dihydroxyacetone phosphate | 171.0053 | 14.75681 | 11 | -0.952697131 | positive |
| Dimethylglycine | 104.07061 | 9.309243 | 93 | -2.456636475 | positive |
| Dimethylurea | 89.07094 | 1.5545219 | 100 | -2.325482459 | positive |
| DL-Stearoylcarnitine | 428.37346 | 2.8312365 | 37 | -1.76526675 | positive |
| Docosenamide | 338.34174 | 0.88990881 | 100 | -4.241967541 | positive |
| Elaidic carnitine | 426.35781 | 2.8475976 | 100 | -3.8157725 | positive |
| Ergothioneine | 230.09577 | 9.5258131 | 82 | -3.546703917 | positive |
| Gamma Glutamylglutamic acid | 277.10358 | 7.6154492 | 14 | 7.656465164 | positive |
| Gln-Glu | 276.11956 | 11.785313 | 68 | -5.106211417 | positive |
| 1-Methylnicotinamide | 137.07094 | 6.7552267 | 69 | -2.827820082 | positive |
| Glycerophosphocholine | 258.1101 | 10.401866 | 79 | -3.795461148 | positive |
| Glycine | 76.03983 | 10.065974 | 50 | -7.863836803 | positive |
| Glycocholic acid | 466.31686 | 7.1744207 | 94 | -4.5920965 | positive |
| Guanidineacetic acid | 118.06163 | 9.9369216 | 47 | -0.064666333 | positive |
| Heptadecanoyl carnitine | 414.35781 | 2.9089729 | 40 | -3.275239504 | positive |
| Heptanoylcarnitine | 274.20183 | 5.971067 | 15 | -5.810186639 | positive |
| Hexacosanoyl carnitine | 540.49866 | 2.4932704 | N/A | -2.772705167 | positive |
| Hexanoylcarnitine | 260.18563 | 4.7867917 | 88 | -3.335831167 | positive |
| Hexylresorcinol | 195.13796 | 0.93173981 | N/A | -3.016450574 | positive |
| Hippuric acid | 180.06552 | 2.50535 | 23 | -2.795790574 | positive |
| (25R)-5beta-cholestane-3alpha,7alpha,12alpha,26-tetrol | 435.34798 | 13.01 | N/A | 0.343170248 | negative |
| 2-Aminobenzoic acid/p-Aminobenzoic acid | 136.0404 | 7.76 | NA | 1.450921885 | negative |
| FA(17:1)-H | 267.2328 | 10.14 | N/A | 0.969161885 | negative |
| FA(19:1)-H | 295.2644 | 11.31 | N/A | -0.169286721 | negative |
| FA(20:2)-H | 307.2643 | 10.86 | 38 | -0.069333223 | negative |
| FA(20:3)-H | 305.2483 | 10.26 | 84 | 0.929355902 | negative |
| FA(21:3)-H | 319.264 | 7.87 | N/A | 1.21736157 | negative |
| FA(22:1)-H | 337.3117 | 16.68 | N/A | -0.441865656 | negative |
| FA(22:2)-H | 335.2962 | 13.71 | N/A | -0.941513636 | negative |
| FA(22:4)-H | 331.2642 | 10.56 | 96 | 0.767742705 | negative |
| FA(22:5)-H | 329.2484 | 10.09 | 86 | 0.851150574 | negative |
| FA(24:4)-H | 359.2962 | 11.92 | N/A | -0.982036557 | negative |
| 2-Furoic acid | 111.00876 | 1.31 | N/A | 1.386237049 | negative |
| FA(24:5)-H | 357.2799 | 10.81 | N/A | 1.222357705 | negative |
| FA(24:6)-H | 355.2646 | 10.34 | N/A | -0.872253362 | negative |
| fumarate | 115.00316 | 1.38 | 94 | 5.709720246 | negative |
| galacturonic acid | 193.03538 | 1.49 | 93 | 1.036047787 | negative |
| Gluconic acid | 195.0505 | 1.5 | 92 | 3.758050984 | negative |
| Glu-Phe | 293.11265 | 3.31 | 83 | 7.063387438 | negative |
| Glutaric acid | 131.03498 | 1.42 | 96 | 1.721131393 | negative |
| Glyceraldehyde 3-phosphate | 168.99023 | 6.82 | NA | 1.993852377 | negative |
| Glycerate 2-phosphate | 184.98566 | 1.35 | 89 | -0.045352951 | negative |
| Glyceric acid | 105.01933 | 1.51 | 93 | 1.26903623 | negative |
| 2-hexadecenal | 237.22239 | 10.38 | N/A | 0.503916885 | negative |

| | | | | | |
|---|---|---|---|---|---|
| Glycerol | 91.04007 | 1.72 | N/A | 1.529935738 | negative |
| Glycerol 3-phosphate | 171.0064 | 1.63 | 92 | 1.94511918 | negative |
| Glycerophosphocholine | 256.09555 | 1.62 | N/A | 1.508262377 | negative |
| Glycochenodeoxycholic acid | 448.30685 | 7.78 | 94 | 0.045744098 | negative |
| Glycocholic acid | 464.30176 | 7.44 | 92 | 0.357869754 | negative |
| Glycyrrhetinic acid | 469.33233 | 8.27 | 82 | -5.688646393 | negative |
| Gly-Phe | 221.09207 | 5.8 | 97 | -1.775333956 | negative |
| Gly-Val | 173.09207 | 2.27 | 95 | 7.157294918 | negative |
| GUDCA | 448.30685 | 7.43 | 94 | 0.045744098 | negative |
| Heptadecanoic acid(17:0) | 269.24806 | 11.02 | N/A | 2.451319508 | negative |
| 2-hydroxy-4-(methylthio)butyric acid | 149.02727 | 2.3 | 83 | 4.813839587 | negative |
| Hexadecanedioic acid | 285.20713 | 6.81 | 86 | 1.116698852 | negative |
| Hexanoylglycine (predicted) | 172.09737 | 2.53 | NA | 4.905341393 | negative |
| Hexylresorcinol | 193.1234 | 7.54 | 61 | 1.132461557 | negative |
| Hippuric acid | 178.05042 | 5.76 | 96 | 3.128845372 | negative |
| Histidine | 154.06168 | 1.73 | 91 | 5.39885582 | negative |
| Hydrocinnamic acid | 149.0608 | 6.22 | 71 | 1.652262705 | negative |
| Hydroxycaprylic acid | 159.10267 | 6.49 | 77 | 1.738953607 | negative |
| Hydroxyphenyllactic acid | 181.05063 | 2.27 | 92 | 1.665757213 | negative |
| Hypoxanthine | 135.03123 | 2.27 | 83 | -7.827784016 | negative |
| Ile-Arg | 286.18736 | 5.91 | 91 | 4.259590617 | negative |
| 2-Hydroxyglutarate | 147.0299 | 1.38 | 94 | 1.192431393 | negative |
| Indole | 116.05057 | 6.02 | 86 | 1.514789262 | negative |
| indole-3-ethanol | 160.07679 | 2.26 | 17 | 2.023209381 | negative |
| Indole-5,6-quinone | 146.02475 | 7.14 | N/A | 2.197851721 | negative |
| Indoleacetaldehyde | 158.06114 | 6.04 | 72 | 1.482664262 | negative |
| Indoleacrylic acid | 186.05605 | 6.43 | 83 | 1.599956033 | negative |
| Indoxyl sulfate | 212.00178 | 6.38 | 81 | 2.310246612 | negative |
| Inosine | 267.07297 | 2.63 | 88 | -1.143420248 | negative |
| 2-Isopropylmalic acid/3-Propylmalate | 175.0612 | 5.91 | N/A | -5.258326967 | negative |
| Ketoleucine | 129.05571 | 5.63 | 28 | 1.215012213 | negative |
| 2-keto valeric acid | 115.04007 | 2.49 | 80 | 1.153678279 | negative |
| Kynurenic acid | 188.03532 | 5.85 | 92 | 1.888873554 | negative |
| lactate | 89.0239 | 1.59 | N/A | 6.391336281 | negative |
| L-alanine | 88.03988 | 1.32 | 59 | 6.791698361 | negative |
| L-Arginine | 173.1044 | 1.78 | N/A | 1.225454918 | negative |
| L-Asparagine | 131.04621 | 1.59 | 87 | 1.79327877 | negative |
| Lauric acid(12:0) | 199.1698 | 8.59 | N/A | 2.48011623 | negative |
| lauroylcarnitine | 342.26445 | 9.32 | N/A | 1.712378678 | negative |
| L-glutamic acid | 146.04536 | 1.47 | 89 | 5.25654918 | negative |
| L-Glutamine | 145.06186 | 1.6 | 93 | 2.146055246 | negative |
| L-Homocysteine | 134.02812 | 1.82 | N/A | 1.837039917 | negative |
| 2-Ketobutyric acid | 101.02442 | 1.64 | 73 | 1.569862459 | negative |
| Linoleic acid(18:2) | 279.23295 | 9.94 | 79 | -1.766820984 | negative |
| L-Isoleucine | 130.08683 | 2.41 | 70 | 4.503543033 | negative |
| Lithocholic acid | 375.29047 | 8.61 | 84 | 2.091611204 | negative |
| L-kynurenine | 207.07699 | 4.81 | 92 | -0.293676803 | negative |
| L-Leucine | 130.08683 | 2.55 | 70 | 4.503543033 | negative |
| L-lysine | 145.09825 | 1.72 | 63 | 1.887511967 | negative |
| L-methionine | 148.04325 | 2.3 | 60 | 5.607413279 | negative |
| L-Ornithine | 131.0826 | 1.65 | 81 | 1.613099836 | negative |
| L-Phenylalanine | 164.07118 | 5.12 | 98 | 3.656768361 | negative |
| L-proline | 114.05553 | 1.78 | 82 | 5.786023279 | negative |
| 2-Ketohexanoic acid | 129.05571 | 5.46 | 28 | 1.215012213 | negative |
| L-rhamnose | 163.0612 | 1.77 | 80 | 1.499540984 | negative |
| L-serine | 104.03479 | 1.6 | 82 | 7.103653607 | negative |
| L-Tryptophan | 203.0826 | 6.05 | 83 | -0.276757623 | negative |
| L-tryptophanamide | 202.09806 | 1.34 | NA | -8.533186356 | negative |
| L-Tyrosine | 180.06609 | 2.3 | 97 | 2.806520328 | negative |
| L-valine | 116.0717 | 1.84 | 87 | 1.341419426 | negative |
| LysoPA(16:0)-H | 409.23621 | 11.1 | 92 | -0.795704098 | negative |
| LysoPA(16:1)-H | 407.22055 | 10.27 | 83 | -0.299064959 | negative |
| LysoPC(16:0)+CH3COO | 554.34635 | 11.11 | 96 | 0.181618347 | negative |
| 2-Methoxyestradiol | 301.18092 | 7.98 | N/A | 0.844905 | negative |
| LysoPC(18:2)+CH3COO | 578.34635 | 10.57 | 95 | 1.050144754 | negative |
| LysoPC(20:3)+CH3COO | 604.362 | 10.92 | 97 | 0.346038689 | negative |
| LysoPC(20:4)+CH3COO | 602.34635 | 10.52 | 97 | 1.010794262 | negative |

| | | | | | |
|---|---|---|---|---|---|
| LysoPE(16:0)-H | 452.27827 | 10.82 | 95 | -0.380441803 | negative |
| LysoPE(16:0e)-H | 438.299 | 11.15 | N/A | 1.032053967 | negative |
| LysoPE(16:1)-H | 450.26262 | 10.1 | 99 | -5.169216777 | negative |
| LysoPE(17:1)-H | 464.27879 | 11.1 | 75 | -1.305285 | negative |
| LysoPE(18:0e)-H | 466.3303 | 11.54 | N/A | -0.024186116 | negative |
| LysoPE(18:0p)-H | 464.31465 | 9.78 | 69 | -1.148628595 | negative |
| LysoPE(18:1)-H | 478.29434 | 11 | 98 | -1.280200082 | negative |
| (2S,4R,5S)-Muscarine | 172.1343 | 6.7 | N/A | 1.709573607 | negative |
| 2-Phenylglycine | 150.05605 | 6.07 | N/A | 1.776424426 | negative |
| LysoPE(18:2)-H | 476.27827 | 10.38 | 96 | -0.022512131 | negative |
| LysoPE(20:3)-H | 502.29395 | 10.67 | 94 | 0.316859918 | negative |
| LysoPE(20:4)-H | 500.27827 | 10.34 | 94 | 0.508577869 | negative |
| LysoPE(20:5)-H | 498.26328 | 9.89 | 88 | -0.442492787 | negative |
| LysoPE(22:5)-H | 526.29421 | 10.48 | 95 | -0.245318361 | negative |
| LysoPE(22:6)-H | 524.27827 | 10.25 | 90 | -0.384971639 | negative |
| LysoPG(16:0)-H | 483.27332 | 9.56 | 98 | -0.568392149 | negative |
| LysoPG(18:1)-H | 509.28873 | 9.67 | 92 | -0.230110744 | negative |
| LysoPG(18:2)-H | 507.27311 | 9.34 | 96 | -0.268225372 | negative |
| LysoPI(16:0)-H | 571.28889 | 9.33 | 97 | 0.449401901 | negative |
| 2-Pyrocatechuic acid | 153.01933 | 6.49 | 96 | 1.83013082 | negative |
| LysoPI(17:0)-H | 585.30446 | 9.58 | 92 | 1.73041425 | negative |
| LysoPI(18:1)-H | 597.30454 | 9.44 | 93 | 0.757019835 | negative |
| LysoPI(18:2)-H | 595.28889 | 9.13 | 80 | 0.313501818 | negative |
| LysoPI(20:2)-H | 623.32019 | 9.58 | 94 | 0.798491885 | negative |
| LysoPI(20:3)-H | 621.30454 | 9.31 | 97 | 0.968904793 | negative |
| LysoPI(20:4)-H | 619.28889 | 9.15 | 99 | -0.606022645 | negative |
| LysoPI(22:4)-H | 647.32124 | 9.51 | 92 | -0.255003667 | negative |
| LysoPI(22:5)-H | 645.30579 | 9.25 | 84 | -0.752498182 | negative |
| LysoPI(22:6)-H | 643.28972 | 9.12 | 98 | -0.038788512 | negative |
| LysoPS(18:0)-H | 524.2994 | 9.8 | 96 | 0.431211721 | negative |
| 3,4-Dihydroxy-L-phenylalanine(Levodopa) | 196.06101 | 2.26 | 84 | 3.738959508 | negative |
| LysoPS(18:1)-H | 522.28375 | 9.43 | 94 | 0.368781818 | negative |
| LysoPS(20:4)-H | 544.2681 | 9.11 | 93 | 0.574982562 | negative |
| Lys-Tyr | 308.16048 | 7.02 | 51 | -3.231757167 | negative |
| Malic acid | 133.01425 | 1.38 | 96 | 1.504148115 | negative |
| m-Coumaric acid/2-Hydroxycinnamic acid | 163.04007 | 2.3 | 90 | 1.995785984 | negative |
| Met-Phe | 295.11109 | 6.55 | 67 | 2.30336124 | negative |
| Myristic acid(14:0) | 227.20111 | 9.46 | 51 | 1.999222787 | negative |
| Myristoleic acid(14:1) | 225.18546 | 8.85 | N/A | 3.42670623 | negative |
| N-Acetyl-DL-methionine | 190.05382 | 3.38 | 96 | 4.517386281 | negative |
| N-acetyl-D-tryptophan | 245.09264 | 5.77 | NA | 3.043987686 | negative |
| 3a,7a-Dihydroxy-5b-cholestane | 403.35815 | 9.1 | N/A | 1.074288333 | negative |
| N-Acetylglutamic acid | 188.05645 | 1.48 | 91 | 1.553035738 | negative |
| N-Acetylglutamine | 187.07243 | 1.31 | NA | 1.269461721 | negative |
| N-Acetylglycine | 116.03479 | 1.38 | 89 | 5.778334672 | negative |
| N-Acetyl-L-aspartic acid | 174.04027 | 1.46 | 93 | 4.523516475 | negative |
| N-Acetyl-L-phenylalanine | 206.08227 | 6.1 | NA | 1.258562893 | negative |
| N-Acetyl-L-tyrosine | 222.07718 | 2.3 | NA | 0.996238898 | negative |
| N-Acetylneuraminic acid | 308.09818 | 1.39 | NA | 1.757032727 | negative |
| NADP(nicotinamide adenine dinucleotide phosphate) | 742.06818 | 1.43 | N/A | 2.349470087 | negative |
| N-alpha-Acetyl-L-lysine(Acetyllysine) | 187.10882 | 1.78 | 89 | 1.412012479 | negative |
| N-Formylmethionine | 176.03817 | 2.57 | 97 | 4.325336393 | negative |
| 3alpha,7alpha-Dihydroxy-5beta-cholestanate | 433.33233 | 11.21 | N/A | 0.45891562 | negative |
| N-methyl-L-histidine | 168.07785 | 1.82 | N/A | 1.579874711 | negative |
| N-oleoyl taurine | 388.2527 | 9.68 | 89 | 0.873521901 | negative |
| N-palmitoyl taurine | 362.23705 | 9.57 | 94 | 0.784559091 | negative |
| Oleic acid(18:1) | 281.2486 | 10.66 | N/A | -1.703411885 | negative |
| Oxalic acid | 88.98748 | 1.38 | N/A | 8.188885738 | negative |
| Oxoadipic acid | 159.0299 | 1.63 | 83 | 1.939258852 | negative |
| Palmitic acid(16:0) | 255.23295 | 10.38 | 69 | -1.282198525 | negative |
| Palmitoleic acid(16:1) | 253.2173 | 9.66 | N/A | -0.756051885 | negative |
| Pantothenic acid | 218.1034 | 2.3 | 91 | 0.987488512 | negative |
| p-Cresol sulfate | 187.0065 | 6.72 | 98 | 1.762285 | negative |
| 3-dehydroshikimate | 171.02937 | 1.62 | NA | -9.077985246 | negative |
| Pelargonic acid(9:0) | 157.1234 | 7.42 | N/A | 1.255810738 | negative |
| Pentadecanoic acid(15:0) | 241.2173 | 9.91 | 12 | 1.03924541 | negative |
| PG(16:0_18:1)-H | 747.51953 | 11.46 | 98 | -0.086002869 | negative |

| | | | | | |
|---|---|---|---|---|---|
| PG(16:0_18:2)-H | 745.50334 | 11.3 | 96 | 1.354303636 | negative |
| Phe-Ile | 277.15576 | 6.89 | 92 | 0.704797059 | negative |
| Phenaceturic acid | 192.06607 | 6.03 | 87 | 4.342609417 | negative |
| Phenylacetic acid | 135.0446 | 2.29 | 76 | 5.832467459 | negative |
| Phe-Phe | 311.13902 | 7.08 | 97 | 4.901814833 | negative |
| Phe-Trp | 350.14991 | 7.12 | 92 | 1.764466942 | negative |
| Phosphoric acid | 96.96962 | 1.41 | N/A | 0.810010246 | negative |
| 3-hydroxy-3-methylglutarate | 161.04555 | 1.65 | NA | 2.107679016 | negative |
| PI(16:0_18:2)-H | 833.52021 | 11.19 | 99 | -0.189175537 | negative |
| PI(16:0_20:4)-H | 857.52056 | 11.14 | 99 | -0.826739587 | negative |
| PI(16:0_22:6)-H | 881.51998 | 11.09 | 97 | -0.011232727 | negative |
| PI(17:0_20:4)-H | 871.53556 | 11.25 | 97 | 0.034483636 | negative |
| PI(18:0_20:4)-H | 885.55207 | 11.35 | 98 | -3.374336116 | negative |
| PI(18:0_22:6)-H | 909.55008 | 11.28 | 96 | 1.476417273 | negative |
| PI(18:1_20:4)-H | 883.5352 | 11.17 | 97 | 0.57677595 | negative |
| Pipecolic acid | 128.0717 | 2.3 | 82 | 1.759203115 | negative |
| Pivalic acid(Trimethylacetic acid) | 101.0608 | 2.53 | 83 | 1.338043197 | negative |
| prasterone sulfate | 367.15792 | 7.71 | 90 | 0.882408595 | negative |
| 3-hydroxybenzyl alcohol | 123.04515 | 6.65 | 68 | 1.857072975 | negative |
| Pregnanediol | 319.26425 | 7.87 | N/A | 0.434310496 | negative |
| Purine | 119.03632 | 1.62 | 31 | -9.720739754 | negative |
| Pyroglutamic acid(5-oxo-D-proline) | 128.03531 | 1.63 | 79 | 1.945612705 | negative |
| Pyrophosphate | 176.93595 | 1.39 | 56 | 1.227499917 | negative |
| Pyruvic acid | 87.00877 | 1.66 | N/A | 1.486930902 | negative |
| Quercetin | 301.03537 | 1.44 | N/A | 1.526911818 | negative |
| Quinolinic acid | 166.01458 | 1.64 | NA | -6.39906959 | negative |
| Retinoic acid | 299.20165 | 9.2 | N/A | 0.22372418 | negative |
| Ribothymidine | 257.07791 | 3.74 | 74 | -6.606371721 | negative |
| S-(-)-Carbidopa | 225.08807 | 2.29 | NA | -7.394602167 | negative |
| 3-Hydroxybutyric acid | 103.03955 | 1.67 | NA | 6.567094262 | negative |
| Salicylic acid | 137.02441 | 6 | 89 | 1.791298115 | negative |
| Salicyluric acid | 194.04588 | 2.3 | 55 | 0.938856639 | negative |
| Sebacic acid | 201.11323 | 6.6 | 82 | 1.498628361 | negative |
| Shikimic acid | 173.04503 | 2.29 | NA | 4.193158033 | negative |
| sphingosine-1-phosphate | 378.24148 | 8.99 | N/A | 0.55973377 | negative |
| Stearic acid(18:0) | 283.26425 | 12.29 | N/A | -1.31328459 | negative |
| Stearidonic acid(18:4) | 275.20111 | 9.13 | N/A | 2.409042951 | negative |
| succinate | 117.01881 | 1.38 | 96 | 5.612561721 | negative |
| succinate semialdehyde | 101.02441 | 1.64 | 73 | 1.668852131 | negative |
| Sulfate | 96.9601 | 1.37 | 22 | 1.219814098 | negative |
| 3-Hydroxycapric acid | 187.13397 | 7.41 | 87 | 1.451890492 | negative |
| Taurine | 124.00738 | 1.6 | 82 | 1.745364426 | negative |
| Tauro-b-muricholic acid/Tauro-a-muricholic acid | 514.2844 | 7.19 | 81 | 0.403368403 | negative |
| Taurochenodeoxycholic acid | 498.28948 | 7.89 | 77 | 0.699777049 | negative |
| Taurocholic acid | 514.2844 | 7.55 | 81 | 0.403368403 | negative |
| Taurolithocholic acid | 482.29457 | 7.16 | N/A | 2.500452297 | negative |
| Tauro-w-muricholic acid | 514.2844 | 7.34 | 81 | 0.403368403 | negative |
| Tetradecanedioic acid | 257.17583 | 6.36 | 83 | 1.424079918 | negative |
| Thr-Ala | 189.08808 | 1.36 | 86 | 1.1813285 | negative |
| Threonic acid | 135.0299 | 1.47 | 78 | 2.021808033 | negative |
| trans-aconitate | 173.00916 | 1.33 | 91 | 1.182850656 | negative |
| (RS)-Mevalonic acid | 147.06628 | 1.83 | N/A | 1.822012951 | negative |
| 3-Hydroxyisovaleric acid | 117.05571 | 2.26 | 95 | 1.465117377 | negative |
| Traumatic acid | 227.12888 | 5.83 | 75 | 1.769221721 | negative |
| Tridecylic acid(C13:0) | 213.18546 | 8.9 | N/A | 4.04962918 | negative |
| Tryptamine | 159.09277 | 6.04 | N/A | 1.567284545 | negative |
| Tryptophanol | 160.07679 | 1.8 | 17 | 2.023209381 | negative |
| TUDCA | 498.28948 | 7.52 | 77 | 0.699777049 | negative |
| Undecanoic acid(11:0) | 185.1547 | 8.18 | N/A | 1.41912877 | negative |
| Uracil | 111.01948 | 2.44 | N/A | 6.313725207 | negative |
| Uric acid | 167.02054 | 1.69 | 94 | 4.142331967 | negative |
| Uridine | 243.06174 | 2.43 | 83 | 2.882872541 | negative |
| Urocanic acid | 137.03565 | 2.3 | 83 | 1.299932459 | negative |
| 3-Indoleacetic acid | 174.05605 | 5.88 | 85 | 1.367566942 | negative |
| Ursodeoxycholic acid | 391.28538 | 7.41 | 98 | 0.455502705 | negative |
| Valproic acid(8:0) | 143.10775 | 7.02 | N/A | 1.696825164 | negative |
| w-MCA | 407.2803 | 7.2 | 77 | 0.431651885 | negative |

| | | | | | |
|---|---|---|---|---|---|
| Xanthine | 151.02615 | 1.72 | 89 | 1.805777541 | negative |
| 3-Indolepropionic acid | 188.0717 | 6.37 | 87 | 1.172285455 | negative |
| 3-methoxy-L-tyrosine | 210.07772 | 2.68 | 93 | -2.60260913 | negative |
| 3-Methyl-2-oxovaleric acid | 129.05571 | 4.95 | 28 | 1.215012213 | negative |
| 3-Phosphonoalanine | 168.00673 | 1.43 | N/A | -0.302979672 | negative |
| 4-acetamidobutanoate | 144.06662 | 1.81 | 88 | 1.805749098 | negative |
| 4-aminobutanoate(GABA) | 102.05605 | 1.48 | 85 | 1.595971967 | negative |
| 4-Aminopyridine | 93.04582 | 1.78 | N/A | 0.49081877 | negative |
| 4-Ethylphenol | 121.06588 | 7.18 | N/A | 1.330609224 | negative |
| 12(S)-HPETE | 335.22278 | 7.2 | N/A | 1.047554711 | negative |
| 4-ethylphenylsulfate(predicted) | 201.0227 | 7.16 | 77 | 1.249242083 | negative |
| 4-Hydroxybenzenesulfonic acid | 172.9914 | 6.17 | 60 | -0.468172975 | negative |
| 4-hydroxybenzoate | 137.02441 | 6.01 | 93 | 1.791298115 | negative |
| 4-hydroxyphenylacetate | 151.04007 | 2.31 | 92 | 1.995357541 | negative |
| 4-Pyridoxic acid | 182.04536 | 5.7 | 96 | 3.900326557 | negative |
| 4-Vinylphenol sulfate | 199.00705 | 7.06 | 92 | 1.253100579 | negative |
| 5,6-Epoxy-8,11,14-eicosatrienoic acid/5-HETE | 319.22787 | 8.68 | N/A | 0.507776475 | negative |
| 5-Acetamidovalerate | 158.08227 | 1.83 | N/A | 1.628334098 | negative |
| 5-Hydroxyindole-3-acetic acid | 190.05042 | 4.83 | 75 | 4.661130248 | negative |
| 5-Hydroxy-L-tryptophan | 219.07751 | 2.59 | NA | 1.057910744 | negative |
| 13(S)-HPOT | 309.20713 | 6.94 | 82 | 1.13519041 | negative |
| 5-hydroxytryptophol | 176.0717 | 6.32 | N/A | 1.45343 | negative |
| 7alpha,25-Dihydroxy-4-cholesten-3-one/Calcitriol | 415.32177 | 9.49 | N/A | 0.455547787 | negative |
| Acetyl-N-formyl-5-methoxykynurenamine | 263.10373 | 6 | N/A | 0.084373967 | negative |
| Adenosine | 266.08895 | 1.51 | 55 | -1.385129835 | negative |
| Adenylsuccinic acid | 462.06677 | 1.48 | N/A | 3.524132609 | negative |
| Adipic acid | 145.05063 | 1.82 | NA | 2.085216967 | negative |
| Adonitol | 151.0612 | 2.3 | NA | 10.86941754 | negative |
| alpha-Ketoglutaric acid(Oxoglutaric acid) | 145.01425 | 1.38 | 82 | 2.679438689 | negative |
| Alpha-Linolenic acid(18:3) | 277.2173 | 9.47 | 86 | -0.895427131 | negative |
| a-MCA/b-MCA | 407.2803 | 7.04 | 77 | 0.431651885 | negative |
| 16(17)-EpDPE | 343.22787 | 8.65 | 81 | 2.515448288 | negative |
| Arabinonic acid | 165.03991 | 1.46 | N/A | 4.285450656 | negative |
| Arachidic acid(20:0) | 311.29555 | 17.59 | 91 | 1.046994508 | negative |
| Arachidonic acid(20:4) | 303.23295 | 9.86 | 87 | -0.722866311 | negative |
| Ascorbic acid | 175.02429 | 1.43 | 91 | 4.110318607 | negative |
| Asymmetric dimethylarginine | 201.1357 | 2.3 | N/A | 1.632337934 | negative |
| Azelaic acid | 187.09758 | 1.79 | 84 | 1.848128689 | negative |
| Benzenebutanoic acid | 163.07645 | 7.3 | NA | 2.026507049 | negative |
| beta-alanine | 88.03988 | 1.61 | 59 | 6.791698361 | negative |
| Bilirubin | 583.25621 | 7.43 | N/A | 0.679676446 | negative |
| Biliverdin | 581.24056 | 7.2 | 96 | 1.179791405 | negative |
| 1-Hydroxypyrene | 219.08154 | 2.96 | NA | -0.067706111 | negative |
| caffeate | 179.03498 | 3.25 | NA | 2.20767541 | negative |
| Calcidiol/7a-Hydroxy-cholestene-3-one | 399.32685 | 8.67 | N/A | 0.05140541 | negative |
| Capric acid(10:0) | 171.1385 | 7.79 | N/A | 4.844778934 | negative |
| Chenodeoxycholic acid | 391.28538 | 7.81 | 98 | 0.455502705 | negative |
| cholesterol sulfate | 465.3044 | 10.67 | 83 | -0.217543852 | negative |
| Cholic acid | 407.2803 | 7.43 | 77 | 0.431651885 | negative |
| Cinnamic acid | 147.0446 | 4.74 | NA | 2.87707418 | negative |
| Cinnamoylglycine | 204.06662 | 6.04 | NA | 0.931595082 | negative |
| Citraconic acid | 129.01933 | 1.31 | N/A | 1.602497541 | negative |
| citrate | 191.01973 | 1.37 | 95 | 1.161487869 | negative |
| 1-methyladenosine | 280.10461 | 1.46 | N/A | -1.97819287 | negative |
| Citrulline | 174.08789 | 1.66 | 91 | 4.603778115 | negative |
| cortisol | 361.20205 | 7.96 | 87 | -0.6786775 | negative |
| Creatine | 130.0622 | 1.71 | 87 | 1.439783884 | negative |
| creatinine | 112.05111 | 1.79 | 90 | 6.370200082 | negative |
| Cytidine | 242.07824 | 1.64 | 30 | 8.652255 | negative |
| D-(+)-Glucose | 179.05611 | 1.65 | 80 | 1.677504754 | negative |
| Deoxyadenosine | 250.09404 | 1.47 | NA | -1.767143333 | negative |
| Deoxycholic acid | 391.28538 | 7.9 | 98 | 0.455502705 | negative |
| D-glucono-1,5-lactone | 177.04046 | 1.63 | 95 | 1.813087213 | negative |
| D-Glucuronic acid | 193.03538 | 1.5 | 75 | 1.036047787 | negative |
| 2,5-Dihydroxyphenylacetic acid | 167.03443 | 3.16 | 88 | 4.619712787 | negative |
| D-Glutamylglycine | 203.06734 | 1.5 | 87 | 1.590475984 | negative |
| dIDP | 411.01124 | 1.42 | N/A | 3.882227411 | negative |

| | | | | | |
|---|---|---|---|---|---|
| dihydroxymandelic acid-1 | 183.02938 | 2.31 | NA | 1.232488264 | negative |
| dihydroxymandelic acid-2 | 183.02938 | 2.75 | NA | 1.232488264 | negative |
| D-lactose | 341.10894 | 1.82 | 92 | 0.477393361 | negative |
| Docosahexaenoic acid(22:6) | 327.23295 | 9.7 | 89 | -0.077423443 | negative |
| Dulcitol | 181.07176 | 1.66 | 88 | -7.552688852 | negative |
| Eicosanoic acid(20:1) | 309.27936 | 12.91 | N/A | 2.193411074 | negative |
| Eicosapentaenoic acid(20:5) | 301.2173 | 9.44 | 81 | 0.374246803 | negative |
| erythritol | 121.05011 | 1.68 | 33 | 6.105231557 | negative |

**Table S3: Differential metabolites between R and NR patients.**

| feature | metadata | value | coef | pval | qval | mean_NR | mean_R | median_NR | median_R |
|---|---|---|---|---|---|---|---|---|---|
| Beta-Alanine | Response | R | 0.448892711 | 0.0014139 | 0.06413238 | -8.522238811 | -8.116432916 | -8.457208783 | -8.131872993 |
| Creatine | Response | R | 0.522595122 | 0.001051347 | 0.06413238 | -10.10963036 | -9.554208319 | -9.992882755 | -9.444527482 |
| L-Glutamine | Response | R | 0.329876726 | 0.001113002 | 0.06413238 | -6.962771305 | -6.660330196 | -6.971517998 | -6.743466808 |
| Sphingosine 1-phosphate | Response | R | 0.330838905 | 0.000916236 | 0.06413238 | -8.270194548 | -7.975492104 | -8.318958738 | -7.960541066 |
| Asymmetric dimethylarginine | Response | R | 0.294371766 | 0.00243597 | 0.068289717 | -10.91213943 | -10.65258463 | -10.90906539 | -10.62765429 |
| L-Histidine | Response | R | 0.358132959 | 0.002683119 | 0.068289717 | -7.205364461 | -6.857546649 | -7.235862333 | -6.924708148 |
| L-Asparagine | Response | R | 0.430911712 | 0.001711073 | 0.068289717 | -10.10725465 | -9.702170827 | -10.2353378 | -9.725885339 |
| L-Lysine | Response | R | 0.339577148 | 0.002537755 | 0.068289717 | -8.072312355 | -7.763055873 | -7.993409719 | -7.743892377 |
| LysoPE(18:2) | Response | R | 0.489650165 | 0.002466148 | 0.068289717 | -6.720094017 | -6.297466341 | -6.757419379 | -6.470593537 |
| N-Acetyl-L-phenylalanine | Response | R | 0.494872938 | 0.002364799 | 0.068289717 | -12.55705951 | -12.13014216 | -12.49272285 | -12.16522075 |
| Valproic acid | Response | R | 0.452516959 | 0.002011042 | 0.068289717 | -11.21699775 | -10.78175017 | -11.09710176 | -10.83470282 |
| L-Methionine | Response | R | 0.383549436 | 0.003318751 | 0.071171105 | -8.51329257 | -8.169734291 | -8.504088198 | -8.261349462 |
| L-Serine | Response | R | 0.356865541 | 0.003372676 | 0.071171105 | -9.260356452 | -8.952269225 | -9.311121353 | -9.029439214 |
| Ribitol | Response | R | 0.286916042 | 0.005294101 | 0.081734879 | -8.508527012 | -8.25493091 | -8.501600139 | -8.272872824 |
| Gluconolactone | Response | R | 0.296127085 | 0.006304425 | 0.081734879 | -9.544057259 | -9.294534061 | -9.555470559 | -9.416002749 |
| Glyceraldehyde 3-phosphate | Response | R | 0.278650072 | 0.006429326 | 0.081734879 | -10.59632425 | -10.35762041 | -10.71181103 | -10.44153753 |
| L-Tyrosine | Response | R | 0.335444574 | 0.006653596 | 0.081734879 | -6.7452422 | -6.444166101 | -6.765462491 | -6.54981803 |
| LysoPA(16:0/0:0) | Response | R | 0.252186362 | 0.00552952 | 0.081734879 | -4.745457532 | -4.520005431 | -4.740112394 | -4.557556867 |
| LysoPE(17:1) | Response | R | 0.245749105 | 0.00665077 | 0.081734879 | -6.848647046 | -6.632629782 | -6.853505319 | -6.669654703 |
| LysoPE(18:1) | Response | R | 0.400923732 | 0.005406217 | 0.081734879 | -7.354444931 | -7.01071384 | -7.471370504 | -7.145892081 |
| LysoPG(18:2) | Response | R | 0.293607611 | 0.005969703 | 0.081734879 | -10.5651542 | -10.3183094 | -10.69209354 | -10.41483812 |
| N-Acetyl-L-methionine | Response | R | 0.495797809 | 0.004640464 | 0.081734879 | -11.65418897 | -11.25050009 | -11.69114745 | -11.10868422 |
| Pyroglutamic acid | Response | R | 0.276261507 | 0.006468916 | 0.081734879 | -7.511285533 | -7.28421201 | -7.62855609 | -7.334739624 |
| Threoninyl-Alanine | Response | R | 0.318689949 | 0.005977989 | 0.081734879 | -9.218121403 | -8.944213679 | -9.195484037 | -8.929600584 |
| Tryptophanamide | Response | R | 0.331710076 | 0.007004187 | 0.082482141 | -8.235963908 | -7.948500584 | -8.200950811 | -7.867430164 |
| (+/-)-2-Hydroxy-4-(methylthio)butanoic acid | Response | R | 0.489385152 | 0.008589034 | 0.083025978 | -12.17188995 | -11.7891042 | -12.09607813 | -11.95261393 |
| 4-Ethylphenol | Response | R | 0.799853318 | 0.008892428 | 0.083025978 | -12.57837129 | -11.83250016 | -12.90384059 | -12.05183997 |
| Cinnamoylglycine | Response | R | 0.421657093 | 0.00804294 | 0.083025978 | -8.602234851 | -8.273233345 | -8.680604578 | -8.212399892 |
| Citrulline | Response | R | 0.35773184 | 0.007872211 | 0.083025978 | -9.597836494 | -9.294149898 | -9.675412407 | -9.369870301 |
| Dodecanoylcarnitine | Response | R | 0.240558208 | 0.007782651 | 0.083025978 | -10.77789391 | -10.57122019 | -10.79103604 | -10.64016009 |
| L-Proline | Response | R | 0.395397255 | 0.007548458 | 0.083025978 | -6.804399995 | -6.473632999 | -6.94495096 | -6.564824018 |
| LysoPC(16:0) | Response | R | 0.244428081 | 0.00781195 | 0.083025978 | -8.842450181 | -8.629831028 | -8.780640736 | -8.664696405 |
| LysoPE(20:3) | Response | R | 0.382845387 | 0.009636336 | 0.083025978 | -9.459124484 | -9.098155884 | -9.527623964 | -9.114949251 |
| LysoPG(18:1) | Response | R | 0.234503307 | 0.009681103 | 0.083025978 | -10.03851186 | -9.849803442 | -9.979549329 | -9.900030598 |
| 2-Hydroxycinnamic acid | Response | R | 0.312799388 | 0.009596357 | 0.083025978 | -10.51386971 | -10.2467852 | -10.55587322 | -10.26152758 |
| N-Formyl-L-methionine | Response | R | 0.277950074 | 0.009006109 | 0.083025978 | -10.33399207 | -10.11295847 | -10.37193916 | -10.14185075 |
| Oxoadipic acid | Response | R | 0.297982657 | 0.009559845 | 0.083025978 | -9.865237045 | -9.609254552 | -9.957261128 | -9.730630488 |
| Phosphoric acid | Response | R | 0.262084382 | 0.009208545 | 0.083025978 | -4.99699367 | -4.751827387 | -4.985202116 | -4.807035965 |
| Purine | Response | R | 0.265470028 | 0.007644356 | 0.083025978 | -8.390430465 | -8.181860268 | -8.403970648 | -8.220417285 |
| Ribothymidine | Response | R | 0.270429474 | 0.008210326 | 0.083025978 | -11.46256641 | -11.23064002 | -11.37588701 | -11.27114611 |
| L-Leucine | Response | R | 0.316815889 | 0.010081851 | 0.084623198 | -5.764494121 | -5.486298866 | -5.706790249 | -5.506938834 |
| Galactitol | Response | R | 0.761869339 | 0.010199766 | 0.084711739 | -10.40966021 | -9.707476605 | -10.62669535 | -9.957250964 |
| Glucose | Response | R | 0.274892514 | 0.010425814 | 0.085687163 | -6.67492284 | -6.445904804 | -6.746840394 | -6.570825088 |
| 3-Hydroxymethylglutaric acid | Response | R | 0.277780561 | 0.010956298 | 0.085939947 | -7.7704851 | -7.547939714 | -7.869920824 | -7.641733442 |
| 9E-Heptadecenoic acid | Response | R | -0.250854628 | 0.010770116 | 0.085939947 | -6.323394199 | -6.53425575 | -6.231363161 | -6.460469275 |
| Gluconic acid | Response | R | 0.401180194 | 0.011110107 | 0.085939947 | -8.725782937 | -8.332897184 | -8.805705137 | -8.493432359 |
| L-Dopa | Response | R | 0.323530598 | 0.0117019 | 0.088776917 | -10.81744539 | -10.51973164 | -10.89479956 | -10.64152758 |
| 2-Ketobutyric acid | Response | R | 0.289030798 | 0.014075419 | 0.089046636 | -7.860899784 | -7.609593119 | -7.831040017 | -7.704842706 |
| 4-Aminobutanoate | Response | R | 0.374116557 | 0.013809695 | 0.089046636 | -9.743314724 | -9.422780197 | -9.779929352 | -9.434728534 |
| 4-Ethylphenylsulfate | Response | R | 1.326939411 | 0.013808021 | 0.089046636 | -10.70002462 | -9.462613162 | -10.7094566 | -9.377152718 |
| Capric acid | Response | R | 0.217417616 | 0.014368447 | 0.089046636 | -8.005360196 | -7.797232243 | -8.047000103 | -7.834284124 |
| Glyceric acid | Response | R | 0.285214677 | 0.013247699 | 0.089046636 | -9.151649493 | -8.874449798 | -9.179735599 | -8.900174997 |
| Indoleacetaldehyde | Response | R | 0.379242268 | 0.01295142 | 0.089046636 | -11.36769321 | -11.06950205 | -11.48868871 | -11.0604 |
| L-Glutamic acid | Response | R | 0.360974322 | 0.013263162 | 0.089046636 | -7.142476417 | -6.836329734 | -7.168718258 | -6.838763842 |
| L-Phenylalanine | Response | R | 0.305870084 | 0.012520996 | 0.089046636 | -6.826760953 | -6.562712458 | -6.838772115 | -6.572393679 |
| L-Valine | Response | R | 0.301826446 | 0.014287682 | 0.089046636 | -6.081677152 | -5.815352763 | -6.041957222 | -5.837606075 |
| LysoPE(16:0e) | Response | R | 0.47416151 | 0.01332334 | 0.089046636 | -10.60328958 | -10.09411262 | -10.62258388 | -10.03416238 |
| LysoPG(16:0) | Response | R | 0.332790758 | 0.011886155 | 0.089046636 | -11.33316469 | -11.04907527 | -11.38692935 | -11.0217658 |
| Phenylacetylglycine | Response | R | 0.379559606 | 0.01317576 | 0.089046636 | -12.17040076 | -11.85660722 | -12.21704391 | -11.93418594 |
| Succinic acid semialdehyde | Response | R | 0.289030798 | 0.014075419 | 0.089046636 | -7.860899784 | -7.609593119 | -7.831040017 | -7.704842706 |
| Eicosenoic acid | Response | R | -1.185172085 | 0.014717898 | 0.089090962 | -6.034189723 | -6.944904074 | -6.021017316 | -6.203244131 |
| Acetylglycine | Response | R | 0.282826188 | 0.015017868 | 0.089090962 | -8.087151594 | -7.842593529 | -8.133608886 | -7.794930265 |
| Palmitoleic acid | Response | R | -0.300403397 | 0.01483534 | 0.089090962 | -4.048809965 | -4.303509324 | -4.104726751 | -4.187946286 |
| (2S,4R,5S)-Muscarine | Response | R | 0.299412703 | 0.01587412 | 0.091577097 | -11.25666963 | -10.95788127 | -11.26927198 | -11.017052 |
| Mevalonic acid | Response | R | 0.437574627 | 0.016133354 | 0.091577097 | -11.49900394 | -11.09862077 | -11.36864835 | -11.15773857 |
| Caffeate | Response | R | 0.396680647 | 0.01770629 | 0.097809357 | -11.09185494 | -10.77185003 | -11.19091475 | -10.70873497 |
| 2-Hydroxyglutarate | Response | R | 0.263576843 | 0.018027253 | 0.098334783 | -9.013861664 | -8.759217689 | -8.994866237 | -8.838260564 |
| LysoPE(20:4) | Response | R | 0.259108358 | 0.018071665 | 0.098334783 | -6.698516673 | -6.483260579 | -6.65108143 | -6.55828978 |
| Cholesterol sulfate | Response | R | 0.348792464 | 0.019199858 | 0.100426295 | -7.187872066 | -6.960927459 | -7.251648346 | -7.030578411 |
| Lysyl-Tyrosine | Response | R | 0.314288896 | 0.019041208 | 0.100426295 | -13.18279266 | -12.91163014 | -13.12343757 | -12.98452939 |
| 10Z-Nonadecenoic acid | Response | R | -0.21689145 | 0.019452226 | 0.100972408 | -7.320688598 | -7.513168327 | -7.358499535 | -7.414173348 |
| 12(S)-HPETE | Response | R | 0.349887572 | 0.020004711 | 0.102491671 | -10.89876994 | -10.60929532 | -10.82144439 | -10.6794134 |
| Threonic acid | Response | R | 0.297755597 | 0.02125887 | 0.108214504 | -7.731181123 | -7.420264143 | -7.741088784 | -7.450518452 |
| N-Acetyl-L-aspartic acid | Response | R | 0.230449058 | 0.025475082 | 0.119641905 | -8.994826056 | -8.795432357 | -8.978325399 | -8.879138686 |
| Pyruvic acid | Response | R | 0.283600229 | 0.025002815 | 0.119641905 | -7.217069497 | -6.971323969 | -7.236655775 | -6.982982293 |
| Homocysteine | Response | R | 0.360378748 | 0.026239704 | 0.121070913 | -10.95190987 | -10.64061942 | -11.02019066 | -10.72378636 |
| 5-Hydroxyindoleacetic acid | Response | R | 0.393355374 | 0.026605321 | 0.122044177 | -11.79955705 | -11.43297525 | -11.73368473 | -11.44150748 |
| 3-Hydroxybutyric acid | Response | R | -0.26325274 | 0.027018282 | 0.123222108 | -6.105711906 | -6.400834417 | -6.207106376 | -6.361470649 |
| L-Alanine | Response | R | 0.286021514 | 0.027306823 | 0.123239251 | -8.067326726 | -7.818429675 | -8.017313865 | -7.760325389 |
| PG(16:0_18:2) | Response | R | 0.438706629 | 0.02749305 | 0.123239251 | -11.65954338 | -11.28489396 | -11.64822641 | -11.29620159 |
| Glycerol | Response | R | 0.169185032 | 0.028070971 | 0.123353314 | -8.741939883 | -8.586410259 | -8.715936215 | -8.617111818 |
| Hydroxyphenyllactic acid | Response | R | 0.279709889 | 0.028701301 | 0.123353314 | -8.509100904 | -8.316431267 | -8.615659068 | -8.261985997 |
| Dehydroepiandrosterone sulfate | Response | R | 0.473277694 | 0.030511512 | 0.124090636 | -6.724189458 | -6.316435195 | -6.700641992 | -6.293455705 |
| Uracil | Response | R | 0.220428473 | 0.031301909 | 0.124733363 | -9.625385952 | -9.43482898 | -9.626523966 | -9.483136232 |
| L-Arginine | Response | R | 0.30522666 | 0.031864372 | 0.126336631 | -7.695912858 | -7.420516296 | -7.708464836 | -7.461591153 |
| LysoPE(22:5) | Response | R | 0.235870868 | 0.033730098 | 0.129819743 | -9.66942236 | -9.472888096 | -9.67173304 | -9.489954652 |
| Malic acid | Response | R | 0.233080936 | 0.033667404 | 0.129819743 | -7.79291684 | -7.599508941 | -7.875941725 | -7.564143554 |
| Eicosadienoic acid | Response | R | -0.13089756 | 0.035107495 | 0.131278737 | -5.333985234 | -5.450121235 | -5.316049407 | -5.414752357 |
| Uridine | Response | R | 0.251851222 | 0.037827719 | 0.137539494 | -8.2549723 | -8.013231222 | -8.231405473 | -8.014361455 |
| Kynurenic acid | Response | R | 0.311589549 | 0.0380457 | 0.13769751 | -11.72186157 | -11.44728555 | -11.7193308 | -11.48003696 |
| 3-Hydroxyisovaleric acid | Response | R | 0.310886178 | 0.038625878 | 0.138526444 | -7.813467988 | -7.567414394 | -7.801304984 | -7.491113104 |
| 3-Methoxytyrosine | Response | R | 0.244057769 | 0.038541466 | 0.138526444 | -12.55241666 | -12.34694875 | -12.55129184 | -12.38159618 |
| Indole | Response | R | 0.27860184 | 0.039188404 | 0.138653143 | -8.891662279 | -8.629536824 | -8.743982746 | -8.639834771 |
| Pelargonic acid | Response | R | 0.248425079 | 0.039187457 | 0.138653143 | -7.958772638 | -7.737397495 | -7.993823925 | -7.81521231 |
| 2-Ketohexanoic acid | Response | R | 0.401066046 | 0.041168403 | 0.143725089 | -6.874203495 | -6.454807566 | -6.495754044 | -6.52883687 |
| 4-Acetamidobutanoic acid | Response | R | 0.276126951 | 0.041152602 | 0.143725089 | -10.8035147 | -10.55501467 | -10.83271467 | -10.56703971 |
| Sulfate | Response | R | 0.249949405 | 0.041492014 | 0.144216737 | -5.473005463 | -5.25656479 | -5.458159031 | -5.367410472 |
| Indoleacetic acid | Response | R | 0.39664488 | 0.042211803 | 0.146075053 | -9.904830693 | -9.51505492 | -9.989175718 | -9.558797485 |

| | | | | | | | | | |
|---|---|---|---|---|---|---|---|---|---|
| 4-Vinylphenol sulfate | Response | R | 0.518588472 | 0.043234388 | 0.146245696 | -9.18571134 | -8.773331885 | -9.260135897 | -8.731728548 |
| Myristoleic acid | Response | R | -0.247590377 | 0.043255296 | 0.146245696 | -7.312498947 | -7.510276169 | -7.325101522 | -7.373479051 |
| N-Acetylglutamine | Response | R | 0.597586083 | 0.043367953 | 0.146245696 | -7.074497168 | -6.594300274 | -6.773845461 | -6.473533003 |
| Fumaric acid | Response | R | 0.229127919 | 0.04494057 | 0.147129085 | -10.09422613 | -9.899644566 | -10.21546686 | -9.96017164 |
| L-Tryptophan | Response | R | 0.269187268 | 0.044717662 | 0.147129085 | -6.260813756 | -6.005191376 | -6.14485271 | -6.029731995 |
| LysoPC(18:2) | Response | R | 0.375339735 | 0.044824771 | 0.147129085 | -9.906577539 | -9.577090155 | -9.917501425 | -9.645165796 |
| Glycyl-Valine | Response | R | 0.606333988 | 0.046919415 | 0.149876189 | -12.36759992 | -11.87211592 | -12.43091234 | -12.1775095 |
| Docosapentaenoic acid (22n-6) | Response | R | -0.159824464 | 0.047899161 | 0.15177686 | -6.288793749 | -6.403214988 | -6.276112781 | -6.359801513 |
| Oleic acid | Response | R | -0.12709121 | 0.048469214 | 0.151841734 | -1.180468794 | -1.287177217 | -1.173599673 | -1.213233084 |
| PG(16:0_18:1) | Response | R | 0.257360521 | 0.0489408 | 0.152024767 | -9.567994054 | -9.370571022 | -9.508080077 | -9.410824621 |
| Tryptamine | Response | R | 0.271080906 | 0.049558554 | 0.153339996 | -11.71810647 | -11.45668306 | -11.59414408 | -11.50891244 |
| 3a,7a-Dihydroxycoprostanic acid | Response | R | 0.717424985 | 0.050145888 | 0.153949828 | -11.96403599 | -11.31204646 | -11.81650031 | -11.15837511 |
| 3a,7a-Dihydroxy-5b-cholestane | Response | R | 0.350541518 | 0.052524838 | 0.156977641 | -10.67254424 | -10.43832489 | -10.58875603 | -10.29537873 |
| dIDP | Response | R | 0.664323536 | 0.052183311 | 0.156977641 | -11.45464974 | -10.71180611 | -11.37853089 | -10.46871153 |
| Dihomo-gamma-linolenic acid | Response | R | -0.102671961 | 0.051882448 | 0.156977641 | -5.853995615 | -5.927980701 | -5.794102191 | -5.906924141 |
| LysoPE(22:6) | Response | R | 0.220552344 | 0.052062778 | 0.156977641 | -7.150515523 | -6.943343138 | -7.022616659 | -6.985128572 |
| Indoxyl sulfate | Response | R | 0.420097011 | 0.053611459 | 0.159020456 | -7.256118687 | -6.916777981 | -7.374065547 | -6.879296068 |
| Pivalic acid | Response | R | 0.275544217 | 0.055152092 | 0.162369405 | -9.516146673 | -9.275258906 | -9.601472957 | -9.270892267 |
| LysoPE(16:0) | Response | R | 0.217215315 | 0.056321447 | 0.165195619 | -6.767138809 | -6.567867099 | -6.694265198 | -6.531740613 |
| Phenylacetic acid | Response | R | 0.242102599 | 0.060171061 | 0.174540321 | -9.426892907 | -9.254877688 | -9.452296413 | -9.17398984 |
| Cholic acid | Response | R | 0.605701586 | 0.06249841 | 0.177347139 | -9.4857708 | -9.059748491 | -9.935145166 | -9.48266886 |
| Adrenic acid | Response | R | -0.120339198 | 0.063233431 | 0.177548674 | -6.425868046 | -6.54176209 | -6.416132421 | -6.523669412 |
| 4-Aminopyridine | Response | R | 0.577571682 | 0.063711525 | 0.177626831 | -11.16815534 | -10.56129723 | -10.98899012 | -10.66476574 |
| LysoPE(18:0e) | Response | R | 0.260392694 | 0.065558345 | 0.182132162 | -10.44482468 | -10.20502985 | -10.43518654 | -10.28801895 |
| Acetyl-N-formyl-5-methoxykynurenamine | Response | R | 0.480833109 | 0.066969633 | 0.183820696 | -7.858693772 | -7.489747085 | -8.135833371 | -7.561227163 |
| Lauric acid | Response | R | 0.135353089 | 0.068717507 | 0.18695901 | -6.33523558 | -6.212068832 | -6.344098297 | -6.224375915 |
| 5-Hydroxy-L-tryptophan | Response | R | 0.211969813 | 0.069847778 | 0.188732524 | -12.01507824 | -11.8300633 | -12.04736804 | -11.82599903 |
| 27-Deoxy-5b-cyprinol | Response | R | 0.501184035 | 0.074779146 | 0.195366709 | -9.526117509 | -9.081736824 | -9.395486827 | -8.937187868 |
| FA(21:3) | Response | R | 0.303677991 | 0.074613623 | 0.195366709 | -10.6674094 | -10.50047236 | -10.59508705 | -10.45680297 |
| Pregnanediol | Response | R | 0.303677991 | 0.074613623 | 0.195366709 | -10.6674094 | -10.50047236 | -10.59508705 | -10.45680297 |
| Urocanic acid | Response | R | 0.329334825 | 0.074771716 | 0.195366709 | -10.86097416 | -10.5969491 | -11.026878 | -10.80992606 |
| N-Acetylglutamic acid | Response | R | 0.406529026 | 0.076672071 | 0.199651036 | -10.21962331 | -9.823163059 | -10.13918005 | -9.752941284 |
| Chenodeoxycholic acid glycine conjugate | Response | R | 0.564767388 | 0.082534326 | 0.20798366 | -7.758203784 | -7.320756239 | -7.956642021 | -7.193828588 |
| N-Methyl-L-histidine | Response | R | 0.42028385 | 0.081999203 | 0.20798366 | -10.61960372 | -10.21662217 | -10.6204215 | -10.12847352 |
| Ornithine | Response | R | 0.273712141 | 0.08442291 | 0.209890104 | -8.191773974 | -7.937735609 | -8.199183208 | -7.883380728 |
| D-lactose | Response | R | 0.391936977 | 0.089505444 | 0.217291679 | -10.68579333 | -10.37096049 | -10.65625997 | -10.61975856 |
| LysoPE(20:5) | Response | R | 0.351512837 | 0.097265054 | 0.229080977 | -10.44136473 | -10.09144129 | -10.47103824 | -10.18922563 |
| Salicyluric acid | Response | R | 0.355907566 | 0.101976157 | 0.236644671 | -11.4618486 | -11.1765932 | -11.34662047 | -11.29322941 |
| Ketoleucine | Response | R | 0.206745666 | 0.103957187 | 0.239131839 | -6.640670309 | -6.454906079 | -6.445161457 | -6.518331366 |
| Phenylalanyl-Tyrosine | Response | R | -1.04994811 | 0.001516285 | 0.239573082 | -10.66505541 | -11.71383466 | -10.42779988 | -11.64053548 |
| LysoPI(20:2) | Response | R | 0.238442817 | 0.1051639 | 0.24120441 | -11.87442231 | -11.67818105 | -11.92964782 | -11.73558701 |
| 2-Isopropylmalic acid | Response | R | 0.304858469 | 0.106750205 | 0.242028483 | -11.901329 | -11.60439531 | -11.96814291 | -11.69183975 |
| Oleoyltaurine | Response | R | -0.128167893 | 0.107117143 | 0.242164544 | -10.82370077 | -10.96340313 | -10.76100407 | -10.94976553 |
| L-Lactic acid | Response | R | 0.198692295 | 0.108337823 | 0.24352861 | -4.518563792 | -4.366333484 | -4.585277662 | -4.376337045 |
| PI(18:1_20:4) | Response | R | 0.21399878 | 0.109959262 | 0.245078695 | -9.010432054 | -8.834976088 | -9.004312064 | -8.953715167 |

**Table S4: Relationship between arginine and gut microbes.**

| Microbes | Metabolites | p.value | Cor |
|---|---|---|---|
| s__Bacteroides_cellulosilyticus | L-Arginine | 0.002935154 | 0.442928842 |
| s__Alistipes_onderdonkii | L-Arginine | 0.004709645 | 0.423025478 |
| s__Bilophila_wadsworthia | L-Arginine | 0.007556734 | 0.401872959 |
| s__Bacteroides_nordii | L-Arginine | 0.009222991 | 0.392545195 |
| s__Odoribacter_splanchnicus | L-Arginine | 0.010451208 | 0.386558139 |
| s__Bilophila_unclassified | L-Arginine | 0.010754991 | 0.385170863 |
| s__Paraprevotella_unclassified | L-Arginine | 0.012611231 | 0.377355883 |
| s__Veillonella_atypica | L-Arginine | 0.013193228 | -0.375107981 |
| s__Paraprevotella_clara | L-Arginine | 0.016363949 | 0.36416422 |
| s__Veillonella_unclassified | L-Arginine | 0.020988945 | -0.351053813 |
| s__Eubacterium_ventriosum | L-Arginine | 0.02839577 | 0.334408117 |
| s__Veillonella_parvula | L-Arginine | 0.032646079 | -0.32643481 |
| s__Eubacterium_hallii | L-Arginine | 0.03709575 | 0.318957464 |
| s__Alistipes_senegalensis | L-Arginine | 0.038462499 | 0.316808903 |
| s__Subdoligranulum_unclassified | L-Arginine | 0.043442035 | 0.30947406 |
| s__Alistipes_shahii | L-Arginine | 0.051688134 | 0.298708431 |
| s__Clostridium_bartlettii | L-Arginine | 0.064424019 | 0.284529575 |
| s__Bacteroides_massiliensis | L-Arginine | 0.079660417 | 0.270243295 |
| s__Bacteroides_caccae | L-Arginine | 0.083499661 | 0.266986679 |
| s__Alistipes_putredinis | L-Arginine | 0.088960258 | 0.262550148 |
| s__Coprococcus_comes | L-Arginine | 0.095883286 | 0.257219406 |
| s__Parabacteroides_goldsteinii | L-Arginine | 0.105718555 | 0.250134251 |
| s__Veillonella_dispar | L-Arginine | 0.106854732 | -0.249348736 |
| s__Eggerthella_unclassified | L-Arginine | 0.135345839 | 0.231448681 |
| s__Bacteroides_thetaiotaomicron | L-Arginine | 0.140415786 | 0.228567129 |
| s__Escherichia_unclassified | L-Arginine | 0.141870219 | -0.227754748 |
| s__Lachnospiraceae_bacterium_7_1_58FAA | L-Arginine | 0.144118211 | 0.226511152 |
| s__Bacteroides_uniformis | L-Arginine | 0.152215755 | 0.222146704 |
| s__Alistipes_indistinctus | L-Arginine | 0.15814858 | 0.219055926 |
| s__Parabacteroides_merdae | L-Arginine | 0.165920835 | 0.215132226 |
| s__Anaerotruncus_colihominis | L-Arginine | 0.167671511 | 0.214266901 |
| s__Lachnospiraceae_bacterium_5_1_63FAA | L-Arginine | 0.171008597 | 0.212635368 |
| s__Bacteroides_dorei | L-Arginine | 0.179955995 | 0.208371192 |
| s__Bacteroides_intestinalis | L-Arginine | 0.192771673 | 0.202519898 |
| s__Streptococcus_parasanguinis | L-Arginine | 0.210299987 | -0.194946509 |
| s__Clostridium_leptum | L-Arginine | 0.215427368 | 0.192815009 |
| s__Ruminococcus_lactaris | L-Arginine | 0.218311549 | 0.191631467 |
| s__Barnesiella_intestinihominis | L-Arginine | 0.253472172 | 0.177996736 |
| s__Oscillibacter_unclassified | L-Arginine | 0.25593321 | 0.177091664 |
| s__Ruminococcus_obeum | L-Arginine | 0.28691281 | 0.166162601 |
| s__Escherichia_coli | L-Arginine | 0.302697155 | -0.160889265 |
| s__Lachnospiraceae_bacterium_1_1_57FAA | L-Arginine | 0.308549694 | 0.158978971 |
| s__Bacteroides_ovatus | L-Arginine | 0.315519326 | 0.156733903 |
| s__Clostridium_asparagiforme | L-Arginine | 0.316813452 | 0.156320506 |
| s__Klebsiella_pneumoniae | L-Arginine | 0.354285132 | -0.144780929 |
| s__Clostridium_hathewayi | L-Arginine | 0.360948167 | 0.142808815 |
| s__Lachnospiraceae_bacterium_3_1_46FAA | L-Arginine | 0.390270478 | -0.134379709 |
| s__Dialister_invisus | L-Arginine | 0.392695022 | -0.133699747 |
| s__Alistipes_finegoldii | L-Arginine | 0.403275013 | 0.130760827 |
| s__Lachnospiraceae_bacterium_2_1_58FAA | L-Arginine | 0.471746078 | 0.112715131 |
| s__Dorea_longicatena | L-Arginine | 0.48019662 | 0.110589737 |
| s__Clostridium_clostridioforme | L-Arginine | 0.495321934 | 0.106833891 |
| s__Streptococcus_salivarius | L-Arginine | 0.500028154 | -0.105677388 |
| s__Clostridium_bolteae | L-Arginine | 0.524740773 | -0.099692986 |
| s__Bacteroides_xylanisolvens | L-Arginine | 0.52571046 | 0.099461065 |
| s__Flavonifractor_plautii | L-Arginine | 0.556036221 | 0.092309954 |
| s__Bacteroides_salyersiae | L-Arginine | 0.568742392 | 0.089368872 |
| s__Parasutterella_excrementihominis | L-Arginine | 0.572658652 | 0.08846852 |
| s__Ruminococcus_gnavus | L-Arginine | 0.586991642 | -0.085196969 |
| s__Faecalibacterium_prausnitzii | L-Arginine | 0.620312002 | 0.077724909 |
| s__Parabacteroides_unclassified | L-Arginine | 0.620712646 | 0.077636126 |
| s__Coprococcus_catus | L-Arginine | 0.626582284 | 0.076338134 |
| less_0.01p_total | L-Arginine | 0.658459056 | 0.069163395 |
| s__Bacteroides_vulgatus | L-Arginine | 0.66007691 | 0.069023773 |
| s__Clostridium_symbiosum | L-Arginine | 0.67588034 | -0.065623252 |
| s__Bacteroidales_bacterium_ph8 | L-Arginine | 0.688234548 | 0.062985546 |
| s__Eubacterium_rectale | L-Arginine | 0.693197068 | 0.061930867 |
| s__Ruminococcus_callidus | L-Arginine | 0.698037106 | 0.060904811 |
| s__Bacteroides_finegoldii | L-Arginine | 0.703235926 | -0.059805484 |
| s__Ruminococcus_torques | L-Arginine | 0.706468576 | 0.059123344 |

| | | | |
|---|---|---|---|
| s__Collinsella_aerofaciens | L-Arginine | 0.706576977 | -0.059100488 |
| s__Ruminococcus_bromii | L-Arginine | 0.710880428 | 0.058194105 |
| s__Eubacterium_eligens | L-Arginine | 0.718731938 | -0.056545222 |
| s__Dorea_formicigenerans | L-Arginine | 0.736333575 | 0.052870184 |
| s__Eubacterium_ramulus | L-Arginine | 0.757124973 | -0.048564687 |
| s__Clostridium_citroniae | L-Arginine | 0.780866264 | 0.043690812 |
| s__Akkermansia_muciniphila | L-Arginine | 0.781277179 | 0.043606824 |
| s__Bifidobacterium_pseudocatenulatum | L-Arginine | 0.79075315 | -0.041673281 |
| s__Roseburia_hominis | L-Arginine | 0.792161875 | -0.041386361 |
| s__Burkholderiales_bacterium_1_1_47 | L-Arginine | 0.79604309 | 0.040596545 |
| s__Enterobacter_cloacae | L-Arginine | 0.799697994 | 0.039853686 |
| s__Bifidobacterium_longum | L-Arginine | 0.802480474 | 0.039288724 |
| s__Parabacteroides_distasonis | L-Arginine | 0.814139826 | 0.036926608 |
| s__Bacteroides_fragilis | L-Arginine | 0.81501621 | 0.036749389 |
| s__Haemophilus_parainfluenzae | L-Arginine | 0.830698652 | -0.033585534 |
| s__Bacteroides_stercoris | L-Arginine | 0.860434042 | 0.027621377 |
| s__Roseburia_inulinivorans | L-Arginine | 0.873760015 | 0.024961414 |
| s__Sutterella_wadsworthensis | L-Arginine | 0.87548183 | -0.024618244 |
| s__Holdemania_filiformis | L-Arginine | 0.885277188 | 0.022668067 |
| s__Fusobacterium_nucleatum | L-Arginine | 0.890887772 | -0.021552576 |
| s__Roseburia_intestinalis | L-Arginine | 0.960860904 | 0.007710712 |
| s__Bacteroides_faecis | L-Arginine | 0.993554122 | 0.001269412 |

**Table S5. Differential metabolites between mice transplanted with R or NR patients' stool samples.**

| feature | metadata | value | coef | pval | qval | mean_NR | mean_R | median_NR | median_R |
|---|---|---|---|---|---|---|---|---|---|
| Acylcarnitine(22:4) | Response | R | -3.36E-06 | 1.63E-06 | 0.000599771 | 7.31E-06 | 3.95E-06 | 7.48E-06 | 3.76E-06 |
| Linoleic Acid | Response | R | 5.06E-06 | 0.000570232 | 0.041969093 | 1.10E-05 | 1.60E-05 | 1.05E-05 | 1.60E-05 |
| LysoPC(18:0) | Response | R | 0.010780663 | 0.000523106 | 0.041969093 | 0.040774072 | 0.051554735 | 0.042537559 | 0.050267898 |
| LysoPC(19:0) | Response | R | 0.000287757 | 0.000436581 | 0.041969093 | 0.000619665 | 0.000907422 | 0.000625699 | 0.000878651 |
| Gamma Glutamylglutamic Acid | Response | R | -5.35E-06 | 0.000253417 | 0.041969093 | 1.52E-05 | 9.83E-06 | 1.57E-05 | 9.24E-06 |
| D-Glutamylglycine | Response | R | -5.42E-05 | 0.000884263 | 0.04648697 | 0.000162132 | 0.000107949 | 0.000178204 | 0.000104818 |
| LysoPC(20:0) | Response | R | 0.000137822 | 0.000805064 | 0.04648697 | 0.000283982 | 0.000421804 | 0.000272887 | 0.000443555 |
| PC(40:6) | Response | R | 0.002195642 | 0.001877226 | 0.086352399 | 0.006158541 | 0.008354182 | 0.006204875 | 0.008238601 |
| 3-Methylhistamine | Response | R | 0.001019374 | 0.002956183 | 0.11096723 | 0.002086298 | 0.003105672 | 0.00211754 | 0.002852767 |
| Adenosine Monophosphate | Response | R | 0.000155069 | 0.003893124 | 0.11096723 | 0.000246813 | 0.000401882 | 0.000253408 | 0.000354286 |
| Acylcarnitine(20:5) | Response | R | 8.97E-05 | 0.003030515 | 0.11096723 | 0.000224763 | 0.000314478 | 0.000230244 | 0.000298209 |
| Cortisol | Response | R | 4.74E-06 | 0.003920038 | 0.11096723 | 1.25E-05 | 1.73E-05 | 1.23E-05 | 1.76E-05 |
| Uracil | Response | R | -2.43E-05 | 0.003330787 | 0.11096723 | 6.19E-05 | 3.76E-05 | 5.96E-05 | 3.60E-05 |
| 1H-Indole-4-Carboxaldehyde | Response | R | 0.000161136 | 0.005212117 | 0.111666126 | 0.000640686 | 0.000801822 | 0.000634234 | 0.000819109 |
| Nicotinamide Ribose | Response | R | -1.24E-05 | 0.004786751 | 0.111666126 | 3.75E-05 | 2.50E-05 | 3.36E-05 | 2.40E-05 |
| Alpha-Linolenic Acid | Response | R | 2.43E-05 | 0.005565329 | 0.111666126 | 0.000107933 | 0.000132273 | 0.000112872 | 0.000124969 |
| Tyr-Glu | Response | R | -7.53E-06 | 0.006372252 | 0.111666126 | 2.57E-05 | 1.82E-05 | 2.56E-05 | 1.52E-05 |
| Gamma-Glutamyltyrosine | Response | R | -7.53E-06 | 0.006372252 | 0.111666126 | 2.57E-05 | 1.82E-05 | 2.56E-05 | 1.52E-05 |
| N-Palmitoylsphingosine | Response | R | -2.03E-05 | 0.006047386 | 0.111666126 | 4.19E-05 | 2.16E-05 | 4.29E-05 | 2.34E-05 |
| PC(42:9) | Response | R | 1.93E-05 | 0.00494532 | 0.111666126 | 0.000183301 | 0.000202558 | 0.000179548 | 0.000203918 |
| Ala-Arg | Response | R | 8.48E-06 | 0.006329248 | 0.111666126 | 1.61E-06 | 1.01E-05 | 1.02E-06 | 7.36E-06 |
| PC(35:5) | Response | R | -0.00014263 | 0.006860615 | 0.114759371 | 0.000292577 | 0.000149947 | 0.000277104 | 0.000101345 |
| Indoleacrylic Acid | Response | R | 0.001524749 | 0.007581585 | 0.114788737 | 0.006054424 | 0.007579173 | 0.005973173 | 0.007684586 |
| L-Tryptophan | Response | R | 0.001494609 | 0.007798148 | 0.114788737 | 0.005601801 | 0.00709641 | 0.005735311 | 0.007145565 |
| 13Z-Docosenamide | Response | R | -0.005367568 | 0.007539399 | 0.114788737 | 0.009452593 | 0.004085025 | 0.007787383 | 0.00379379 |
| PC(35:6) | Response | R | -0.0002458 | 0.008908416 | 0.126088347 | 0.000533524 | 0.000287724 | 0.000506425 | 0.00018972 |
| Thr-Glu | Response | R | -1.65E-05 | 0.011218287 | 0.132544859 | 5.84E-05 | 4.19E-05 | 5.51E-05 | 3.92E-05 |
| Lys-Glu | Response | R | -3.10E-05 | 0.011601794 | 0.132544859 | 0.000163851 | 0.000132862 | 0.000160185 | 0.000126338 |
| Sphinganine 1-Phosphate | Response | R | -6.01E-05 | 0.009835244 | 0.132544859 | 0.000192109 | 0.000131966 | 0.000202106 | 0.000128164 |
| LysoPE(18:1) | Response | R | -0.00013983 | 0.010300477 | 0.132544859 | 0.00074608 | 0.000606251 | 0.000772736 | 0.000594769 |
| 1-Methyladenine | Response | R | -4.18E-05 | 0.012245992 | 0.132544859 | 0.000153556 | 0.000111718 | 0.000148104 | 0.000103924 |
| L-Cystathionine | Response | R | -5.29E-06 | 0.012070093 | 0.132544859 | 1.77E-05 | 1.24E-05 | 1.74E-05 | 1.12E-05 |
| Tauro Cholic Acid | Response | R | -4.64E-05 | 0.011334833 | 0.132544859 | 6.09E-05 | 1.45E-05 | 3.12E-05 | 8.64E-06 |
| Tauro Cholic Acid+NH4 | Response | R | -0.000189731 | 0.010779234 | 0.132544859 | 0.000251331 | 6.16E-05 | 0.00013213 | 4.43E-05 |
| L-Arginine | Response | R | 0.002279402 | 0.015728693 | 0.150974202 | 0.000409914 | 0.002689316 | 0.000196532 | 0.001768548 |
| Deoxyuridine | Response | R | 2.10E-06 | 0.015903005 | 0.150974202 | 4.65E-06 | 6.75E-06 | 4.61E-06 | 6.75E-06 |
| Glu-His | Response | R | -5.50E-06 | 0.014593299 | 0.150974202 | 1.89E-05 | 1.34E-05 | 1.73E-05 | 1.24E-05 |
| Acylcarnitine(22:6) | Response | R | 1.45E-05 | 0.015999983 | 0.150974202 | 3.35E-05 | 4.80E-05 | 3.43E-05 | 5.07E-05 |
| Glu-Met | Response | R | 6.69E-06 | 0.015292004 | 0.150974202 | 4.94E-06 | 1.16E-05 | 5.15E-06 | 1.11E-05 |
| Sphingosine-1-Phosphate | Response | R | -6.57E-05 | 0.017111542 | 0.157426186 | 0.000276464 | 0.000210759 | 0.000264735 | 0.000201147 |
| Glycerophosphocholine | Response | R | 0.008902735 | 0.019512702 | 0.163555105 | 0.031805348 | 0.040708083 | 0.030946239 | 0.039680402 |
| 1-Methyladenosine | Response | R | -0.000139468 | 0.018418547 | 0.163555105 | 0.000543208 | 0.00040374 | 0.000525352 | 0.000366338 |
| Guanosine 5'-Monophosphate | Response | R | 7.56E-06 | 0.018947462 | 0.163555105 | 1.67E-05 | 2.42E-05 | 1.71E-05 | 2.19E-05 |
| Hydrocinnamic Acid | Response | R | 1.08E-05 | 0.019555502 | 0.163555105 | 1.74E-05 | 2.83E-05 | 1.66E-05 | 3.03E-05 |
| LysoPC(22:4) | Response | R | -5.49E-05 | 0.021098055 | 0.172535205 | 0.000209826 | 0.000154894 | 0.000188799 | 0.000158496 |
| N-(3-Acetamidopropyl)Pyrrolidin-2-One | Response | R | -0.000234011 | 0.02318859 | 0.18272811 | 0.000428174 | 0.000194163 | 0.000385204 | 0.000169116 |
| PC(40:7) | Response | R | -0.001268225 | 0.023337557 | 0.18272811 | 0.006718076 | 0.005449851 | 0.006677605 | 0.005645344 |
| Nicotine | Response | R | -9.73E-06 | 0.025543544 | 0.195833839 | 4.54E-05 | 3.57E-05 | 4.45E-05 | 3.43E-05 |
| N-Acetylserotonin | Response | R | 3.25E-06 | 0.027955638 | 0.201719117 | 4.69E-06 | 7.95E-06 | 4.85E-06 | 8.25E-06 |
| S-Glutathionyl-L-Cysteine | Response | R | -6.14E-06 | 0.027916805 | 0.201719117 | 2.42E-05 | 1.80E-05 | 2.28E-05 | 1.73E-05 |
| PE(p38:6) | Response | R | 4.93E-05 | 0.02749477 | 0.201719117 | 0.000182 | 0.000231322 | 0.000171816 | 0.000213577 |
| Tauro-b-Muricholic Acid+NH4 | Response | R | -0.000552601 | 0.033024226 | 0.23370991 | 0.000787828 | 0.000235227 | 0.000342873 | 0.000169243 |
| 1-(Beta-D-Ribofuranosyl)-1,4-Dihydronicotinamide | Response | R | -2.07E-05 | 0.037379599 | 0.241327937 | 7.81E-05 | 5.73E-05 | 7.36E-05 | 5.39E-05 |
| Uridine 5'-Monophosphate | Response | R | 1.19E-06 | 0.037117456 | 0.241327937 | 2.35E-06 | 3.55E-06 | 2.51E-06 | 3.08E-06 |
| C18:3-OH Carnitine | Response | R | -5.32E-06 | 0.035784454 | 0.241327937 | 2.14E-05 | 1.61E-05 | 2.43E-05 | 1.67E-05 |
| Acylcarnitine(20:2) | Response | R | -1.43E-05 | 0.036394504 | 0.241327937 | 6.78E-05 | 5.35E-05 | 7.22E-05 | 5.44E-05 |
| Tauro-b-Muricholic Acid | Response | R | -0.000119307 | 0.03494191 | 0.241327937 | 0.000167622 | 4.83E-05 | 7.25E-05 | 3.32E-05 |
| Dimethylglycine | Response | R | -0.000498267 | 0.039556055 | 0.250469241 | 0.001783931 | 0.001285664 | 0.001592809 | 0.001207945 |
| Benzylamine | Response | R | -5.98E-05 | 0.040156753 | 0.250469241 | 0.00065874 | 0.000598894 | 0.000650723 | 0.000594261 |
| 3-Hydroxyanthranilic Acid | Response | R | -0.000195773 | 0.046789708 | 0.286976875 | 0.000758296 | 0.000562523 | 0.000676938 | 0.000558176 |
| Guanine | Response | R | -1.63E-05 | 0.049939936 | 0.301276992 | 4.24E-05 | 2.61E-05 | 3.40E-05 | 2.23E-05 |
| Docosahexaenoic Acid | Response | R | 0.006646195 | 0.000358507 | 0.09392878 | 0.023227984 | 0.029874179 | 0.02401482 | 0.028660284 |
| Asp-His | Response | R | 0.00010052 | 0.001251709 | 0.163973868 | 0.000129049 | 0.00022957 | 0.000138832 | 0.000230587 |
| 16-Hydroxypalmitic Acid | Response | R | 0.000112352 | 0.005638886 | 0.197393873 | 0.000456026 | 0.000568378 | 0.000454183 | 0.000589036 |
| Caffeate | Response | R | -3.55E-05 | 0.007534117 | 0.197393873 | 8.32E-05 | 4.77E-05 | 7.83E-05 | 5.20E-05 |
| Corticosterone | Response | R | 4.52E-05 | 0.006701029 | 0.197393873 | 0.000109843 | 0.00015504 | 0.000112466 | 0.000142042 |
| FA(22:4) | Response | R | -0.000482039 | 0.003342678 | 0.197393873 | 0.002037316 | 0.001555277 | 0.001891996 | 0.001569137 |
| Glu-Ser | Response | R | -1.62E-05 | 0.006915235 | 0.197393873 | 3.09E-05 | 1.47E-05 | 2.67E-05 | 1.28E-05 |
| LysoPE(22:4) | Response | R | -2.47E-05 | 0.002882273 | 0.197393873 | 6.44E-05 | 3.98E-05 | 5.66E-05 | 3.93E-05 |
| LysoPS(22:6) | Response | R | -1.81E-05 | 0.00534502 | 0.197393873 | 5.59E-05 | 3.78E-05 | 4.96E-05 | 3.71E-05 |
| N-Acetyl-Dl-Methionine | Response | R | 5.98E-06 | 0.005344462 | 0.197393873 | 3.55E-06 | 9.53E-06 | 3.76E-06 | 9.79E-06 |
| 2-Ketobutyric Acid | Response | R | 0.000171135 | 0.012329587 | 0.232440318 | 0.000405435 | 0.00057657 | 0.000431762 | 0.000554611 |
| FA(24:5) | Response | R | 0.000102757 | 0.013307652 | 0.232440318 | 0.000481727 | 0.000584484 | 0.000476347 | 0.000603036 |
| LysoPS(18:1) | Response | R | -1.52E-05 | 0.009765338 | 0.232440318 | 2.54E-05 | 1.02E-05 | 2.62E-05 | 1.05E-05 |
| PI(16:0_20:4) | Response | R | 0.000358217 | 0.012044202 | 0.232440318 | 0.001003175 | 0.001361392 | 0.000966729 | 0.001332669 |
| Thymidine | Response | R | 6.30E-05 | 0.012728514 | 0.232440318 | 0.000119222 | 0.000182222 | 0.000108731 | 0.000163423 |
| Alpha-Hydroxyisobutyric Acid | Response | R | -0.000538237 | 0.016830032 | 0.244970471 | 0.001533257 | 0.00099502 | 0.001570844 | 0.000984631 |
| LysoPG(18:1) | Response | R | -3.40E-05 | 0.015239217 | 0.244970471 | 0.000140785 | 0.000106791 | 0.000130376 | 0.000106352 |
| Malic Acid | Response | R | 0.000260325 | 0.016175654 | 0.244970471 | 0.000625084 | 0.000885409 | 0.000701165 | 0.000885192 |
| 3-Hydroxy-3-Methylglutarate | Response | R | 0.000255989 | 0.018675835 | 0.257529929 | 0.000576893 | 0.000832882 | 0.000570785 | 0.000788102 |
| FA(24:6) | Response | R | 0.000249013 | 0.020632934 | 0.258372519 | 0.000659962 | 0.000908975 | 0.000655561 | 0.000907756 |
| Glutaric Acid | Response | R | 4.70E-05 | 0.020709248 | 0.258372519 | 0.000141773 | 0.000188809 | 0.000144169 | 0.000185667 |
| Eicosapentaenoic Acid | Response | R | 0.001007868 | 0.023265483 | 0.277070748 | 0.002769465 | 0.003777333 | 0.00271157 | 0.003499488 |
| Valproic Acid | Response | R | -8.40E-06 | 0.024521242 | 0.27932893 | 4.66E-05 | 3.82E-05 | 4.59E-05 | 3.63E-05 |
| LysoPE(20:5) | Response | R | 9.24E-05 | 0.028236446 | 0.308247872 | 0.000380737 | 0.000473107 | 0.00037565 | 0.000474461 |
| LysoPG(20:4) | Response | R | -1.25E-05 | 0.030141511 | 0.315883039 | 3.86E-05 | 2.61E-05 | 3.49E-05 | 2.57E-05 |
| Taurocholic Acid | Response | R | -0.002276359 | 0.032358747 | 0.3260766 | 0.00299632 | 0.00071996 | 0.00119515 | 0.000488757 |
| LysoPC(20:5) | Response | R | 0.000277813 | 0.034978744 | 0.327301106 | 0.000786038 | 0.001063851 | 0.000822435 | 0.001021145 |
| Uridine | Response | R | -0.000143769 | 0.034427765 | 0.327301106 | 0.000378913 | 0.000235144 | 0.000344446 | 0.000207449 |
| 13(S)-Hpot | Response | R | -2.15E-05 | 0.037822296 | 0.336447573 | 6.94E-05 | 4.79E-05 | 7.25E-05 | 4.52E-05 |
| PG(18:1_18:2) | Response | R | -2.25E-05 | 0.038524531 | 0.336447573 | 5.57E-05 | 3.32E-05 | 4.85E-05 | 2.43E-05 |
| Stearic Acid | Response | R | 0.008467287 | 0.042056788 | 0.34433995 | 0.05856762 | 0.067034907 | 0.059248405 | 0.064362989 |
| Urocanic Acid | Response | R | 1.52E-05 | 0.041008763 | 0.34433995 | 1.94E-05 | 3.46E-05 | 2.18E-05 | 2.76E-05 |
| Fumarate | Response | R | 2.68E-05 | 0.04902042 | 0.346239381 | 7.38E-05 | 0.000100535 | 8.29E-05 | 9.85E-05 |
| Lactate | Response | R | 0.004570946 | 0.045615262 | 0.346239381 | 0.016702694 | 0.021273639 | 0.01496542 | 0.019386525 |
| Pregnenolone | Response | R | 5.14E-05 | 0.044776246 | 0.346239381 | 0.000187952 | 0.000239379 | 0.000190774 | 0.000253486 |
| Tryptamine | Response | R | 4.40E-06 | 0.049746452 | 0.346239381 | 1.57E-05 | 2.01E-05 | 1.55E-05 | 1.99E-05 |

**Table S6. Differentially expressed host genes based on RNA-seq analysis.**

| gene_id | mean_R | mean_NR | median_R | median_NR | log2FoldChange | pvalue | padj | gene_name | gene_chr | gene_start | gene_end | gene_strand | gene_length | gene_biotype | gene_descriptio | tf_family |
|---|---|---|---|---|---|---|---|---|---|---|---|---|---|---|---|---|
| 170930 | 1566.197739 | 555.4413452 | 1581.716187 | 589.1634171 | 1.496240136 | 2.26E-16 | 5.88E-12 | Sumo2 | NC_000077.7 | 115413935 | 115427056 | - | 998 | protein_coding | small ubiquitin-li | - |
| 78294 | 9180.475121 | 4087.816039 | 9448.19745 | 4026.599725 | 1.167276132 | 1.15E-13 | 1.50E-09 | Rps27a | NC_000077.7 | 29495842 | 29498040 | - | 821 | protein_coding | ribosomal protein | ubiquitin |
| 16016 | 68.6163979 | 4.51266894 | 57.24506591 | 2.309413328 | 3.871746248 | 1.91E-09 | 1.66E-05 | Ighg2b | NC_000078.7 | 113267934 | 113271553 | - | 1222 | C_region | - && P01867.3 F | C1-set |
| 50908 | 1628.659136 | 612.4153716 | 1914.052023 | 555.4345962 | 1.411027331 | 6.16E-09 | 4.01E-05 | C1s1 | NC_000072.7 | 124507303 | 124519340 | - | 3103 | protein_coding | complement com | Trypsin |
| 22287 | 26.71550617 | 1.9583745 | 28.06321581 | 2.129811955 | 3.65170004 | 3.91E-08 | 0.000169915 | Scgb1a1 | NC_000085.7 | 9061006 | 9065320 | - | 468 | protein_coding | secretoglobin%2 | - |
| 57277 | 1282.836374 | 231.3753675 | 1407.61053 | 224.984718 | 2.470464027 | 1.13E-07 | 0.000419538 | Slurp1 | NC_000081.7 | 74598493 | 74599872 | - | 522 | protein_coding | secreted Ly6/Pla | - |
| 170835 | 26.08244045 | 1.9583745 | 23.93695417 | 2.129811955 | 3.62054119 | 4.34E-07 | 0.001258148 | Inpp5j | NC_000077.7 | 3444272 | 3455521 | - | 3968 | protein_coding | inositol polyphos | Exo_endo_phos |
| 11898 | 643.7130425 | 199.7781883 | 642.5542381 | 216.7253435 | 1.687543698 | 4.35E-07 | 0.001258148 | Ass1 | NC_000068.8 | 31360282 | 31410682 | + | 1631 | protein_coding | argininosuccinate | - |
| 14998 | 1107.530798 | 437.9490491 | 1244.494976 | 411.8619686 | 1.338166172 | 5.89E-07 | 0.001496557 | H2-DMa | NC_000083.7 | 34338667 | 34358075 | + | 8325 | protein_coding | histocompatibility | C1-set |
| 72361 | 133.6572324 | 23.4611248 | 82.13170367 | 23.40400556 | 2.504954421 | 6.32E-07 | 0.001496557 | Ces2g | NC_000074.7 | 105688350 | 105696169 | + | 2859 | protein_coding | carboxylesterase | COesterase |
| 22271 | 1625.05884 | 783.3973071 | 1738.390305 | 772.1549079 | 1.052734104 | 7.36E-07 | 0.001597688 | Upp1 | NC_000077.7 | 9068008 | 9086170 | + | 1739 | protein_coding | uridine phosphory | - |
| 71939 | 941.2783061 | 154.5751276 | 926.4057281 | 122.2884173 | 2.604857772 | 9.70E-07 | 0.001944028 | Apol6 | NC_000081.7 | 76929195 | 76941308 | + | 3529 | protein_coding | apolipoprotein L | - |
| 56620 | 763.9060175 | 362.4049725 | 857.718221 | 386.5898353 | 1.074943561 | 1.16E-06 | 0.002156188 | Clec4n | NC_000072.7 | 123206802 | 123223983 | + | 1245 | protein_coding | C-type lectin dom | Lectin_C |
| 238393 | 757.3340984 | 120.0605157 | 857.7134703 | 100.5595174 | 2.655458759 | 2.04E-06 | 0.00323546 | Serpina3f | NC_000078.7 | 104180803 | 104187388 | + | 2290 | protein_coding | serine (or cystein | Serpin |
| 24108 | 800.2182529 | 153.4379225 | 716.4217989 | 114.9932608 | 2.381185751 | 2.11E-06 | 0.00323546 | Ubd | NC_000083.7 | 37504783 | 37506992 | + | 937 | protein_coding | ubiquitin D && I | ubiquitin |
| 328561 | 383.0272044 | 108.4479005 | 358.7367635 | 100.6418025 | 1.819156735 | 2.23E-06 | 0.00323546 | Apol10b | NC_000081.7 | 77468354 | 77482796 | - | 2744 | protein_coding | apolipoprotein L | - |
| 20389 | 13.45892852 | 0.630532202 | 12.21375044 | 0.016232487 | 4.094277229 | 2.71E-06 | 0.003712656 | Sftpc | NC_000080.7 | 70758381 | 70761521 | - | 804 | protein_coding | surfactant associa | - |
| 19171 | 2458.123199 | 1125.918572 | 2316.537979 | 1050.619509 | 1.126365376 | 3.35E-06 | 0.004361683 | Psmb10 | NC_000074.7 | 106662360 | 106665024 | - | 1270 | protein_coding | proteasome (pros | Proteasome |
| 16161 | 584.6399741 | 231.128152 | 499.1370127 | 232.2761836 | 1.338253411 | 3.96E-06 | 0.004689545 | Il12rb1 | NC_000074.7 | 71261005 | 71276186 | + | 5590 | protein_coding | interleukin 12 rec | - |
| 21355 | 2414.922518 | 1115.623801 | 2196.984705 | 1000.677395 | 1.114089601 | 3.96E-06 | 0.004689545 | Tap2 | NC_000083.7 | 34423453 | 34435295 | + | 2490 | protein_coding | transporter 2%2C | ABC_membrane |
| 225594 | 599.1593247 | 133.9854267 | 797.2474023 | 118.7478115 | 2.159739144 | 5.01E-06 | 0.005411496 | Gm4841 | NC_000084.7 | 60401373 | 60406339 | - | 2838 | protein_coding | predicted gene 48 | - |
| 20715 | 1411.751061 | 552.6356257 | 1562.658794 | 532.5619749 | 1.352763829 | 5.24E-06 | 0.005411496 | Serpina3g | NC_000078.7 | 104202406 | 104208198 | + | 1908 | protein_coding | serine (or cystein | Serpin |
| 16912 | 2884.140337 | 967.6450308 | 2831.183497 | 762.8430777 | 1.575444304 | 5.40E-06 | 0.005411496 | Psmb9 | NC_000083.7 | 34400961 | 34406304 | - | 1110 | protein_coding | proteasome (pros | Proteasome |
| 16142 | 21.49676528 | 0.661271116 | 15.65651382 | 0.909372407 | 4.755685905 | 1.03E-05 | 0.008949417 | Iglv1 | NC_000082.7 | 18903767 | 18904210 | - | 351 | V_segment | - && P01727.1 F | V-set |
| 223626 | 719.2366313 | 349.138674 | 719.492914 | 352.9590397 | 1.042335109 | 1.47E-05 | 0.011970226 | Them6 | NC_000081.7 | 74593083 | 74596222 | + | 1488 | protein_coding | thioesterase supe | - |
| 76074 | 616.8233185 | 188.9159609 | 582.2749815 | 175.9485982 | 1.706315633 | 1.66E-05 | 0.012757116 | Gbp8 | NC_000071.7 | 105160379 | 105201475 | - | 4396 | protein_coding | guanylate-binding | - |
| 18400 | 129.0266331 | 44.03353454 | 153.181433 | 43.53602924 | 1.547847416 | 2.49E-05 | 0.016608505 | Slc22a18 | NC_000073.7 | 143027463 | 143053071 | + | 1791 | protein_coding | solute carrier fam | MFS_1 |
| 53897 | 7.518221091 | 1.39E-17 | 8.309028025 | 1.39E-17 | 5.937389735 | 2.49E-05 | 0.016608505 | Gal3st1 | NC_000077.7 | 3933636 | 3949328 | + | 2902 | protein_coding | galactose-3-O-su | - |
| 21354 | 5441.907392 | 2222.469708 | 5669.604167 | 1854.417345 | 1.291897428 | 2.60E-05 | 0.016787509 | Tap1 | NC_000083.7 | 34406530 | 34416199 | + | 2950 | protein_coding | transporter 1%2C | ABC_membrane |
| 14969 | 5090.036548 | 2002.283744 | 5026.040477 | 1818.201555 | 1.34594984 | 2.64E-05 | 0.016787509 | H2-Eb1 | NC_000083.7 | 34524841 | 34535648 | + | 1669 | protein_coding | histocompatibility | - |
| 16913 | 4500.659125 | 1979.333613 | 3984.28362 | 1667.513904 | 1.185082895 | 2.85E-05 | 0.016899129 | Psmb8 | NC_000083.7 | 34417169 | 34420428 | + | 1223 | protein_coding | proteasome (pros | Proteasome |
| 16149 | 11198.29938 | 4359.714117 | 11300.77109 | 3974.952226 | 1.360937493 | 2.93E-05 | 0.016980743 | Cd74 | NC_000084.7 | 60936921 | 60945724 | + | 1415 | protein_coding | CD74 antigen (in | - |
| 14469 | 11708.67356 | 2944.241557 | 12257.29994 | 2334.356971 | 1.991537093 | 3.03E-05 | 0.016980743 | Gbp2 | NC_000069.7 | 142326424 | 142343769 | + | 2455 | protein_coding | guanylate binding | - |
| 14999 | 1482.955528 | 681.6650305 | 1450.430165 | 605.9775918 | 1.121196786 | 3.06E-05 | 0.016980743 | H2-DMb1 | NC_000083.7 | 34372165 | 34379203 | + | 1284 | protein_coding | histocompatibility | C1-set |
| 244238 | 189.9063829 | 79.82533887 | 216.3161807 | 77.65949707 | 1.249333826 | 4.04E-05 | 0.021100182 | Mrgpre | NC_000073.7 | 143332100 | 143338237 | - | 3779 | protein_coding | MAS-related GPI | - |
| 213233 | 1054.15583 | 503.9187204 | 1060.52008 | 501.7466043 | 1.064494126 | 4.12E-05 | 0.021100182 | Tapbpl | NC_000072.7 | 125200890 | 125208823 | - | 2492 | protein_coding | TAP binding prot | V-set |
| 15930 | 141.0031999 | 32.46421332 | 165.616507 | 34.01691758 | 2.114623029 | 4.37E-05 | 0.021885111 | Ido1 | NC_000074.7 | 25074149 | 25086968 | - | 1771 | protein_coding | indoleamine 2%2 | - |
| 17472 | 4112.778936 | 1014.051805 | 3440.529403 | 942.6423645 | 2.019815293 | 5.04E-05 | 0.023378136 | Gbp4 | NC_000071.7 | 105263633 | 105287452 | - | 5236 | protein_coding | guanylate binding | - |
| 11988 | 4953.000488 | 2338.044252 | 4865.697727 | 2163.916757 | 1.082924304 | 5.10E-05 | 0.023378136 | Slc7a2 | NC_000074.7 | 41315404 | 41375107 | + | 9067 | protein_coding | solute carrier fam | - |
| 381058 | 159.9505261 | 33.21647319 | 160.1203663 | 29.86936083 | 2.262504668 | 5.93E-05 | 0.024695672 | Unc93a | NC_000083.7 | 13327504 | 13362553 | - | 3412 | protein_coding | unc-93 homolog | - |
| 14612 | 144.3176412 | 37.53781427 | 136.2299736 | 40.38274838 | 1.938275338 | 5.96E-05 | 0.024695672 | Gja4 | NC_000070.7 | 127205213 | 127207832 | - | 1686 | protein_coding | gap junction prot | Connexin |
| 14621 | 417.5875432 | 158.9488929 | 453.2840633 | 168.8181951 | 1.392654794 | 5.97E-05 | 0.024695672 | Gjb4 | NC_000070.7 | 127234902 | 127247929 | - | 2772 | protein_coding | gap junction prot | Connexin |
| 55932 | 2753.049633 | 866.2225458 | 2736.562637 | 769.2970012 | 1.668030694 | 6.40E-05 | 0.025658528 | Gbp3 | NC_000069.7 | 142265781 | 142278974 | + | 3175 | protein_coding | guanylate binding | - |
| 433470 | 1028.544696 | 436.3005628 | 1167.549803 | 445.5186042 | 1.236853039 | 6.93E-05 | 0.026393494 | AA467197 | NC_000068.8 | 122479807 | 122482996 | + | 640 | protein_coding | expressed sequen | - |
| 14961 | 4259.687157 | 1817.176525 | 3335.540826 | 1703.000778 | 1.228970243 | 6.97E-05 | 0.026393494 | H2-Ab1 | NC_000083.7 | 34482201 | 34488392 | + | 1193 | protein_coding | histocompatibility | C1-set |
| novel.8 | 358.1848109 | 95.43278768 | 395.0516762 | 93.42911412 | 1.906771745 | 7.07E-05 | 0.026393494 | - | NC_000067.7 | 90785647 | 90797372 | + | 8217 | - | - && - && - | - |
| 74481 | 600.5574526 | 201.0090902 | 610.3143888 | 224.2224098 | 1.578477976 | 7.09E-05 | 0.026393494 | Batf2 | NC_000085.7 | 6214424 | 6222506 | + | 2051 | protein_coding | basic leucine zipp | bZIP_1 |
| 16145 | 7217.433312 | 2006.226996 | 7176.188726 | 1889.010175 | 1.846924205 | 7.19E-05 | 0.026393494 | Igtp | NC_000077.7 | 58090382 | 58098418 | + | 2023 | protein_coding | interferon gamma | IIGP |
| 17199 | 37.15399362 | 8.967820507 | 35.36774725 | 6.723576266 | 2.039664904 | 7.47E-05 | 0.026663204 | Mc1r | NC_000074.7 | 124133821 | 124137483 | + | 3663 | protein_coding | melanocortin 1 re | - |
| 16362 | 5076.980133 | 2405.616389 | 4952.225694 | 2485.233574 | 1.077493569 | 8.39E-05 | 0.028759879 | Irf1 | NC_000077.7 | 53660841 | 53669200 | + | 3359 | protein_coding | interferon regulat | IRF |
| 404711 | 43.79005337 | 4.098212702 | 36.6443329 | 0.091556275 | 3.369943446 | 0.000104835 | 0.035022907 | Ighg2c | NC_000078.7 | 113251009 | 113252552 | - | 1007 | C_region | - && P01864.1 F | C1-set |
| 229900 | 5652.709958 | 1884.877443 | 6053.248591 | 1753.804549 | 1.584394763 | 0.000123041 | 0.037775241 | Gbp7 | NC_000069.7 | 142236097 | 142255912 | + | 5622 | protein_coding | guanylate binding | - |
| 60440 | 8879.558991 | 2565.792339 | 10696.59408 | 2586.827826 | 1.791039116 | 0.000123294 | 0.037775241 | Iigp1 | NC_000084.7 | 60506688 | 60525706 | + | 3481 | protein_coding | interferon inducib | - |
| 20706 | 745.5772357 | 208.7054676 | 930.2090587 | 163.1029065 | 1.836060607 | 0.000123724 | 0.037775241 | Serpinb9b | NC_000079.7 | 33211397 | 33224571 | + | 1959 | protein_coding | serine (or cystein | Serpin |
| 22375 | 3790.607707 | 1588.632407 | 3499.754154 | 1619.615841 | 1.2545646 | 0.000124671 | 0.037775241 | Wars | NC_000078.7 | 108825956 | 108860100 | - | 2883 | protein_coding | tryptophanyl-tRN | - |
| 18053 | 2040.665718 | 1019.78791 | 1694.833178 | 922.6545872 | 1.000729831 | 0.000141428 | 0.041219514 | Ngfr | NC_000077.7 | 95459644 | 95478524 | - | 3409 | protein_coding | nerve growth fact | - |
| 21356 | 6656.505565 | 3306.362782 | 6485.820746 | 2989.084295 | 1.00951698 | 0.000147777 | 0.042316286 | Tapbp | NC_000083.7 | 34138452 | 34148264 | + | 2750 | protein_coding | TAP binding prot | C1-set |
| 229898 | 1478.421101 | 451.4173028 | 1489.925691 | 377.2761835 | 1.711119188 | 0.000166657 | 0.045956021 | Gbp5 | NC_000069.7 | 142202695 | 142228105 | + | 3021 | protein_coding | guanylate binding | - |
| 74748 | 201.4380674 | 84.80776059 | 174.1075093 | 67.87515411 | 1.247290571 | 0.000169306 | 0.045956021 | Slamf8 | NC_000067.7 | 172408944 | 172418482 | - | 2387 | protein_coding | SLAM family me | - |
| 18126 | 537.3687903 | 129.1068378 | 481.4429294 | 110.7759365 | 2.054802855 | 0.000176372 | 0.046896838 | Nos2 | NC_000077.7 | 78811613 | 78851052 | + | 4296 | protein_coding | nitric oxide synth | NAD_binding_1 |

**Table S7. Differential metabolites between AR and LR patients' baseline samples.**

| feature | metadata | value | coef | pval | qval | mean_LR | mean_AR | median_LR | median_AR |
|---|---|---|---|---|---|---|---|---|---|
| Glycoursodeoxycholic ac | Group | LR | 1.499990246 | 0.021197 | 0.165318993 | -9.577712897 | -11.1552287 | -9.124217925 | -10.934592 |
| Salicyluric acid | Group | LR | 0.756319896 | 0.018188041 | 0.165318993 | -11.00874353 | -11.86664184 | -11.15983845 | -11.97255095 |
| Hydrocinnamic acid | Group | LR | -1.155468288 | 0.044511484 | 0.212845826 | -10.72891498 | -9.768066638 | -10.92720042 | -9.555914114 |
| 2-Hexadecenal | Group | LR | -0.094814844 | 0.051051218 | 0.224889008 | -9.570638032 | -9.470470011 | -9.552853385 | -9.506114581 |
| Biliverdin | Group | LR | -0.419237548 | 0.053580602 | 0.226515491 | -10.55158467 | -10.16875181 | -10.5139897 | -10.08432497 |
